# Conformational Basis of Functionally Selective Allosteric Modulation of the Angiotensin II type 1 Receptor by Small Molecules

**DOI:** 10.64898/2026.07.15.736838

**Authors:** Samuel Liu, Peng Xiao, Matthias Elgeti, Eve J. Fine, Emilio Y. Lucero, Mikkel Vestergaard, Junyan Wang, Arun Jyothidasan, Angus Li, Changxiu Qu, Eva Olsen, Georgios Mazis, Josephine K. Madsen, Carl-Mikael Suomivuori, Jihee Kim, Natalia Pakharukova, Rashad Rahman, Stephanie M. Kereliuk, Walter J. Koch, Ryan T. Strachan, Dean P. Staus, Ali Masoudi, Wayne L. Hubbell, Alem W. Kahsai, Ron O. Dror, Howard A. Rockman, Jin-Peng Sun, Seungkirl Ahn, Robert J. Lefkowitz

## Abstract

Blockade of signaling through the angiotensin II type 1 receptor (AT1R), a prototypical G protein-coupled receptor (GPCR), by angiotensin receptor blockers (ARBs) is a major therapeutic approach to treating a wide variety of cardiovascular and renal diseases^1^. Like most GPCRs, the AT1R signals through two transducers, G proteins and β-arrestins^2,3^. Previous reports have described β-arrestin-biased *peptide orthosteric* agonists for the AT1R with potential therapeutic advantages over currently available unbiased ARBs^4–6^. Here we report the DNA-encoded library screening-guided isolation and pharmacological characterization of the first *small molecule AT1R allosteric* ligands. We use cryo-electron microscopy, double electron-electron resonance spectroscopy, molecular dynamics simulations, and targeted mutagenesis to determine their binding sites, binding modes and conformational mechanisms driving their unique and divergent modulatory effects on G protein and β-arrestin pathways. Our findings uncover new mechanisms for precisely controlling the dynamic behavior of the AT1R with implications for drug development targeting this pathophysiologically important receptor family.

## Main

G protein-coupled receptors (GPCRs) propagate signals through two transducer protein families, G proteins and β-arrestins^2,3^. Selective activation of one GPCR signaling pathway over the other is desirable in a multitude of situations where the signaling outcome of one pathway is beneficial while the other is deleterious^7^. Certain ligands, known as functionally selective or biased ligands, selectively activate or inhibit one pathway over others, enabling tighter control over specific receptor signaling. This has the potential to reduce unwanted side effects associated with unbiased drugs, offering safer, more efficient, and more targeted treatments^8^.

The Angiotensin II type 1 receptor (AT1R) is a prototypical GPCR and major drug target important for cardiovascular and renal homeostasis^1^. Activation of the heterotrimeric G_q_ family of G proteins by the unbiased endogenous agonist Angiotensin-II (AngII) has pathologically deleterious impacts on patients with cardiovascular disease^9^. Conversely, β-arrestin-mediated pathways have cardio-protective effects, such as increased cardiac contractility and increased cell survival signaling^4–6^. Unfortunately, existing β-arrestin-biased *orthosteric peptide* agonists which exclusively activate AT1R-mediated β-arrestin pathways^10,11^ are rapidly degraded within the body presumably due to their nature as AT1R peptide ligands, limiting their therapeutic application^12^.

To date, the overwhelming majority of GPCR drugs target the orthosteric site where endogenous ligands bind. Recently however, functionally active allosteric ligands which bind to topographically distinct sites across the extracellular, transmembrane, and intracellular surface of the receptors have been discovered^13,14^. Such allosteric ligands often possess pharmacological properties not achievable with orthosteric ligands and hold great therapeutic promise^13–16^. Of particular interest has been the ability of allosteric ligands to bias receptor signaling, either by displaying intrinsic selective activity towards either G-protein or β-arrestin-mediated signaling (i.e., biased allosteric agonism) or by converting the signaling of an unbiased orthosteric ligand into that of a biased agent (i.e., biased allosteric modulation)^17,18^. Thus AT1R-specific allosteric ligands that favor “bias” towards β-arrestin signaling potentially hold great therapeutic benefits for patients with cardiovascular disease that currently marketed drugs do not possess.

To date, very few small molecule allosteric drugs targeting known GPCR allosteric sites have been approved by the U.S. Food and Drug Administration (FDA)^14,19,20^. Strategies incorporating high throughput functional assays have been established to identify and develop GPCR allosteric modulators^21^. However, such high throughput screens are limited in size and diversity, costly, prone to false positives, and can require long timelines. Furthermore, structure-guided approaches are hampered by the limited structural information on the varied allosteric ligand binding sites and their often non-canonical modulatory mechanisms^13,22^. Nonetheless, GPCRs are allosteric machines that adopt ligand-stabilized conformational states to transduce extracellular signals to intracellular signaling cascades^23^, providing distinct conformational states and binding sites that can be utilized to enrich for new chemical matter with desired properties. These considerations prompted us to leverage our previous conformation-guided DEL experience^16,24,25^ to isolate new allosteric molecules from ultra-high throughput DEL libraries and delineate the structural mechanisms for their allosteric modulation.

### Isolation of biased allosteric modulator candidates for the AT1R by conformation-guided DEL screening

To obtain AMs for the AT1R, we conducted high-throughput screening using a DEL approach recently developed for isolating allosteric GPCR ligands^16,24,25^. In this affinity-based screening method, immobilized receptors are used to pull down small molecule binders from vast libraries of organic compounds, allowing for the identification of binders to a variety of receptor sites. Each compound is tagged with a unique DNA barcode, facilitating straightforward post-screening identification. We screened DELs comprising over 670 million unique molecules against the purified AT1R, which was reconstituted into high-density lipoprotein (HDL) particles (nanodiscs) to provide a more physiological screening environment. Furthermore, a key advantage of DEL screening is the ability to tailor screening conditions to isolate molecules with specific functional profiles, leveraging the allosteric nature of GPCR function^26^. To accomplish this, the receptor orthosteric site was occupied by β-arrestin-biased agonists TRV023 and TRV026, the G protein-biased TRV055, or the endogenous unbiased Ang II (Fig. 1a). These four orthosteric peptide ligands stabilize the AT1R in distinct conformational ensembles^27^, which, we hypothesized, would enhance the capture of allosteric ligands with corresponding functional preferences.

**Fig. 1.**
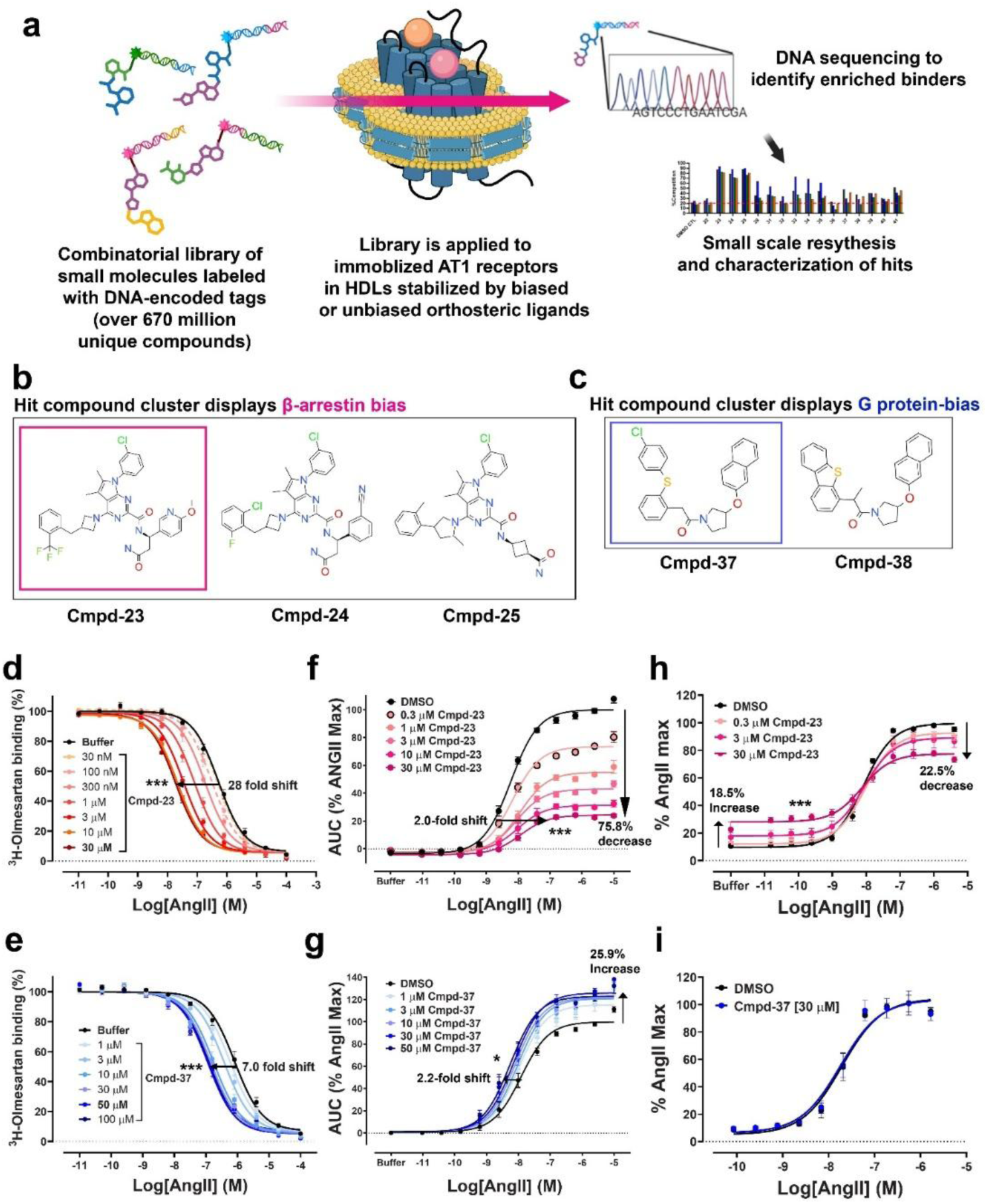
Isolation of Cmpd-23 and Cmpd-37 by conformation-guided DEL screening and their divergent pharmacological profiles. (**a**) An illustration of the DEL screening procedure, created in https://BioRender.com. (**b,c**) Chemical structures of the candidate compound cluster displaying (b) β-arrestin bias and (c) G protein bias respectively. (**d,e**) Radioligand competition binding assays. Effect of (d) Cmpd-23 and (e) Cmpd-37 on AngII binding to the AT1R. Binding of [^3^H]-Olmesartan to the AT1R against AngII in a dose-dependent manner was measured in the absence (DMSO) or the presence of (d) various concentrations of Cmpd-23 or (e) Cmpd-37 as indicated. Curve fits were obtained as described in ‘Methods’ with data sets obtained from at least three independent experiments. Each data point was normalized to the percentage of the maximum [^3^H]- Olmesartan binding level obtained from the dimethylsulfoxide (DMSO)-treated vehicle control curve and represents mean ± SEM. (**f,g**) Effects of (f) Cmpd-23 and (g) Cmpd-37 on agonist-stimulated Gq activation. Orthosteric agonist- stimulated Gq-mediated signaling in the presence of one of the allosteric compounds was monitored by the FLIPR-6 intracellular calcium mobilization assay as described in ‘Methods’. Each data point was normalized to the maximal level of the AngII-induced activity in the vehicle (0.3% DMSO) control and expressed as a percentage. The shift of curves was expressed as fold changes in EC_50_ and % alterations in Emax values. (**h,i**) Effects of (h) Cmpd-23 and (i) Cmpd-37 on agonist-stimulated β-arrestin recruitment. Orthosteric agonist-induced β-arrestin recruitment to the AT1R in the presence of the allosteric compounds was determined by the PathHunter enzyme complementation assay as described in ‘Methods’. Values were expressed as percentages over the maximum agonist-stimulated response in the vehicle-treated condition. All plotted values in the figure represent mean ± SEM obtained from at least four independent experiments.

DEL screening identifies compounds showing enriched binding to the target protein compared to empty HDL or unoccupied receptor. These enriched compounds are analyzed in the context of other binders from the entire chemical space of the initial screening library, allowing for prioritization of specific clusters of candidates for further study. Analysis of our screening output yielded 7 promising families of compounds sharing chemical scaffolds; a total of 37 putative AT1R binders were selected for resynthesis on a limited scale for pharmacological evaluation. Positive allosteric modulator (PAM) activity, which typically increases the binding affinity of an orthosteric agonist for the receptor, was assessed via orthosteric agonist competition binding against a radioligand tracer (Extended Data Fig. 1a). Initial assays identified four clusters—26A, 26B, 31B, and 31D—with significant PAM effects across all four orthosteric agonists (Extended Data Fig. 1b), amongst of which, compounds in cluster 31D and 26B displayed potentially biased modulation of AT1R in initial assays. Particularly, cluster 31D compounds differentially modulated Gαq and β-arrestin assay responses (Extended Data Fig. 1c- f), and 26B compounds showed preference for AngII and TRV055 in competition binding assays (Extended Data Fig. 1b). However, due to the extremely limited quantity of resynthesized compounds available at small scale, we were unable to determine their exact pharmacological properties at this stage.

Based on these initial results, compounds were unblinded, and representative compounds with the most favorable profiles from cluster 31D and 26B were selected for large-scale synthesis and further characterization (Fig. 1b,c). The primary goal at this step was to identify putative allosteric modulators with biased modulation of AT1R signaling. Thus, we focused on the profiling of compound (Cmpd)-23 and Cmpd-37, given their preferential modulation of AT1R signaling towards β-arrestin- and G protein-pathways, respectively.

### Divergent pharmacological profiles of Cmpd-23 and Cmpd-37

Through radioligand competition binding assays, we first examined the cooperative effects of Cmpd-23 and Cmpd-37 on the binding affinity of the endogenous orthosteric agonist AngII. Both compounds enhanced AngII binding to the receptor as evidenced by leftward shifts in AngII competition dose-response curves, with Cmpd-23 exhibiting substantially stronger positive cooperativity compared to Cmpd-37 (Fig. 1d,e). Next, we assessed the degree of cooperativity with four different orthosteric peptide agonists, which can vary due to allosteric probe dependence^15^. Interestingly, Cmpd-23 displayed modestly stronger cooperativity with the β-arrestin-biased orthosteric agonist TRV023 when compared to the other peptide agonists (Extended Data Fig. 2a). By contrast, Cmpd-37 displayed cooperative effects with the G protein-biased TRV055 and the unbiased AngII, but not significantly with the β-arrestin-biased TRV023 and TRV026 (Extended Data Fig. 2b). The specificity of Cmpd-23 and Cmpd-37 PAM activity for the AT1R over the AT2R was also determined by [^3^H]-AngII binding in the presence of these allosteric compounds. Both Cmpd-23 and Cmpd-37 increased [^3^H]-AngII binding to the AT1R without inducing detectable increases in [^3^H]-AngII binding to the AT2R (Extended Data Fig. 2c).

We next evaluated the signaling bias of Cmpd-23 and Cmpd-37 through measuring their dose-dependent effects on agonist-stimulated, Gq protein-mediated intracellular calcium signaling and β-arrestin recruitment to the AT1R. Despite the strong PAM activity of Cmpd-23 observed in binding assays (Fig. 1d and Extended Data Fig. 2a), it surprisingly inhibited AngII-stimulated intracellular calcium responses in U2OS cells, stably transfected with an inducible human AT1R expression plasmid construct^28^. These cells were utilized in the absence of an inducing reagent to minimize receptor expression^28^. Cmpd-23 markedly reduced the maximal response with modest rightward shifts of the AngII dose-response curve, consistent with strong negative allosteric modulation of efficacy (Fig. 1f). In contrast, Cmpd-37 potentiated calcium signaling, shifting the AngII dose-response curve approximately two-fold to the left and increasing the maximal response by ∼26% (Fig. 1g). Interestingly, Cmpd-37 exhibits stronger positive allosteric activity with a gain-of-function G protein-biased agonist TRV055 (Extended Data Fig. 2d) than with an unbiased AngII. Additionally, Cmpd-37 enabled the β-arrestin-biased agonist TRV023 to induce a substantial calcium response—approximately 40% of the maximal response to AngII—in cells with a high level of AT1R overexpression (Extended Data Fig. 2e). We also confirmed that both Cmpd-23 and Cmpd-37 have absolutely no intrinsic agonism for stimulating calcium responses on their own in the absence of an orthosteric agonist (Extended Data Fig. 2f). Additionally, the specificity of Cmpd-23’s inhibitory and Cmpd-37’s potentiating effects on AT1R activation was validated by their minimal effect on calcium responses stimulated by carbachol, a muscarinic receptor agonist and histamine, a histamine receptor agonist in U2OS cells (Extended Data Fig. 2g-j).

Despite its inhibitory effects on AngII-stimulated Gq activation shown in Fig. 1f, Cmpd- 23 exhibited relatively modest but significant agonistic activity in β-arrestin recruitment to the AT1R in the absence of orthosteric agonists (Fig. 1h and Extended Data Fig. 2k), which was completely abolished by an orthosteric inverse agonist (Extended Data Fig. 2l). Interestingly, Cmpd-23 also reduced the maximal response to AngII (Fig. 1h), but not significantly to TRV023 (Extended Data Fig. 2k), indicating a complex pharmacological profile with a potentially more intricate conformational basis of allosteric modulation. Cmpd-23 also exhibited PAM activity by enhancing β-arrestin endosomal targeting in response to TRV023 stimulation (Extended Data Fig. 2m). In comparison, Cmpd-37 had no effect on AngII-induced β-arrestin recruitment to the AT1R (Fig. 1i), unlike its positive effect on Gq-mediated calcium signaling (Fig. 1g). Cmpd-37 slightly potentiated TRV055-induced β-arrestin recruitment to the AT1R (Extended Data Fig. 2n), though much less than its potentiation of the TRV055-stimulated calcium response (Extended Data Fig. 2d).

To more precisely characterize the complex modulatory effects of our compounds, we quantified their cooperative effects on orthosteric ligand affinity (α) and efficacy (β) in ligand-binding and functional assays (Fig.1d-i) by the extended allosteric ternary complex model^29^ and combined operational model of allostery^30–32^, respectively. Fitting our results to these models revealed that Cmpd-23 exerts substantial positive cooperativity on AngII affinity across binding (α=33), calcium mobilization (α=24), and β-arrestin recruitment (α=18) assays (Extended Data Table 1). As expected from the curve shift behavior in Fig. 1f and 1h, Cmpd-23 possesses substantially stronger negative cooperativity toward AngII efficacy in calcium mobilization (β=0.006) than β-arrestin recruitment (β=0.06) (Extended Data Table 1). Furthermore, Cmpd-23 displays weak allosteric partial agonism for β-arrestin signaling (τ_B_=0.43) relative to AngII (τ_A_=36). Taken together, we determined that Cmpd-23 is a classical example of what is known as a PAM-antagonist^33^. Thus, functionally, it strongly antagonizes G protein activation and signaling. But simultaneously it is positively cooperative with orthosteric agonist binding as shown by radioligand binding. Cmpd-23 is also partially biased since it powerfully blocks G protein signaling compared to its relatively more permissive effect on β -arrestin activity. It also displays weak allosteric agonism (ago-PAM) towards β-arrestin recruitment. As a result, Cmpd- 23 converts AngII from an unbiased agonist to a partially β-arrestin biased agonist. On the other hand, pharmacological quantification confirms that Cmpd-37 is a G protein-biased PAM that selectively enhances Gq signaling in the presence of AngII, while leaving β-arrestin signaling nearly unchanged (Extended Data Table 1).

### Allosteric activity of Cmpd-23 in cardiomyocytes

ARB-mediated blockade of intracellular calcium signaling reduces AngII-stimulated vasoconstriction in the heart and peripheral vasculature, offering a major therapeutic approach for hypertension and heart failure^9^. In U2OS cells exogenously expressing the AT1R^28^, we found that Cmpd-23 markedly inhibits AngII-stimulated calcium responses (Fig. 1f). To further evaluate the inhibitory activity of Cmpd-23 in physiologically relevant cells endogenously expressing the AT1R, we utilized isolated cardiomyocytes from neonatal rats. In these cells, Cmpd-23 blocked the AngII-stimulated calcium response as robustly as the control antagonist losartan (Fig. 2a,b).

**Fig. 2.**
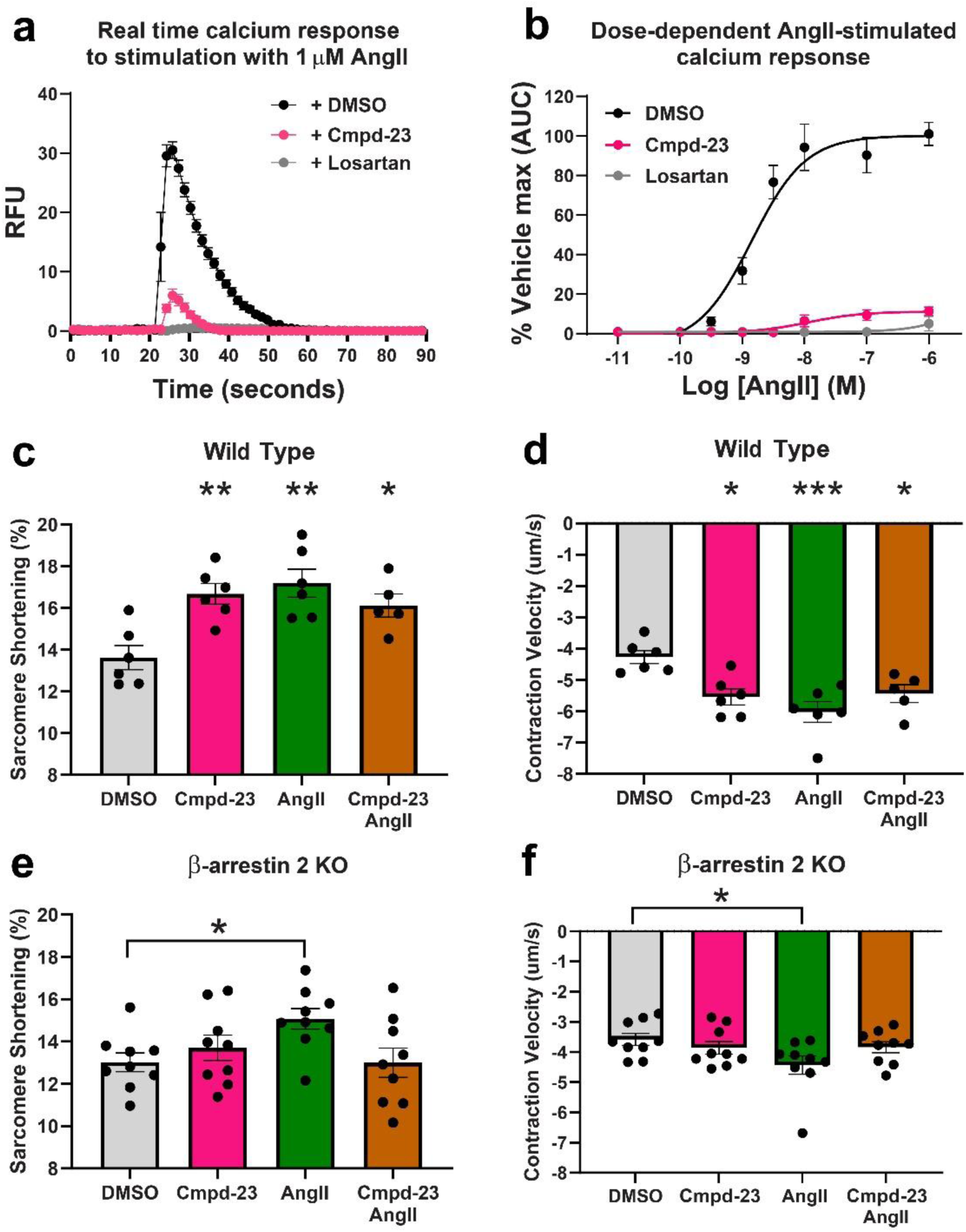
Allosteric activity of Cmpd-23 in cardiomyocytes. (**a,b**) The inhibitory effect of Cmpd-23 at 30 μM on AngII-stimulated intracellular calcium response in isolated neonatal rat cardiomyocytes, monitored as described in ‘Methods’. (a) A representative tracing of the real time changes in the intracellular calcium levels monitored after stimulation with AngII at 1 μM, the highest tested concentration. (b) Areas under the curve (AUC) were calculated at different concentrations of AngII to obtain agonist dose-response curves. Each data point was obtained from four independent experiments and normalized to the percentage of the maximal level of the AngII-induced activity in the vehicle (0.3% DMSO) control. Losartan was used as a control to show blockade of the signal by an orthosteric antagonist. (**c-f**) Contractility of adult cardiomyocyte isolated from hearts of wild type mice (c,d) and β-arrestin2 KO (e,f) was measured as (c,e) sarcomere shortening and (d,f) contraction velocity after stimulation with either AngII at 10 μM or Cmpd-23 at 25 μM alone or together. Values shown are expressed as percentage of sarcomere shortening and μm/sec velocity of shortening, respectively from 5-6 wild-type hearts and 9 β-arrestin2 KO hearts with 8-12 cells analyzed per condition per heart. Data expressed as the mean ± SEM. *p<0.05, **p<0.01, ***p<0.001 vs. DMSO, one-way ANOVA followed by Dunnett’s post-hoc test.

Previous studies have demonstrated that AT1R activation of the β-arrestin pathway exerts cardioprotective effects, including enhanced cardiomyocyte contractility and promotion of cell survival signaling^4–6^. However, existing ARBs do not differentiate between G protein and β- arrestin pathways and thus inhibit these beneficial arrestin-mediated effects. As a PAM- antagonist that is relatively permissive towards β-arrestin, Cmpd-23 would inhibit AngII-induced Gq signaling while allowing β-arrestin-mediated signaling to continue largely unimpeded. Furthermore, the partial β-arrestin-specific agonistic activity of Cmpd-23 can induce β-arrestin-mediated effects by itself. Consistent with this complex pharmacology, Cmpd-23 increased the contractility of isolated adult mouse cardiomyocytes, as measured by enhanced sarcomere shortening (Fig. 2c) and increased contraction velocity (Fig. 2d). The effects of Cmpd-23 on cardiomyocyte contractility were comparable to those observed following stimulation with AngII. Notably, co-treatment with AngII and Cmpd-23 did not result in a further increase in contractility, possibly due to Cmpd-23’s ability to restrict AngII signaling at the AT1R to the β- arrestin pathway as well as Cmpd-23’s modest negative cooperativity with AngII in the β- arrestin pathway at the high AngII concentrations used in cardiomyocytes. We further confirmed that Cmpd-23-mediated increases in cardiomyocyte contractility were through the AT1R by showing complete blockade of this positive inotropic effect of Cmpd-23 in the presence of the orthosteric inverse agonist olmesartan, as for the case of AngII (Extended Data Fig. 2o and Extended Data Fig. 3).

Importantly, we observed the contractile response induced by Cmpd-23 was completely abolished in cardiomyocytes isolated from β-arrestin2 knockout (KO) mice (Fig. 2e,f), establishing that the enhancement of contractility by Cmpd-23 is strictly β-arrestin2-dependent. In contrast, the AngII-stimulated response, mediated through both G protein and β-arrestin2 signaling, was only partially attenuated in these β-arrestin2 KO cardiomyocytes. In these β- arrestin2 KO cells cotreated with AngII and Cmpd-23 however, we observed complete elimination of the AngII-enhanced contractility, consistent with the ability of Cmpd-23 to completely bias the AngII-stimulated response towards β-arrestin. Taken together, these results demonstrate that Cmpd-23 blocks AngII-stimulated intracellular calcium signaling similarly to classical ARBs while simultaneously enhancing cardiomyocyte contractility through β-arrestin2, an effect not achievable with currently available ARBs.

### Structural characterization of the AT1R bound to Cmpd-23 and Cmpd-37

To date, a limited number of structures of biased allosteric ligands bound to GPCRs have been solved,^18,34–39^ hindering mechanistic understanding of biased allosteric modulation of GPCRs. Accordingly, we determined the binding sites of Cmpd-23 and Cmpd-37 on the AT1R utilizing cryo-electron microscopy (cryo-EM) (Fig. 3a,b). To determine the binding mode of Cmpd-23 to the AT1R, we formed protein complexes consisting of the human wild-type AT1R, Cmpd-23, TRV023, and the previously described active state stabilizing nanobody AT110i1^40^ (Fig. 3a). AT110i1 was used to stabilize the Cmpd-23 bound AT1R since it stabilizes an active conformation recognized by the β-arrestin biased ligand TRV023^41^. To improve resolvability, a “legobody” construct, composed of an MBP-fusion protein and nanobody-specific Fab^42^, which recognizes nanobody AT110i1 was added (Fig. 3a). The final protein components were separately purified and pooled together in the presence of TRV023 and Cmpd-23 to form protein complexes for cryo-EM (Extended Data Fig. 4a,b). The structure of the AT1R-Cmpd-23- TRV023-AT110i1-legobody complex was successfully determined using cryo-EM to a final nominal resolution of 3 Å (Fig. 3a and Extended Data Fig. 4c, Extended Data Table 2). The cryo-EM density map displays well resolved density, allowing us to model most portions of the AT1R and AT110i1 nanobody complex including all seven AT1R transmembrane (TM) helices and partial density for helix-8 (Fig. 3a and Extended Data Fig. 5a). Density for Cmpd-23 and the orthosteric ligand TRV023 were also well resolved. Cmpd-23 was identified to bind between TM-6, TM-7, and helix-8 at the interface between the plasma membrane and the cytoplasm (Fig. 3a,c). Cmpd-23 displays a mixture of polar, hydrophobic and van der Waals interactions with the AT1R (Fig. 3c,d). The trifluoromethylbenzyl group (R1; Fig. 3c,d) of Cmpd-23 forms the deepest point of insertion into a hydrophobic pocket created by I245^6.40^ F248^6.43^ F301^7.52^ and Y302^7.53^. Below, the pyrrolo[2,3-d]pyrimidine group (R2; Fig. 3c,d) of Cmpd-23 engages in hydrophobic and π- π interactions with F304^7.55^. Finally, towards the cytosol, Cmpd-23 forms 3 hydrogen bonds with the AT1R, between the backbone of F304^7.55^ and L305^7.56^ respectively, and one with the sidechain of K310^8.51^ (red colored in Fig. 3c,d). The two groups at the bottom of Cmpd-23, a methoxypyridine and amide group (R3; Fig. 3c,d), fit into a cytoplasm facing pocket delineated by I241^6.36^ on TM6, L305^7.56^ on TM7 and K310^8.51^ on helix-8.

**Fig. 3.**
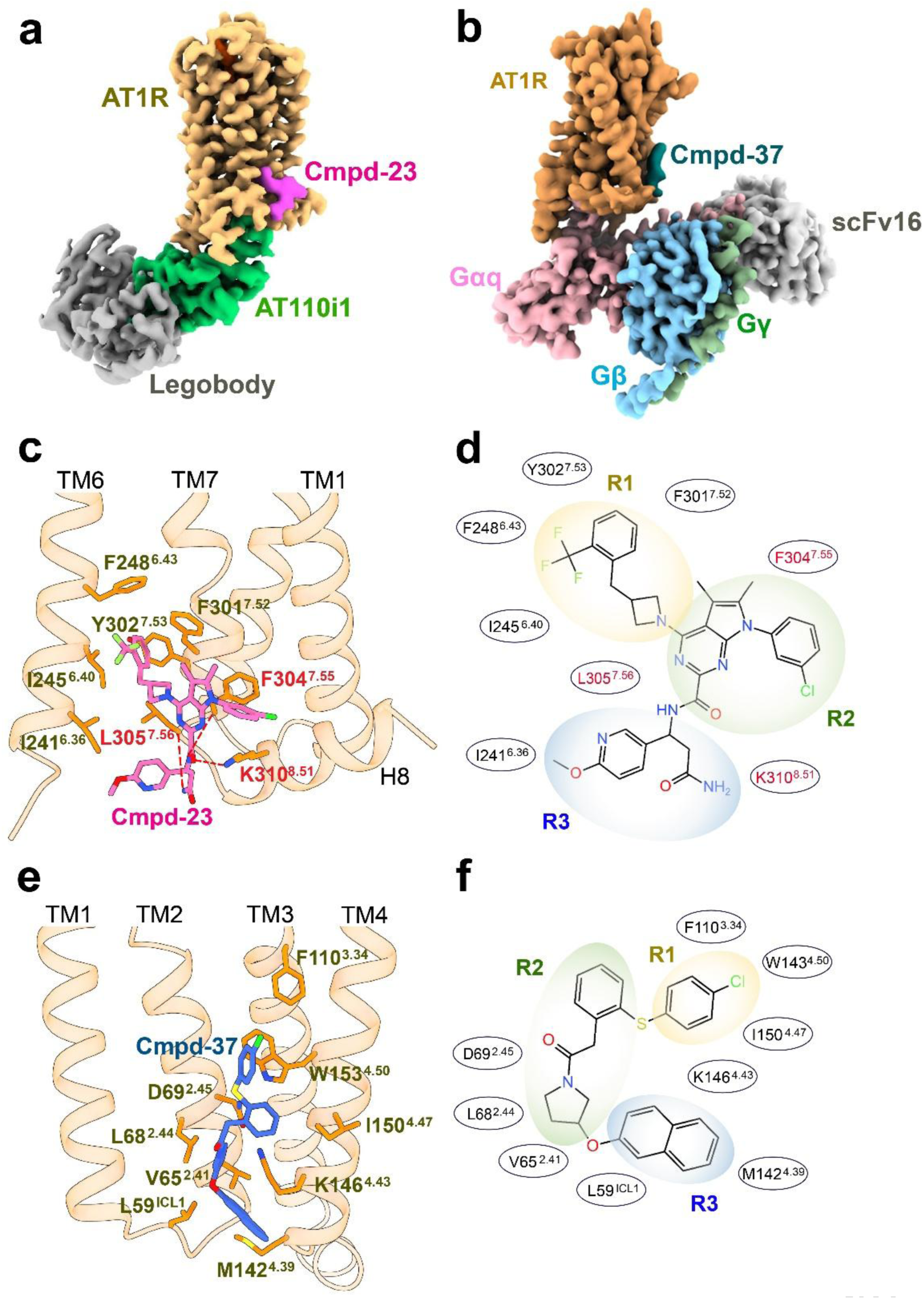
Structural characterization of the AT1R bound to Cmpd-23 and Cmpd-37. (**a,b**) Cryo-EM map of (a) AT1R-Cmpd-23-TRV023-AT110i1-Legobody complex and (b) AT1R-Cmpd37-AngII-Gαiq-scFv16 complex colored by chain. Each indicated complex component is represented by a different color together with Cmpd-23 and Cmpd-37. The contour level for each map is (a) 0.17 and (b) 3.0. (**c**) Detailed depiction of the Cmpd-23 binding site located between TM6, TM7 and H8. AT1R residues interacting with Cmpd-23 are indicated. Hydrogen bonds are displayed as dashed lines. (d) 2D ligand interaction diagram detailing the Cmpd-23 scaffold broken up into 3 regions (R1, R2, R3), with distinct interactions with the labeled residues of the AT1R. Residues highlighted in red participate in hydrogen bonding with Cmpd-23. (e) Detailed depiction of the Cmpd-37 binding pocket which is encompassed by TM1-TM4. AT1R residues interacting with Cmpd-37 are indicated. (f) 2D ligand interaction diagram detailing the Cmpd-37 scaffold broken up into 3 regions (R1, R2, R3), with distinct interactions with the AT1R labeled.

To determine the binding mode of Cmpd-37 to the AT1R, we formed protein complexes consisting of the human AT1R, together with a chimeric Gαi-Gαq protein. This engineered chimeric G protein is composed of a Gαq α5-helix fused to the body of Gαi with an additional N-terminal fusion to scFv16 used to stabilize the AT1R-Giq complex (Fig. 3b). We co-expressed this construct together with the human WT AT1R, Gβ1, and Gγ2 in Sf9 insect cells, which were then purified in the presence of AngII and Cmpd-37 to obtain a stable AT1R-Giq complex (Extended Data Fig. 6a,b). The structure of AngII-AT1R-Giq in complex with Cmpd-37 was determined at a global resolution of 3.2 Å (Fig. 3b and Extended Data Fig. 6c, Extended Data Table 2). The cryo-EM maps allowed us to model most portions of the AT1R, the Giq heterotrimer, Cmpd-37, and scFv16 (Fig. 3b and Extended Data Fig. 5b). Besides cryo-EM density for AngII observed in the orthosteric binding pocket of the AT1R, additional density attributed to Cmpd-37 was detected on the receptor’s cytoplasmic surface in a pocket formed by ICL1, TM2, TM3, and TM4 (Fig. 3b,e). Most of the interactions between Cmpd-37 and AT1R are driven by hydrophobic and van der Waals forces, reflecting the hydrophobic nature of the compound (Fig. 3e,f). At the top of the Cmpd-37 binding pocket, the 4-chlorophenyl group (R1; Fig. 3e,f) forms hydrophobic or π-π interactions with F110^3.34^ and/or W153^4.50^. The core scaffold of Cmpd-37 (R2; Fig. 3e,f) lies on a hydrophobic surface created by D69^2.45^, F110^3.34^, and W153^4.50^, forming both hydrophobic interactions and van der Waals contacts. The naphthalene group (R3; Fig. 3e,f) of Cmpd-37 fits into the cytoplasmic surface pocket created by L59^ICL1^, V65^2.41^, M142^4.39^, and K146^4.43^ facilitating not only hydrophobic interactions, but also a potential cation-π interaction involving K146^4.43^.

### Conformational basis for β-arrestin biased regulation of AT1R signaling by Cmpd-23

Crystallographic and cryo-EM structural approaches often capture only a single, low-energy receptor conformation, which can be substantially influenced by fiducial markers and stabilizing proteins (i.e., nanobody or G-protein). This may obscure how distinct receptor conformations contribute to the action of the bound ligand. Particularly, a critical question in GPCR allostery is whether allosteric modulators bias orthosteric ligand signaling by stabilizing existing biased conformations within the orthosteric ligand-stabilized ensemble or rather introduce new biased states. Accordingly, to better understand the mechanism of Cmpd-23, we employed double electron-electron resonance (DEER) spectroscopy, which has proven useful in elucidating conformational dynamics of GPCRs ^27,43^.

Our previous DEER study identified TM6 and ICL2 configurations as key determinants of functionally distinct AT1R conformations^27^. Thus, AT1R receptors with pairwise spin labels on either TM1-TM6^27^ or TM1-ICL2^27^ were prepared (Fig. 4a and Extended Data Fig. 7a,b) and analyzed in the presence and absence of Cmpd-23 and different orthosteric ligands. First, we observed similar DEER profiles between AngII and TRV023 in the absence of Cmpd-23 (Fig. 4b,c) consistent with our previous findings^27^ showing that both orthosteric ligands stabilize a conformational ensemble featuring a substantial population of the “occluded-2” state. This conformation is characterized by a relatively broad TM1–TM6 distance (>40 Å, Fig. 4b) and a characteristic short TM1–ICL2 distance (26 Å, Fig. 4c), which is known to be connected to β- arrestin bias^27^. Then, in the presence of Cmpd-23, the receptor ensemble shifts further toward the occluded-2 conformation regardless of the bound orthosteric ligand (Fig. 4b,c and Extended Data Fig. 7c-f).

**Fig. 4.**
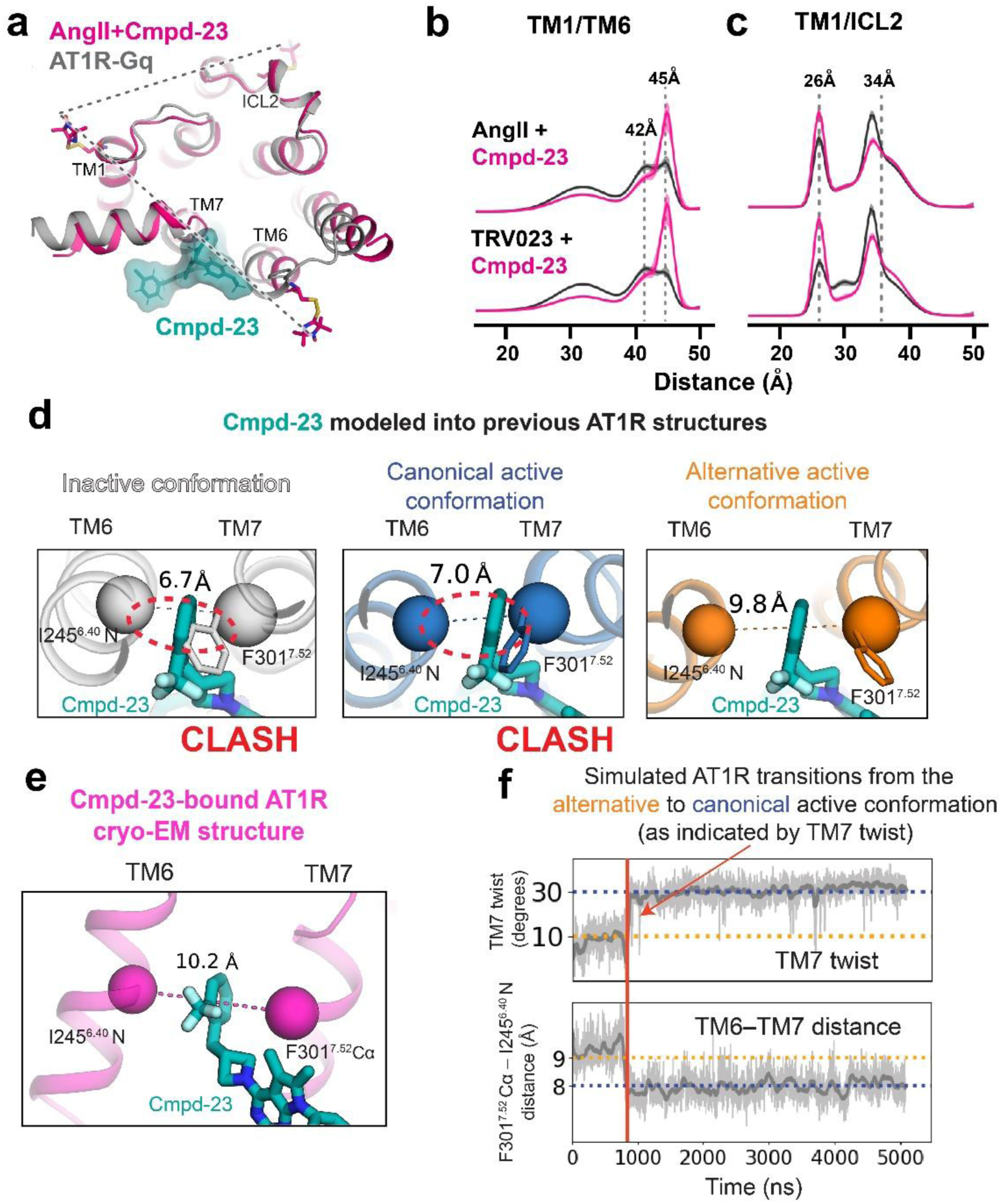
Conformational basis for β-arrestin biased regulation of AT1R signaling by Cmpd-23. (**a**) Comparison of AT1R structures with and without Cmpd-23 and indication of spin label positions. Cmpd-23 forms a wedge between TM6 and TM7. (**b,c**) Effect of Cmpd-23 (b) on TM1/TM6 distance and (c) on TM1/ICL2 distance in the presence of AngII and TRV023. (**d**) Top-down views of Cmpd-23 (cyan) overlaid on representative structures of AT1R without Cmpd-23 bound, in the inactive, canonical active (blue), and the β-arrestin biased alternative active (orange) conformations (PDB entries 8TH4, 7F6G, and 6OS1, respectively). In each panel, the F301^7.52^ Cα – I245^6.40^ N measurement is shown via a dashed line with a nearby label. Only the alternative active conformation (right) has sufficient TM6–TM7 separation for Cmpd-23 to bind without clashing against TM6 or TM7. (e) The Cmpd-23-bound cryo-EM structure, with the F301^7.52^ Cα – I245^6.40^ N distance shown. (f) Time traces showing the twist angle of TM7 (top, see Methods) and the TM6–TM7 separation (bottom, F301^7.52^ Cα – I245^6.40^ N distance) in the same simulation of AT1R without a nanobody bound. The blue and orange dashed lines show approximate values for the canonical and alternative active conformations, respectively, for the TM7 twist and F301^7.52^ Cα – I245^6.40^ N measurements. The TM6– TM7 gap is wider in the alternative active conformation than in the canonical active conformation. Smoothed traces are shown in dark gray, with raw simulation traces in light gray.

Moreover, technical advances in DEER analysis allowed us to further distinguish between two active TM6 positions, one centered around 42 Å distance and one at 45 Å. Cmpd- 23 exclusively stabilized the longer and much narrower TM1-TM6 distance peak, stabilizing a subpopulation within the ensemble of β-arrestin-biased conformations. This specific modulation corresponds directly to a small but distinct outward shift of TM6 caused by the physical interaction between Cmpd-23 and the intracellular helical bundle. *In silico* spin labeling of the high-resolution structures obtained in this study confirmed that the Cmpd-23 bound state is consistent with the extended DEER distance at 45 Å (Extended Data Fig. 8). Therefore, Cmpd- 23 selectively stabilizes a subset of “occluded-2” state together with orthosteric agonists, thereby shifting distinct orthosteric ligand-bound receptor states toward more similar conformations. The minimal changes in the DEER profile in the presence of Cmpd-23 alone are also consistent with the weak intrinsic agonism of Cmpd-23. Note that ligand effects on the AT1R’s conformational ensemble in DEER may be underestimated in the LMNG micellar environment as previously described^27^.

To further explore the molecular mechanism by which Cmpd-23 induces conformational changes, we examined how Cmpd-23 fits into AT1R conformations observed in previous structures and molecular dynamics (MD) simulations^41,44,45^. In agreement with our DEER results, these simulations demonstrate that Cmpd-23 binding is not compatible with the previously observed inactive or canonical active AT1R conformations^41,44,45^ (Fig. 4d, left and middle panels). These conformations have too small a gap between the intracellular portions of TM6 and TM7, where the trifluoromethylbenzyl head of Cmpd-23 inserts. Interestingly, the β-arrestin-biased conformation we previously identified in our MD simulation study^44^, defined by a twist of the intracellular portion of TM7, features a substantially larger TM6–TM7 separation that accommodates Cmpd-23 binding (Fig. 4d, right panel labeled as “alternative”). This larger TM6– TM7 separation is observed in in the Cmpd-23-AT1R-AT110i1 complex reported here (Fig. 4e). To verify that the larger TM6–TM7 separation is not simply an artifact of AT110i1, we examined MD simulations of AT1R without either AT110i1 or Cmpd-23 present. In these simulations, the larger TM6–TM7 separation is consistently observed in this β-arrestin-biased active conformation but not in the canonical active conformation (Fig. 4f and Extended Data Fig. 9). Thus as investigated by both DEER spectroscopy and MD simulations, Cmpd-23 appears to exert its effect via a similar mechanism to that of β-arrestin-biased orthosteric agonists despite binding in a different site: Cmpd-23 provokes a β-arrestin-biased conformation similar to that stabilized by β-arrestin-biased orthosteric ligands, stimulating β-arrestin signaling while blocking Gq signaling, consistent with our pharmacological data (Fig. 1f,h).

An additional regulatory feature of Cmpd-23 binding is its terminal amide group in the R3 region (Fig. 3c,d) that reaches toward the intracellular transducer binding interface (Extended Data Fig. 10). Accordingly, Cmpd-23 binding could directly hinder transducer coupling, representing a second potential mechanism, by which Cmpd-23 could exert its biased modulation. Modeling of Gq or β-arrestin in complex with the AT1R in the Cmpd-23-bound structure suggests that the amide group of Cmpd-23, which binds at the bottom of the AT1R, could destabilize Gq coupling but not β-arrestin coupling to the receptor core (Extended Data Fig. 10). One would also expect Cmpd-23 binding to disrupt G protein coupling more than β- arrestin coupling because β-arrestins have been found to couple to GPCRs in more varied conformations than G proteins^46–48^. As such, the protrusion of Cmpd-23 into the receptor core interface may inhibit G protein coupling more than β-arrestin coupling. At the same time, similar to a transducer, Cmpd-23 binds near the transducer binding site and stabilizes an active-like conformation with increased affinity for AngII, as observed in Fig. 1d. This is consistent with our DEER results where Cmpd-23 stabilizes an active-like conformation with increased TM1- TM6 distance.

### Conformational basis for Gαq biased regulation of AT1R signaling by Cmpd-37

We also investigated the effect of Cmpd-37 binding on the AT1R conformational ensemble using the TM1-TM6 and TM1-ICL2 DEER probes (Fig. 5a and Extended Data Fig. 7a,b). Our TM1-TM6 results show that Cmpd-37 stabilizes the active, outward-tilted TM6 position around 42 Å, while the TM1-ICL2 pair shows destabilization of the short distance at 26 Å (Fig. 5b,c and Extended Data Fig. 7c-f) in the presence of orthosteric agonists. Both conformational changes suggest that Cmpd-37 favors the canonical active, G-protein-biased “open” conformation^27^, which is also consistent with our pharmacological data showing augmentation of the G-protein signaling pathway by Cmpd-37 (Fig. 1g and Extended Data Fig. 2d). A comparison with simulated distances from *in silico* spin labeling of our cryo-EM structure further confirms our interpretation, as TM1-TM6 and TM1-ICL2 distances in our cryo-EM AT1R structure with Cmpd-37 are largely consistent with our DEER experimental data (Extended Data Fig. 8).

**Fig. 5.**
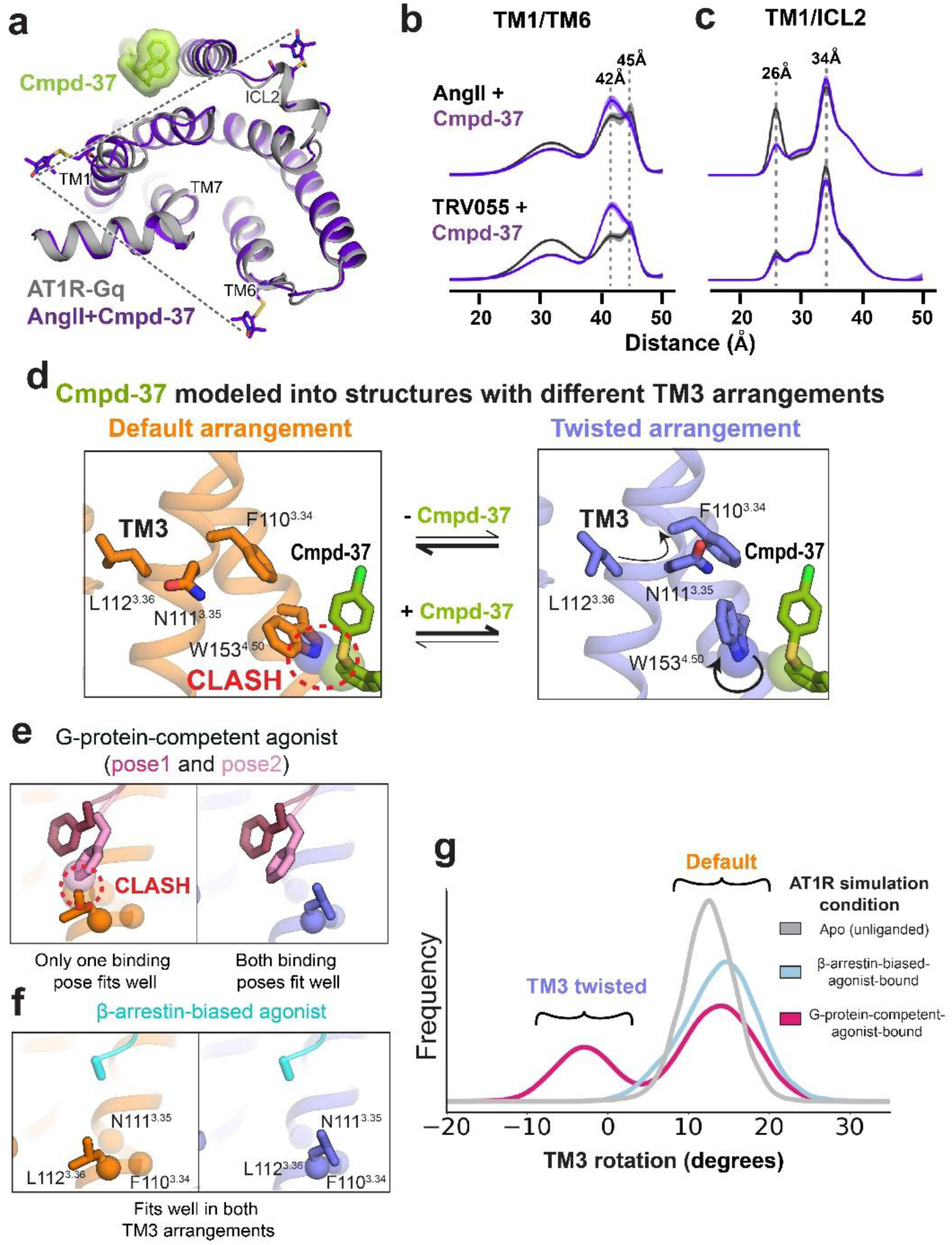
Conformational basis for Gαq biased regulation of AT1R signaling by Cmpd-37. (**a**) Comparison of AT1R structures with and without Cmpd-37 and indication of spin label positions. Cmpd-37 binds to the outer surface of TM2 and TM4. (**b,c**) Effect of Cmpd-37 (b) on TM1/TM6 distance and (c) on TM1/ICL2 distance. (**d**) Cmpd-37 (green) overlaid on representative AT1R structures in which TM3 adopts its default arrangement (left, orange and twisted arrangement (right, dark blue) at the orthosteric binding pocket (PDB entries 6OS1 and 7F6G, respectively). The arrangement of TM3 affects the rotamer of W153^4.50^, allowing Cmpd-37 to bind tightly only when TM3 is in the twisted arrangement. (**e,f**) The two binding poses of G-protein-competent agonist AngII (light pink) observed in simulation are overlaid on the same TM3 structures as in panel A. Only pose 1 (dark pink) is compatible with the default arrangement of TM3, but both poses are compatible with the twisted arrangement of TM3. β-arrestin- biased-agonist TRV023 (light blue) is equally compatible with both TM3 arrangements. TRV023 coordinates from PDB 6OS1 are overlaid on TM3 coordinates from the same TM3 structures as in panel A. (g) Frequency distributions (“histograms”) of the TM3 rotation measurement in unliganded (gray), G- protein-competent-agonist-bound (magenta), and β-arrestin-biased-agonist-bound (light blue) simulations of AT1R. Simulations were performed with G-protein-competent agonists AngII and TRV055, and with β-arrestin-biased-agonists TRV023 and TRV026. The TM3 rotation angle at the beginning of each simulation was 16°, which represents the default TM3 arrangement. The presence of a G-protein- competent agonist increases the frequency of the TM3 twisted arrangement, because G-protein-competent agonists bind preferentially to that arrangement.

An additional finding is that the conformational shifts for Cmpd-37 depend on the nature of the bound orthosteric ligand, and that the strongest stabilization of the “open” conformation is observed with the Gq-biased agonist TRV055 followed by balanced agonist Ang II. (Extended Data Fig. 7e,f). Thus, in addition to stabilizing the “open” conformation induced by orthosteric agonists, Cmpd-37’s modulatory effects are also substantially probe-dependent—varying based on the orthosteric ligand present, a phenomenon also observed with Cmpd-37’s ligand specific cooperativity in binding assays (Fig. 1e and Extended Data Fig. 2b).

To explain this probe-dependent allosteric modulation, we performed structural examination of the Cmpd-37 binding mode with the previously reported crystallographic AT1R structures^41,45^. This revealed that Cmpd-37 exerts its effects by stabilizing a particular arrangement of the orthosteric ligand binding pocket. Specifically, Cmpd-37 binds to a portion of TM3 in an allosteric site adjacent to, but outside of, the orthosteric ligand binding pocket. This site has been observed to adopt two main arrangements differing by a rotation or “twist” around the helical axis^44^. In unliganded AT1R simulations^42^, TM3 nearly always adopts one arrangement, which we refer to as the “default” (Fig. 5d). In the Cmpd-37 bound structure, however, TM3 adopts the other arrangement, which we refer to as the “twisted” arrangement. Cmpd-37 stabilizes the twisted arrangement because Cmpd-37 binding requires that W153^4.50^ adopt a particular rotamer that would clash with F110^3.34^ in the default TM3 arrangement (Fig. 5d and Extended Data Fig. 11a,b).

G-protein-competent agonists such as unbiased AngII and G protein-biased TRV055 stabilize the twisted arrangement of TM3 due to their bulky terminal phenylalanine residue, which adopts two main binding poses^44^. One pose clashes with L112^3.36^ when TM3 is in the default arrangement, but both poses can be adopted by the agonist when TM3 is in its twisted arrangement (Fig. 5e). This terminal residue in β-arrestin-biased agonists is generally smaller or absent, allowing these agonists to bind regardless of which arrangement TM3 adopts (Fig. 5f). In contrast, G-protein-competent agonists bind more tightly to and stabilize the twisted arrangement. Indeed, our simulations show that G-protein-competent agonists substantially increase the population of the twisted arrangement, whereas β-arrestin-biased agonists do not (Fig. 5g and Extended Data Fig. 11c,d).

According to our proposed structural mechanism, Cmpd-37 stabilizes the twisted arrangement of TM3, and thus it should selectively increase the affinity of G-protein-competent orthosteric agonists. Indeed, competition ligand binding data shows precisely such selective positive cooperativity: Cmpd-37 enhances G-protein-competent agonist affinity but not β- arrestin-biased agonist affinity (Fig. 1e and Extended Data Fig. 2b). Our previous MD simulations study suggested that the twisted arrangement of TM3 also favors adoption of the canonical active receptor^44^. On its own, however, the adoption of either TM3 arrangement appears insufficient to stabilize any active conformation of AT1R. Our proposed mechanism thus implies that Cmpd-37 would not stimulate Gq-mediated signaling or β-arrestin recruitment on its own, but that in the presence of an orthosteric agonist such as AngII, Cmpd-37 would increase G-protein signaling much more than β-arrestin recruitment, as observed in pharmacological experiments (Fig. 1g,i and Extended Data Fig. 2d,f,n). Taken altogether, these findings suggest that the actions of Cmpd-37 are mediated by modulating the distribution of the receptor amongst conformational ensembles previously identified with orthosteric ligands rather than by stabilizing unique ones.

### Mutagenesis validates distinct residues responsible for Cmpd-23 and Cmpd-37 binding and activity

To validate the proposed mechanisms of Cmpd-23 and Cmpd-37 binding and action, we mutated residues surrounding the Cmpd-23 and Cmpd-37 binding sites. We identified multiple mutations that should impact Cmpd-23’s and Cmpd-37’s modulatory activity with little to no effect on AngII-induced AT1R signaling.

At the Cmpd-23 binding site, we generated I241A^6.36^ and I245A^6.40^ double, F301A^7.52^, and K310A^8.51^ single mutants (Fig. 6a), all of which impaired Cmpd-23’s ability to suppress AngII-induced Gq signaling measured by the Trupath G protein dissociation assay^49^ (Fig. 6b-f). The I241A^6.36^ and I245A^6.40^ double (blue) and F301A^7.52^ (green) mutants all alter residues important for forming hydrophobic contacts with the benzotrifluoride group on Cmpd-23 (Fig. 6a), so it is also expected that altering these groups would affect Cmpd-23 activity (Fig. 6b-e). K310^8.51^ forms an important hydrogen bond with the main body of Cmpd-23 (Fig. 6a), and thus it was unsurprising that disrupting this bond led to a complete loss of function of Cmpd-23 activity (Fig. 6b,f). A surprising finding was that the F301A^7.52^ mutation appeared to reverse Cmpd-23’s inhibitory activity on Gαq signaling, resulting in slight, but significant, PAM effects (Fig. 6b,e). Further modeling of the F301A^7.52^ mutation shows that removal of the bulky phenylalanine allows a sufficient TM6–TM7 gap for Cmpd-23 binding regardless of which active conformation the AT1R adopts (Extended Data Fig. 12), leading to the reversal of Cmpd- 23’s inhibition of Gq coupling.

**Fig. 6.**
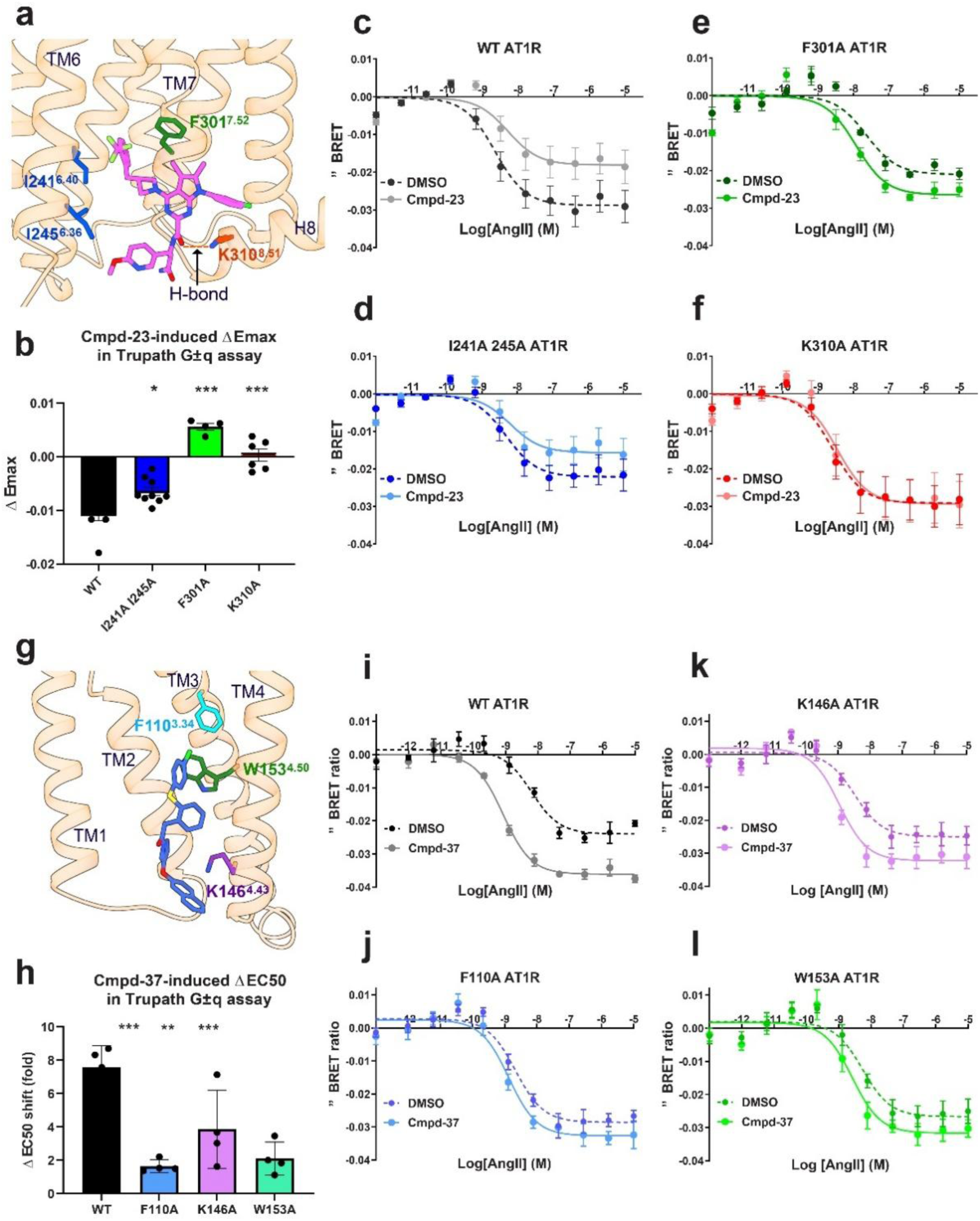
Mutagenesis validates distinct residues responsible for Cmpd-23 and Cmpd-37 activity. (**a**) AT1R residues, mutagenized to validate Cmpd-23 activity, at its binding site. (**b**) Mutant effects on Cmpd-23-induced blockade of Gq dissociation as measured by Trupath assays. Values were expressed as the extent of ΔEmax (changes of Emax between conditions in the absence and presence of Cmpd-23) and represent mean ± SEM obtained from the data sets shown in c-f. (**c-f**) Effects of Cmpd-23-mediated blockade of Gq protein activation in (c) WT or (d) I241A and I245A, (e) F301A, or (f) K310A mutant receptors via Trupath. Each data point was expressed as ΔBRET ratio (changes over the minimum value) and represent mean ± SEM obtained from four independent experiments. (g) AT1R residues, mutagenized to validate the Cmpd-37 activity, at its binding site. (h) Mutant effects on overall Cmpd-37 potentiation of Gq dissociation as measured by Trupath assay. Values are expressed as the extent of ΔEC_50_ (changes of EC_50_ between conditions in the absence and presence of Cmpd-37) and represent mean ± SEM obtained from the data sets shown in i-l. (**i-l**) Effects of Cmpd-37-mediated potentiation of Gq protein activation in (i) WT or (j) mutant F110A, (k) K146A or (l) W153A receptors via Trupath. Changes in BRET (ΔBRET) ratio were measured and expressed with the data sets obtained from four independent experiments.

At the Cmpd-37 site, we generated F110A^3.34^, K146A^4.43^ and W153A^4.50^ mutants (Fig. 6g), all of which impaired the ability of Cmpd-37 to potentiate AngII-induced Gq activation to varying extents (Fig. 6h-l). Notably, we postulated that F110^3.34^ and W153^4.50^ mediate communication with the orthosteric ligand, which could explain the near complete loss of modulation at F110A (Fig. 6h,j,l). Additionally, W153^4.50^ and K146^4.43^ are important for forming the Cmpd-37 binding pocket via π-π and π-cation interactions, respectively. Taken together, these mutagenesis results assign functional relevance to the binding site residues identified in our structural characterization of Cmpd-23 and Cmpd-37.

## DISCUSSION

In this study, we identified small molecule allosteric ligands targeting the AT1R through DEL screening and characterized their biased signaling profiles toward either G protein or β- arrestin pathways. A critical highlight of our study is the identification of new AM binding pockets and the molecular switches these AMs modulate to display their biased activities at the AT1R. Previous studies of β-arrestin biased^18,35,36^ and G-protein biased allosteric ligands^34,39^ have highlighted steric blockade and/or direct interactions with transducers as mechanisms of biased allosteric modulation. In contrast, our study underscores that both Cmpd-23 and Cmpd-37 function through different forms of conformational selection, modulating the receptor conformational ensemble to bias downstream signaling. Cmpd-23 additionally regulates biased receptor signaling through a direct steric mechanism. Future research could further delineate this dual mechanism of receptor regulation and leverage it to achieve even tighter control over transducer selectivity in compound optimization efforts at the Cmpd-23 pocket.

The extensive conformational characterization of Cmpd-23’s and Cmpd-37’s modulatory effects on the AT1R is a strength of our study. Previous biophysical studies have created rich frameworks for understanding AT1R activation within the context of known orthosteric ligands^27,40,41,44,45^. Our study highlights the complementarity of diverse approaches in understanding receptor structural dynamics. Nonetheless, a central question in AT1R molecular pharmacology efforts is whether additional, yet unidentified signaling states of the AT1R exist, and whether newly identified allosteric ligands stabilize pre-existing signaling states or novel ones. Our DEER results indicate that Cmpd-23 selects a conformational sub-ensemble of the previously identified β-arrestin-biased “occluded-2” conformation^27^. Meanwhile, DEER was also able to identify that Cmpd-37 enhanced a conformational ensemble corresponding to the canonical “open” state. Combining our cryo-EM, DEER, MD simulations, and pharmacological results reveal specific molecular mechanisms of each compound.

Our structural study also demonstrates potential applications for structure-guided approaches for biased AM discovery at the AT1R. Identification of Cmpd-23’s binding site further builds the possibility that the TM6-TM7 region may be a hotspot for allosteric regulation of GPCRs^38,39^ while showing a novel allosteric binding mode and regulatory mechanisms resulting in a unique β-arrestin-biased signaling profile. Additionally, the identified binding site of Cmpd-37 in our study overlaps with a known cholesterol binding site at the AT1R^45^. Utilization of lipid-binding pockets located in proximity to conformation-specific molecular switches could provide additional targets for the development of biased AMs for GPCRs.

Interestingly, both Cmpd-23 and Cmpd-37 display potential probe dependent behaviors such as the highest binding cooperativity of Cmpd-23 with TRV023, or the almost exclusive binding cooperativity of Cmpd-37 with AngII and TRV055. However, these behaviors were only partially characterized in this study, and future research should further investigate the specific probe-dependent pharmacological and structural consequences of AT1R allosteric modulator and orthosteric peptide agonist combinations.

Our pharmacological characterization of Cmpd-23 classifies it as a PAM-antagonist with relative signaling bias towards blockade of G-protein signaling and partial β-arrestin-biased allosteric agonism. In biologically relevant murine cardiomyocytes, the β-arrestin signaling of Cmpd-23 enhances contractility comparable to that of AngII. The absence of additive effects when co-administered with AngII and the complete loss of function in β-arrestin2 knockout cardiomyocytes support Cmpd-23’s strictly β-arrestin-dependent signaling mechanism. Additionally, these effects in cardiomyoctes are consistent with the strong blockade of AngII-stimulated, Gq-mediated effects by Cmpd-23 and its relatively modest inhibition of AngII-stimulated β-arrestin signaling. Our results suggest that allosteric ligands possessing similar pharmacological profiles to Cmpd-23 may reduce blood pressure akin to classical ARBs but with added functional benefits such as improving cardiac performance and providing cardioprotection under conditions of injury^5^. Furthermore, allosteric ligands typically offer pharmacological benefits such as robust specificity for the target receptor and a ceiling effect that reduces the risk of overdose.

In contrast, Cmpd-37 selectively enhances the binding and signaling of AngII and the G protein-biased agonist TRV055 but shows no PAM activity for the binding of β-arrestin-biased agonists TRV023 and TRV026. However, in cells overexpressing the AT1R, Cmpd-37 was able to augment the calcium response of the β arrestin-biased agonist TRV023. Even though the “open” state is only a small portion of the overall conformational ensemble stabilized by TRV023^27^, positive coupling may be driven by these low probability sub-states present in the receptor population^50^. Overall, Cmpd-37 significantly potentiates Gq-mediated intracellular calcium responses while leaving β-arrestin recruitment minimally affected, clearly indicating its bias toward G protein signaling. Augmentation of such signaling responses may have utility in promoting activation of the AT1R in conditions where the Gq-dependent activity of the AT1R is desirable (e.g., hypotensive shock^51^).

It is important to note that the existence of system bias, such as shown in the case of expression systems highly overexpressing the AT1R^28^, has been a limitation of previous studies characterizing the signaling profiles of biased orthosteric ligands at the AT1R. Encouragingly, though our study uses a range of experimental systems with multiple cellular backgrounds for the characterization of our compounds, we observed consistent signaling trends and pharmacological parameters across experimental systems, including in cell lines and cardiomyocytes at native levels of receptor expression. Further compelling was the finding of robust conservation of positive cooperativity for AngII affinity across both binding and functional assays. Nonetheless, caution should be taken in extrapolating results to other experimental systems with different degrees of receptor expression or a distinct cell line background. Care should also be taken in translating the primary murine cardiomyocyte data to animal models given the supraphysiological concentrations of AngII used in vitro. Therefore, future pharmacokinetic-pharmacodynamic studies with Cmpd-23, Cmpd-37, or more suitable analogs should be performed in cardiovascular disease models to test the therapeutic potential posited here.

In summary, this study presents novel small-molecule allosteric ligands with divergent modulation of the AT1R identified via DEL screening. Comprehensive pharmacological and structural profiling identified Cmpd-23 as a PAM-antagonist with inhibitory activity toward G protein signaling and modest partial β-arrestin-biased allosteric agonism, while Cmpd-37 functions as a G protein-biased PAM. Our combined cryo-EM, DEER spectroscopic, and MD simulations studies elucidate distinct structural underpinnings exploited by each compound. While not claiming that Cmpd-23 itself is a drug candidate, our study provides proof of principle for the idea that allosteric ligands such as Cmpd-23 or its analogs may have promising therapeutic potential for the treatment of cardiovascular disease. Collectively, our findings expand the scope of DEL-based allosteric ligand discovery and provide a mechanistic foundation for developing next-generation biased GPCR therapeutics.

## METHODS

### Chemical reagents

Expi293 expression media (A1435102), Minimal Essential Media (MEM) (11095-080), Hygromycin B (10687010), glutamine (25030-081), dimethyl sulfoxide (DMSO) anhydrous (A14525), and Expifectamine Transfection Kits (D12345) were purchased from Thermo Fisher Scientific. Blasticidin (ant-bl-1) and Zeocin (R25001) were purchased from Invitrogen. Fetal bovine serum (F2442), Doxycycline hyclate (D9891), sodium butyrate (303410), cholesterol hemisuccinate (CHS) (C6512), benzonase (C6512), and benzamidine hydrochloride hydrate (B6506) were purchased from Sigma. G418 and carbacol were purchased from Sigma-Aldrich. Penicillin-Streptomycin was purchased from Gemini (400-109). Fugene 6 transfection reagent was purchased from Promega (E2692). Commercially available ligands were purchased as follows: AT1R orthosteric peptides and FLAG peptide were custom synthesized from GenScript. Candesartan (4791) and Olmesartan (4616) were purchased from Tocris. Losartan was purchased from TCI (L0232). [3H]-olmesartan was purchased from American Radiolabeled Chemicals (ART 1976). For protein purification, a Sephadex 200 Increase 10/300 GL column was purchased from GE Healthcare (28-9909-44). Ni2+ agarose beads were purchased from Goldbio (H-350-100). For spin labeling of DEER constructs, Bis-(2,2,5,5-Tetramethyl-3-imidazoline-1-oxyl-4-yl) disulfide, biradical (IDSL) was purchased from Enzo (ALX-430-102). D8-Glycerol (DLM-558) and deuterium oxide (DLM-4-99) were purchased from Cambridge Isotopes.

### Cell culture

HEK-293, U2OS, and CHOK1 cells were cultured at 37 °C in a humidified environment (5% CO2) using standard minimum Eagle’s, Dulbecco’s modified Eagle’s, and F-12, growth media, respectively supplemented with 10% FBS and penicillin/streptomycin. HEK-293 cell lines stably expressing the rat AT1R^52^ and U2OS cells stably transfected with the tetracycline-inducible human AT1R expression plasmid^28^ were maintained as described before. CHOK1 cells, stably expressing AT1R-pk and β-arrestin2- EA for the PathHunter β-arrestin2 enzyme complementation assay were obtained from Eurofins and maintained according to the manufacturer’s guidelines. Expi293F cells were cultured in suspension using Expi293F expression medium at 37°C in a humidified incubator with 8% CO₂, agitated at 110 rpm. A specialized derivative line—Expi293F inducible—stably expressing the tetracycline repressor (pcDNA/TR) for enabling Tet-inducible protein expression were routinely maintained with 10 µg/mL blasticidin—which was withdrawn in the final passage before transfection.

### Compound synthesis

Thirty-seven putative AT1R binders were synthesized without their DNA tags from Amgen Research at Copenhagen^53^. All chemicals and solvents, unless otherwise stated, were purchased from standard suppliers—ChemDiv (San Diego CA, USA); PepTech Corporation (Bedford, MA, USA); Matrix Scientific (Columbia, SC, USA); Specs (Zoetermeer, Netherlands); Enamine BB (Kyiv, Ukraine) and were used without further purification. Structural assignment for small scale synthesis compounds was performed by HRMS only and based on fragment structures given by the suppliers. Preparative HPLC was performed on an Agilent Prep 1290 instrument with UV and MS(SQ). Analytical Column: PoroShell 120 SB C18 Column Agilent (4.6 × 150 mm, 4 μM) at a flow rate of 1.5 mL/min. Solvent A: 0.1% formic acid in aqueous and Solvent B: 0.1% formic acid in MeCN. Gradient, 95% A to 100% B over 10−30 min. MS Settings: Positive mode, UV from 190 to 400 nm, MS spectrum from 150 to 1300 m/z. Analytical data (purity at 254 nM, HRMS, and concentration) was established employing an Agilent 1290 Infinity II LC System coupled with a TOF mass spectrometer Agilent 6230B, equipped with a diode array detector and an Antek 8060 nitrogen-specific chemiluminescent detector. Column: Acquity Premier UPLC BEH C18 column (1.7 μm, 2.1 × 50 mm, Waters) at 45 °C. Solvent A: 0.1% formic acid in aqueous and Solvent B: 0.1% formic acid in MeOH. Gradient, 95% A to 100% B over 5.35 min, flow Rate: 0.75 mL/min. MS Settings: Positive mode, mass range from 100 to 3000 m/z, and UV detection from 190 to 500 nm. All compounds were sourced at 95% or greater purity. All of the synthesized compounds were further tested for purity by LC/MS and were found to be pure as judged by peak height and identity.

Cmpd-23 and Cmpd-37 were synthesized as illustrated and described below. General chemistry: All chemicals and solvents, unless otherwise stated, were purchased from standard suppliers (Sigma-Aldrich, Fisher Scientific, Enamine, Combi-Blocks, Santa Cruz Biotechnology, Oakwood Chemical, and Advanced ChemBlocks) and were used without further purification. Silica gel coated with F254 fluorescent indicator on aluminum plates was used for analytical thin layer chromatography (TLC). Compounds were visualized under UV light (254 nm or 366 nm) using silica gel coated with F254 fluorescent indicator on aluminum chromatography plates (Merck, Silica gel 60 F254). Additionally, the course of reactions and compounds were monitored using standard staining procedures such as ninhydrine and KMnO₄. All compounds, unless described otherwise, were purified by flash chromatography using a Büchi Pure C 850 FlashPrep Chromatography System (BÜCHI Corporation, New Castle, DE) with silica gel cartridges (silica 12 g or 4 g; flow rate of eluent at 25 mL/min or 12 mL/min, respectively). All compounds were sourced at 95% or greater purity. All of the synthesized compounds were further tested for purity by LC/MS and were found to be pure as judged by peak height and identity.

### Synthesis of Cmpd-23

Synthesis of 7-(3-chlorophenyl)-5,6-dimethyl-4-(3-(2-(trifluoromethyl)benzyl)azetidin-1-yl)-7H- pyrrolo[2,3-d]pyrimidine-2-carboxylic acid (**3**). To a solution in a charged pressure tube, 4-chloro-7-(3- chlorophenyl)-5,6-dimethyl-7*H*-pyrrolo[2,3-*d*]pyrimidine-2-carboxylic acid, 1, (70 mg, 0.208 mmol) in dry DMF (3 mL) was added KOH (46.7 mg, 0.833 mmol), K_2_CO_3_ (10 mg, 0.0723 mmol), followed by 3- (2-(trifluoromethyl)benzyl)azetidine hydrochloride, 2 (52.4 mg, 0.208 mmol) under argon atmosphere. The reaction mixture was then stirred and heated to 110 °C for 72 h under reflux. The mixture/crude product was poured and quenched by the addition of a saturated solution of NH_4_Cl (75 mL) and acidified with aqueous 2 M HCl to adjust pH to 3–4. The resulting mixture was extracted with DCM (2 × 50 mL) and EtOAc (1 × 50 mL), and the combined organic layers were washed with brine (2 × 100 mL), dried over Na_2_SO₄, and concentrated in vacuo. The crude product was purified via Büchi Pure C 850 FlashPrep Chromatography System (4 g silica; eluting with EtOAc/CH_2_Cl_2_ from 100/0% → 20/80%; gradient) to afford **3** (105 mg, ∼85% yield) as a pale-yellow solid. Purity by HPLC-UV (254 nm) ESI-MS: 98.3%. HRMS (ESI-TOF, positive ion mode, Agilent 6224 TOF LC/MS): for C_26_H_22_ClF_3_N_4_O_2_^+^: calcd *m/z* 515.1456 [M + H] ^+^, observed *m/z* 515.1463 [M + H] ^+^.

*Tert*-butyl (*R*)-5-(7-(3-chlorophenyl)-5,6-dimethyl-4-(3-(2-(trifluoromethyl)benzyl)azetidin-1-yl)-7*H*- pyrrolo[2,3-*d*]pyrimidin-2-yl)-3-(6-methoxypyridin-3-yl)-5-oxopentanoate (**5**): To a stirred solution of 7- (3-chlorophenyl)-5,6-dimethyl-4-(3-(2 (trifluoromethyl)benzyl)azetidin-1-yl)-7H-pyrrolo[2,3- d]pyrimidine-2-carboxylic acid, 3, (92 mg, 0.180 mmol) in DMF (3 mL), HATU (115 mg, 0.300 mmol) and DIEA (260 µL, 1.43 mmol) were added and the mixture was allowed to stir for 30 min under a nitrogen atmosphere. Tert-butyl (R)-3-amino-3-(6-methoxypyridin-3-yl)propanoate chloride, 4 (57 mg, 0.200 mmol), was added to the DMF (3 mL), and the mixture was stirred for 16 h at room temperature. The mixture was then concentrated, diluted with DCM (30 mL), and washed with distilled water and brine. The organic phase was dried over Na_2_SO_4_, filtered, and then concentrated. The crude product was purified via Büchi Pure C 850 FlashPrep Chromatography System (4 g silica; eluting with EtOAc/ CH_2_Cl_2_ from 100/0% → 20/80%; gradient) to afford **5** (107 mg, 74% yield) as a yellowish, semi-solid. Purity by HPLC-UV (254 nm) ESI-MS: 96.5%. HRMS (ESI-TOF; positive mode) for C_39_H_41_ClF_3_N_6_O_4_^+^: calcd *m/z* 749.2830 [M + H] ^+^, observed *m/z* 749.2837 [M + H] ^+^.

(*R*)-5-(7-(3-chlorophenyl)-5,6-dimethyl-4-(3-(2-(trifluoromethyl)benzyl)azetidin-1-yl)-7*H*-pyrrolo[2,3-*d*]pyrimidin-2-yl)-3-(6-methoxypyridin-3-yl)-5-oxopentanoic acid (**6**). To a solution of 5 [tert-butyl (R)- 3-(7-(3-chlorophenyl)-5,6-dimethyl-4-(3-(2-(trifluoromethyl)benzyl)azetidin-1-yl)-7H-pyrrolo[2,3- d]pyrimidine-2-carboxamido)-3-(6-methoxypyridin-3-yl)propanoate], (95 mg, 0.127 mmol) in DCM (2 mL) at 0 °C was added trifluoroacetic acid (TFA, 3 mL). The reaction mixture was stirred at room temperature for 12 h, after which time the solvents were removed by rotary evaporation and washed with DCM. The crude product was purified by flash chromatography (silica; eluting with CH_2_Cl_2_/MeOH from 100/0% → 90/10%; gradient) to afford **6** (75 mg, ∼80% yield, 2 times) as a yellowish solid. Purity by HPLC-UV (254 nm) ESI-MS: 99.8%. HRMS (ESI-TOF; positive mode) for C_36_H_34_ClF_3_N_5_O_4_^+^: calcd *m/z* 692.2246 [M + H] ^+^, observed *m/z* 692.2354 [M + H] ^+^.

(R)-5-(7-(3-chlorophenyl)-5,6-dimethyl-4-(3-(2-(trifluoromethyl)benzyl)azetidin-1-yl)-7H- pyrrolo[2,3-d]pyrimidin-2-yl)-3-(6-methoxypyridin-3-yl)-5-oxopentanamide (**7**, **Cmpd-23**). To an ice-cold stirred solution of (R)-5-(7-(3-chlorophenyl)-5,6-dimethyl-4-(3-(2-(trifluoromethyl)benzyl)azetidin- 1-yl)-7H-pyrrolo[2,3-d]pyrimidin-2-yl)-3-(6-methoxypyridin-3-yl)-5-oxopentanoic acid (**6**) (70 mg, 0.101 mmol) and N-hydroxysuccinimide (13 mg, 0.130 mmol) in dry DMF (3 mL) was added EDC·HCl (25 mg, 0.130 mmol) under nitrogen atmosphere. The mixture was allowed to reach room temperature and stirred overnight. The reaction mixture was then concentrated under reduced pressure, and the residue was diluted with EtOAc (150 mL) and washed with water (3 × 50 mL). The organic phase was dried and concentrated. The crude product 1a was used for next step without further purification. 28% NH_3_ aqueous solution (0.2 mL, 2.7 mmol) was added to a solution of the product obtained above (70 mg, ∼0.101 mmol) in MeCN (3.5 mL) at room temperature. After stirring at room temperature for 9 h, the solvent and volatiles were removed in vacuo, and the solid residue was suspended in H_2_O (40 mL). The resulting mixture was extracted with DCM (3 × 50 mL), and the organic phase was dried over anhydrous Na_2_SO_4_, filtered, and concentrated. The crude product was purified via Büchi Pure C 850 FlashPrep Chromatography System (4 g silica; eluting with EtOAc/CH_2_Cl_2_from 100/0% → 20/80%; gradient) to give **7** (46 mg, 82%; over three times purification) as a pale-yellow solid. Purity by HPLC-UV (254 nm) ESI-MS: 100%. HRMS (ESI-TOF; positive mode) for C_35_H_34_ClF_3_N_7_O_3_^+^: calcd *m/z* 692.2358 [M + H]^+^, observed *m/z* 692.2355 [M + H] ^+^.

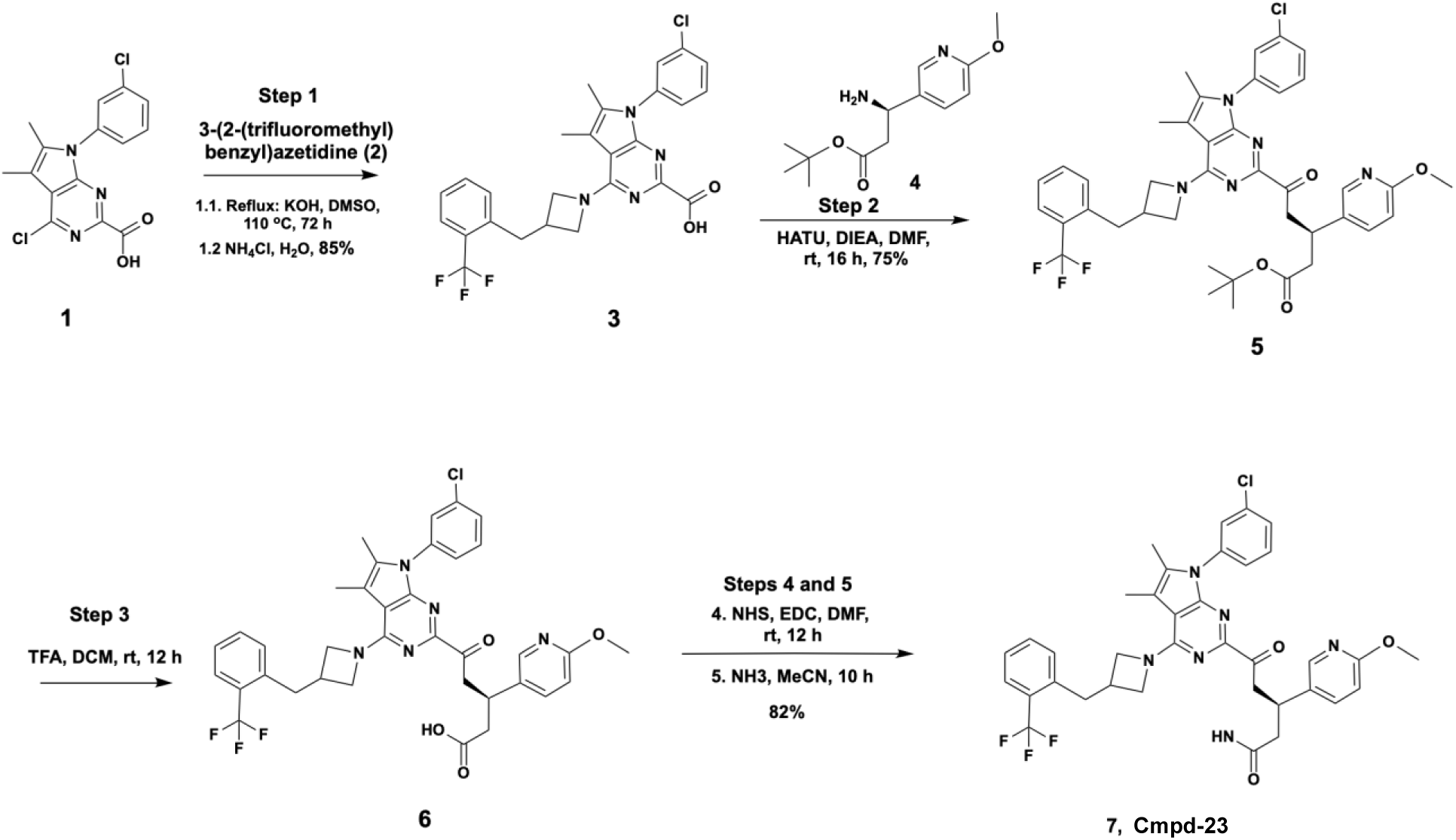

### Synthesis of Cmpd-37

2-(2-((4-chlorophenyl)thio)phenyl)-1-(3-(naphthalen-2-yloxy)pyrrolidin-1-yl)ethan-1-one (**3**, **Cmpd-37**). To a stirred solution of 2-(2-((4-chlorophenyl)thio)phenyl)acetic acid (300 mg, 1.1 mmol) in DMF (5 mL), HATU (500 mg, 1.3 mmol) and DIEA (500 mL, 2.5 mmol) were added and the mixture was allowed to stir for 30 min under nitrogen atmosphere. 2-Naphthyl 3-pyrrolidinyl ether hydrochloride (295 mg, 1.2 mmol) into the DMF (5 mL) was added and the mixture was stirred for 16 h at room temperature. The mixture was then concentrated, diluted with DCM (30 mL) and washed with distilled water and brine. The organic phase was dried over anhydrous Na_2_SO4, filtered and concentrated. The crude product was purified via Büchi Pure C-850 FlashPrep Chromatography System (4 g silica; eluting with EtOAc/CH_2_Cl_2_ from 100/0% → 40/60%; gradient) to afford **3** (370 mg, >65% yield) as white solid. *Purity by HPLC-UV (254 nm) ESI-MS: 98.4* %. HRMS (ESI-TOF; positive mode) for C_28_H_25_ClNO_2_S^+^: calcd *m/z* 474.1289 [M + H] ^+^, observed *m/z* 474.1295 [M + H] ^+^.

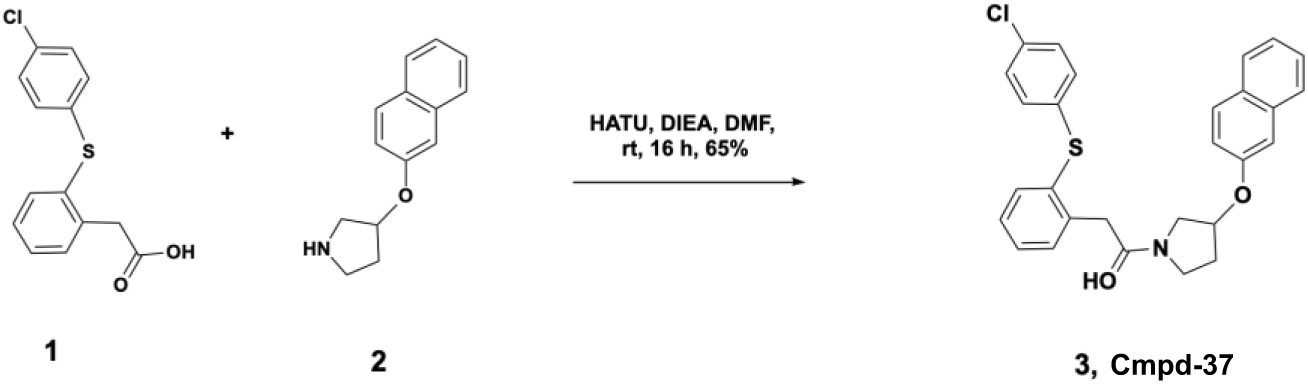

### AT1R expression and purification

Full-length wild-type human AT1R with an N-terminal hemagglutinin membrane insertion signal sequence and FLAG tag^11^ was previously cloned into pACMV-tetO^54^. For DEER experiments, minimal cystine AT1R constructs with site specific cystine mutants for spin labeling TM1-ICL2 (F55C- D236C) and TM1-TM6 (F55C-R139C) were previously generated from the wild-type construct ^27^. For protein expression, ExpiFectamine293 Reagent was used to transfect AT1R constructs into Expi239 cells as previously described ^27^. Briefly, Expifectamine293 reagent was reconstituted in OptiMEM and mixed with DNA at a 3:8 ratio of the reagent over the amount of plasmid DNA according to the manufacturer’s instructions before being added to cells. 44-48 hours post transfection, AT1R expression was induced with 4 μg/mL doxycycline, and cell media was supplemented with 5 mM sodium butyrate and 1 μM losartan. 30-36 hours post induction, Expi293 cells were harvested, and cell pellets were snap frozen in liquid nitrogen and stored for future use.

Frozen cell pellets from Expi293 cells transfected with the pACMV-tetO-Flag-AT1R plasmid construct were melted and lysed in 12.5 mL/g pellet mass of hypotonic lysis buffer (10 mM Tris, pH 7.4, 2 mM EDTA, 10 mM MgCl2, 5 units/mL benzonase, leupeptin, benzamidine and 5 µM losartan) in room temperature. Lysed cell membranes were harvested by centrifugation at 37,500 x g for 10 minutes and resuspended in 12.5 mL/g original cell pellet mass of cold solubilization buffer (20 mM HEPES pH 7.4, 500 mM NaCl, 0.5% lauryl maltose neopentyl glycol (LMNG), 0.05% CHS, 10 mM MgCl2, 5 units/mL benzonase, benzamidine, leupeptin, and 1 µM losartan). After 20 strokes of Dounce homogenization with a tight pestle on ice, cell membranes were solubilized by stirring at room temperature for 1 hour, followed by 1 hour at 4°C. After removing insoluble debris by centrifugation at 37,500 x g for 20 minutes, supernatant was supplemented with CaCl_2_ to 2 mM, and loaded onto M1- FLAG resin (2 mL/mg functional receptor expected) at the ∼2 mL/minute flow rate in a cold-room. Following two washes with 10 column volumes of ice-cold wash buffer (20 mM HEPES pH 7.4, 500 mM NaCl, 0.01% LMNG, 0.01% CHS, 2 mM CaCl_2_, benzamidine, leupeptin), the AT1R was eluted from the resin with ice-cold wash buffer in the absence of CaCl_2_ but supplemented with 0.2 mg/mL FLAG-peptide, and 5 mM EDTA. Eluted fractions containing the AT1R were confirmed by SDS-PAGE followed by Coomassie-staining, collected and concentrated using a 50 kDa MW cut-off concentrator down to less than 250 μL. Then, the monomeric AT1R was isolated and exchanged into HNM buffer (20 mM HEPES pH 7.4, 100mMNaCl, 0.01% LMNG, and 0.001% CHS) by size exclusion chromatography (SEC) on a Superdex 200 Increase 10/300 GL column (GE). The monomeric receptor fraction obtained from size exclusion chromatography was collected and concentrated to a final concentration of ∼1 mg/mL.

For purification of the spin-labeled AT1R for DEER experiments, the minimum cysteine AT1R with either the Cys 55(TM1)–139(ICL2) pair or the Cys 55(TM1)–236(TM6) pair was washed and eluted from M1 resin with the pH 6.8 buffer. The eluted receptor was incubated with a 20-fold molar excess of bis-(2,2,5,5-tetramethyl-3-imidazoline-1-oxyl-4-yl)disulfide biradical (Enzo) for 3 hours at room temperature for spin-labeling. The receptor was then applied onto a Superdex 200 Increase 10/300 GL column (GE) to remove the free spin labeling reagent and any aggregated material. The final spin-labeled AT1R was eluted with HNM (40 mM HEPES pH 7.4, 100 mM NaCl, 0.01% LMNG, and 0.001% CHS) buffer prepared with D_2_O (Cambridge Isotopes) from the column and concentrated to 20-25 μM. Ligand binding was conducted using spin-labeled and concentrated AT1R, along with 1 mM of an orthosteric peptide agonist, 0.2–1 mM of either Cmpd-23 or Cmpd-37, 20% D8-glycerol (Cambridge Isotopes), and 2% dimethyl sulfoxide (DMSO) as the vehicle (final concentration in all samples)- in a final reaction volume of 35 μL for one hour at room temperature. Right after the reaction, each sample, with a different orthosteric and allosteric ligand combination, was flash frozen with liquid nitrogen in 2.0/2.4 mm (i.d./o.d.) borosilicate capillaries (VitroCom, Mountain Lakes, NJ) and stored at -80°C.

### Purification and biotinylation of membrane scaffold protein-1 D1E3 (MSP1D1E3)

Cell pellets from *E. coli* BL21 cells expressing His-tagged MSP1D1E3 were resuspended at room temperature with 5 mL per pellet weight (gram) of resuspension buffer (50 mM Tris, pH 7.4, 300 mM NaCl, 10 mM MgCl2, 5 units/mL benzonase, leupeptin, benzamidine). TritonX-100 was added to the resuspended cells to make the 1% final concentration, and the cell lysate was mixed for 30 minutes at 4°C followed by sonication for 5 minutes (5x1 minutes) on ice. After removing cell debris by centrifugation at 34,000 x g for 30 min, the cell lysate was applied onto 2 mL/L cell culture volume of Ni-NTA resin (Goldbio) with a flow rate at 3-5 mL/minute at 4°C. The Ni-NTA resin was washed with 10 column volumes of wash buffer (20 mM HEPES, pH 8.0, 300 mM NaCl, 1% Triton X-100), followed by three sequential washes using 10 column volumes of wash buffer supplemented with sodium cholate at three decreasing concentrations (50, 20, and 10 mM, in that order). MSP1D1E3 was purified with elution buffer (20 mM HEPES, pH 7.4, 300 mM NaCl, 10 mM sodium cholate), concentrated and dialyzed with storage buffer (20 mM HEPES, pH 8.0, 100 mM NaCl, 1 mM EDTA, 10 mM sodium cholate) overnight at 4°C. The purified MSP1D1E3 protein was biotinylated using EZ-Link Sulfo-NHS-LC-Biotin (ThermoFisher) dissolved in buffer containing 20 mM HEPES, pH 7.4, 300 mM NaCl and 10 mM sodium cholate in accordance with the manufacturer’s guideline. Biotinylated protein was further purified by SEC on a Superdex 200 Increase 10/300 GL column (GE).

### HDL reconstitution of the AT1R

The purified, functional AT1R was reconstituted into high-density lipoprotein (HDL) particles using established protocols described in previous studies^55^. Briefly, FLAG-tagged AT1R was incubated for 1 hour at 4°C with a 50-fold molar excess of biotinylated MSP1D1E3 and 8 mM POPC:POPG lipids (3:2 molar ratio; 1-palmitoyl-2-oleoyl-sn-glycero-3-phosphocholine and 1-palmitoyl-2-oleoyl-sn-glycero-3- phospho-[1′-rac-glycerol]) (Avanti Polar Lipids; Alabaster, AL). Detergent removal was achieved by overnight incubation at 4°C with BioBeads SM-2 (Bio-Rad; Hercules, CA). The resulting HDL particles containing the receptor were then purified using FLAG-M1 affinity chromatography.

### DNA-encoded library (DEL) synthesis

Libraries were built by a tagged-split-and-pool chemistry approach (Chemetics®) at Amgen Research Copenhagen^53^.

### Affinity DEL Selection

Approximately 60 μg biotinylated AT1R-HDL particles, containing ∼24 μg (0.57 nmols) of the AT1R, were immobilized on 25 μL of NeutrAvidin beads (Thermo Fisher Scientific; Waltham, MA) by incubation for 60 minutes in binding buffer (20 mM HEPES, pH 7.4, 150 mM NaCl, 10 mM MgCl_2_, one of the orthosteric peptide agonists at 1 mM). For the case of TRV055, TRV055 was used together with nanobody (Nb)110i1 at ∼1 μM through the whole procedure. Lyophilized library molecules were dissolved in 50 μL of water and dried out again before reconstitution in 25 μL of the binding buffer supplemented with 1 mg/mL of sheared salmon sperm DNA (ssDNA) (Ambion; Waltham, MA). Prior to re-lyophilization after dissolving water, 1 μL of library molecules were allocated for later quantitative PCR (qPCR). The bead-bound AT1R-HDLs were then mixed with library molecules and incubated for 60 minutes at room temperature under vigorous shaking. Following this incubation, the beads were transferred to a micro-column connected to a vacuum apparatus and subsequently washed three times with 100 μL of ice-cold binding buffer containing 1 mg/mL ssDNA. Free biotin at 1 mM and 0.005 % Tween-20 were supplemented into the first and second washing steps to reduce the background non-specific binding of library molecules. During each of the washing steps, excess liquid was removed via vacuum. To release the bound compounds, the NeutrAvidin beads were subjected to two sequential elution steps using 50 μL of water containing 1.5% Fos-choline (Avanti Polar Lipids; Alabaster, AL). The first incubation was performed at 37°C for 15 minutes, followed by a second at 72.5°C for another 15 minutes. After each incubation, the supernatant was collected by centrifugation at 1,000 × g for 1 minute from the beads. To reduce the probability of losing specific binders, 1 μL of 10 mg/mL ssDNA was added to the collected supernatant before it was processed using a nucleotide removal kit (Qiagen; Hilden, Germany) to eliminate denatured proteins and lipid contaminants. The enriched pool of bound compounds was subsequently eluted from the column with 50 μL of water. From this purified eluate, 1 μL was set aside for quantitative PCR analysis, while the remainder was either subjected to an additional round of selection using fresh AT1R-HDL particles or prepared for next-generation sequencing.

### Quantitative PCR (qPCR)

To assess remaining library molecules after each round of affinity selection, library DNA was quantified using quantitative PCR (qPCR) following each round of selection. qPCR reactions were run in JumpStart Taq ReadyMix (Sigma-Aldrich, St. Louis, MO) in accordance with the manufacturer’s protocol using a primer set consisting of a universal forward primer (5′-CAAGTCACCAAGAATTCATG-3′), a library aliquot-specific reverse primer, and a FAM/TAMRA-labeled probe (5′-CAGACGACCTAGGATCACC- 3′). Pre-selection library samples were utilized to generate a standard curve, using at least four serial dilutions on a logarithmic scale which enabled the determination of DNA copy numbers in samples collected after each selection round. DNA samples assessed by qPCR were either directly amplified or diluted in TE buffer supplemented with single-stranded DNA (ssDNA). Reactions were carried out on a QuantStudio-5 Real-Time PCR System (Applied Biosystems, Waltham, MA).

### Next generation sequencing and deconvolution

To facilitate target identification, samples were sequenced using the Ion Torrent Personal Genome Machine (PGM; Thermo Fisher Scientific) in accordance with the manufacturer’s protocols for single-end amplicon sequencing. Briefly, affinity-selected samples were first amplified under two rounds of PCR. The initial round increased DNA yield and facilitated sample tracking, and the second round incorporated the necessary sequencing adapters required for emulsion PCR. In the first round of amplification, a library-specific forward primer (5′-AAGGCCTAGATTCACTCACG-3′) and a unique reverse primer assigned to each library aliquot at the initiation of affinity selection were used to expand DNA molecules. The unique reverse primer facilitated accurate sample tracking and served as an identifier for individual selections. In the second PCR round, primers containing Ion Torrent adapter sequences were used to append sequencing adapters to the amplicons. A five-nucleotide sorting barcode was incorporated between the adapter sequences and the library-specific template, enabling sample multiplexing. The resulting PCR products were purified via gel electrophoresis and quantified using qPCR as previously described, with the exception that the standard curve was generated from plasmid DNA encoding a sequence-verified Ion Torrent library insert. For each emulsion PCR (emPCR) reaction, 4×10⁸ DNA molecules were utilized. emPCR was performed on the Ion OneTouch 2 system (Thermo Fisher Scientific), and the resulting products were manually loaded onto an Ion 318 chip for sequencing. Sequence data were analyzed using a custom BLAST-based algorithm facilitating the identification of synthetic templates from their corresponding coding sequence. This process enabled the deconvolution of each sequence into its defined chemical structure. Following this deconvolution, analysis of sequence frequency between affinity-selected samples and control samples (enriched in the absence of target protein) was used to assess the enrichment and significance of compound selection.

### Radio-ligand binding

Assays were performed largely in accordance with established protocols^27^ utilizing plasma membrane prepared from Expi293 cells transiently transfected with the pACMV-tetO-Flag-AT1R inducible expression plasmid^27^ or the AGTR2-DuET construct^56^ that was a gift from Thomas Sakmar (Addgene plasmid # 213184 ; http://n2t.net/addgene:213184; RRID:Addgene_213184). Experiments were conducted in a buffer containing 10 mM Tris-HCl (pH 7.4), 1 mM EDTA, 25 mM MgCl_2_, and 0.1% bovine serum albumin (BSA). For competition binding experiments, each 250 μL reaction mixture included appropriate amounts of plasma membrane, 4 nM [³H]-olmesartan (specific activity: 10 Ci/mmol), varying concentrations of allosteric compounds, and either a serial dilution or a fixed concentration of orthosteric agonists—selected to approximate IC_20_ competition levels. Non-specific binding was assessed by including 20 μM candesartan in parallel reactions. For [^3^H]-AngII binding, 150 μL reaction mixture included appropriate amounts of plasma membrane, approximately 10 nM [^3^H]- AngII (specific activity: 60 Ci/mmol), and either 50 µM allosteric compounds or 1 µM purified trimeric Gq protein. The amount of non-specific binding was determined in the presence of high-affinity ligands (20 μM Olmesartan for the AT1R and 10 μM CGP-42133 for the AT2R). After a 90-minute incubation at room temperature, binding reaction was terminated via rapid filtration through GF/B glass fiber filters (Brandel). Filters were then washed with 8 mL of ice-cold binding buffer (10 mM Tris-HCl, pH 7.4, 1 mM EDTA, 25 mM MgCl_2_) using a Brandel cell harvester. Radioactivity associated with bound [³H]- olmesartan was measured using a Tri-Carb 2800TR (PerkinElmer) beta counter after a 24-hour extraction in Lefko-fluor scintillation cocktail. Results were reported as specific binding by subtraction of the non-specific binding count from each of the total counts.

### Measurements of intracellular calcium levels

The level of intracellular calcium was monitored using FLIPR-6, an intracellular calcium sensing fluorophore according to the manufacturer’s guidelines (Molecular Devices) with slight modifications. For initial measurement of 37 putative AT1R binders from the DEL screen, HEK-293 cells, stably overexpressing the AT1R, were utilized. For comprehensive characterization of Cmpd-23 and -37, U2OS cells, stably transfected with an inducible human AT1R expression plasmid construct, were utilized in the absence of an inducing reagent to minimize receptor expression. These HEK-293 or U2OS cells stably expressing the AT1R were plated at ∼50,000 or ∼30,000 cells per well, respectively into black, clear-bottom 96-well plates one day prior to the measurement. The following day, lyophilized FLIPR-6 reagents were reconstituted in 10 mL of Hank’s Balanced Salt Solution (HBSS) supplemented with 20 mM HEPES, pH 7.4, and the reconstituted reagents were further diluted 6-fold in the same buffer. After incubation of the cells with the reagents for 1.5 hours at 37°C, cells were pretreated with either DMSO (vehicle control) or varying concentrations of allosteric compounds diluted in HBSS supplemented with extra 20 mM HEPES, pH 7.4 and 0.05% bovine serum albumin (BSA) for 15-30 min. While monitoring intracellular calcium levels using a FlexStation microplate reader (Molecular Devices) at 37 °C, cells were stimulated with orthosteric agonists in a dose-dependent fashion, and intracellular calcium levels were continuously monitored for indicated times (routinely 2-3 min). For generating agonist dose-response curves, area under the time-dependent curve (AUC) generated upon stimulation with an agonist at each concentration was calculated.

For measuring the intrinsic agonistic activity of Cmpd-23 and -37, U2OS cells stably expressing the AT1R were plated, and while monitoring intracellular calcium levels at 37 °C, cells were stimulated with AngII, Comp-23, or Cmpd-37 in a dose-dependent manner to generate dose-response curves as described above.

To monitor changes in intracellular calcium levels in neonatal cardiomyocytes, cells were isolated as described in the section for “Cardiomyocyte Isolation” below. Cells were subsequently plated at a density of ∼50,000 cells per well and maintained for 5-7 days before the experiment. On the day of measurement, cells were incubated with reconstituted and further diluted FLIPR-6 reagents, as described for the U2OS cells above, for 1.5 hours at 37°C. Then, cells were pretreated with DMSO, 30 µM Cmpd-23, or 10 µM Losartan diluted in HBSS supplemented with extra 20 mM HEPES, pH 7.4 and 0.05% BSA for 15-30 min. While monitoring intracellular calcium levels at 37 °C, cells were stimulated with AngII in a dose-dependent manner. Intracellular calcium levels were continuously monitored to generate dose-response curves as described for the U2OS cells above.

### Measurements of β-arrestin recruitment

β-Arrestin recruitment was evaluated using the PathHunter™ chemiluminescence-based enzyme fragment complementation assay (Eurofins), following previously published protocols^24^ with minor modifications. CHOK1 cells stably expressing β-arrestin2 tagged with an enzyme acceptor and ProLink-tagged AT1Rs were seeded into white, clear-bottom 96-well plates at a density of 30,000 cells per well. The next day, cells were pre-treated with either 0.3% DMSO or varying concentrations of allosteric modulators diluted in HBSS supplemented with 20 mM HEPES, pH 7.4 and 0.05% BSA. Afterward, cells were stimulated with agonists for 30 to 45 minutes at 37°C. The assay was terminated by adding the PathHunter detection reagents (Eurofins), followed by an additional incubation for 1 hour at 27°C. Luminescence was measured using a ClarioStar microplate reader (BMG Labtech).

### β-arrestin endocytosis assay

β-Arrestin endocytosis was quantified using a NanoLuc enzyme complementation assay, adapted slightly from previously described methods^57^. Human AT1R was transiently expressed in HEK-293 cells along with the endosome-targeted FYVE domain of endofin attached to the N-terminal fragment of NanoLuc (EeN) and β-arrestin2 attached to the C-terminal fragment of NanoLuc ^53^ in a 1:1:1 ratio using FuGENE 6 (Promega) according to the manufacturer’s instructions with a 5:1 ratio of the reagent over the amount of plasmid DNA. The following day, transfected cells were seeded into white, clear-bottom 96-well plates at a density of 60,000 cells per well, treated with either DMSO or 30 μM Cmpd-23 diluted in HBSS supplemented with 20 mM HEPES, pH 7.4 and 0.05% BSA, and then stimulated with varying concentrations of AngII for 15 minutes. Endocytosis of β-arrestin was monitored by measuring chemiluminescence signals generated through the complementation of NanoLuc fragments within endosomes in the presence of coelenterazine H (Cayman Chemical Company). Luminescence was recorded using a ClarioStar microplate reader (BMG Labtech).

### Cardiomyocyte Isolation

All animal experiments were approved by the Institutional Animal Care and Use Committee (IACUC) at Duke University Medical Center and performed in accordance with relevant guidelines and regulations. Ventricular cardiomyocytes were freshly isolated from 12-16-week-old C57BL/6J wild-type and homozygous β-arrestin2 KO mice using a standard Langendorff perfusion system^4^. The heart was initially cannulated through the aorta and then perfused with an oxygenated buffer at 37 °C (120 mM NaCl, 14.8 mM KCl, 0.6 mM KH_2_PO_4_, 0.6 mM Na_2_HPO_4_, 1.2 mM MgSO_4_×7H_2_O, 10 mM HEPES, 4.6 mM NaHCO_3_, 30 mM taurine, and 5.6 mM glucose, pH 7.3). After 2 minutes, the buffer was switched to a digestion solution containing 2.4 mg/mL collagenase (Worthington). Following 8 minutes of digestion, the ventricles were transferred to a stopping buffer that included 10% calf serum. Myocytes were then manually dissociated and brought to a physiological calcium concentration (1.2 mM).

Primary neonatal rat ventricular myocytes were isolated from 1-2 day-old Sprague Dawley rat pups (n = 20) as previously described^58^. Briefly, ventricles were excised, minced, and subjected to three serial digestions in ADS buffer containing pancreatin and collagenase II at 37°C, shaking at 80 rpm. Supernatants from the second and third digestions were combined with Ham’s F-10 containing 10% FBS, and centrifuged. Cells were resuspended in Ham’s F-10 complete medium, filtered through a 70μm strainer, and pre-plated for 2 hours to reduce fibroblast contamination. Non-adherent cardiomyocytes were collected, counted via trypan blue exclusion, and seeded onto Poly-D-Lysine 96-well plates at 1×105 cells/well. Cells were maintained in Ham’s F-10 complete medium (10% horse serum, 5% FBS, 1% penicillin-streptomycin) on Day 1, followed by Ham’s F-10 with 5% FBS, 1% penicillin-streptomycin medium on Day 2 onward, with daily media changes until the assay was performed on Day 7.

### Contractility Study

Isolated adult cardiomyocyte was first pre-treated with DMSO (0.1%) or Cmpd-23 (25 µM) for 20 minutes before being stimulated with HBSS as basal control or angiotensin II (10 µM) as positive control. The cells were plated in a chamber (Ionoptix, Westwood, MA) on a Nikon Eclipse TE300 inverted microscope (40X 0.9 NA objective, MRF00400, Nikon). Cardiomyocytes were paced at 1 Hz and 20 V (MyoPacer, Ionoptix), and sarcomere length was documented by the MyoCam-S camera (Ionoptix). Ten consecutive contractions per cell (8-12 cells per condition) were averaged to quantify contractility and kinetic parameters (IonWizard 7.2). Only those cells exhibiting rod-shaped single cardiomyocyte morphology and normal responsiveness to electrical stimulation were included. Contractility measurements were obtained from 8 to 12 isolated cardiomyocytes per heart and averaged for a single data point. Statistical comparisons between conditions were performed using one-way ANOVA followed by Dunnett’s post-hoc test.

### Expression and purification of protein components of the Cmpd-23 complex and complex formation

WT human AT1R was expressed and purified from Expi293 cells as described above. The active state stabilizing nanobody (Nb) AT110i1 was expressed and purified as described previously^40^. Briefly, Nb AT110i1 was cloned into a pET26b vector and transformed into T7 Express lysY BL21 *E. coli*. Transformed bacterial cells were grown in terrific broth media containing 1 mM MgCl_2_ and 0.01% glucose at 37°C and protein expression was induced with 1mM IPTG upon reaching an OD600 =∼0.7. The incubation temperature was lowered to 27°C, and cells were grown overnight before being harvested. Bacterial cell pellets were then resuspended and lysed in osmotic shock buffer (200 mM Tris-HCl pH 7.4, 0.5 mM EDTA, 500 mM Sucrose, 2x leupeptin, 2x benzamide). AT110i1 was then purified with Ni-NTA chromatography as described^40^ before being dialyzed into nanobody buffer (20 mM HEPES, pH7.4, 100 mM NaCl), and snap frozen for future use.

The legobody construct consists of two separate protein components, a Fab (legobody-Fab) and MBP- Protein A fusion protein (Legobody-MBP) which were purified separately. Legobody-Fab heavy chain and light chain plasmids were separately cloned into pCAGEN plasmids and Legobody-MBP was cloned into a pET26b plasmid^42^. All legobody plasmids were obtained from Addgene. The Legobody-Fab is construct was expressed in Expi239 cells utilizing the same transfection method for the AT1R inducible plasmid transfection as described above. At ∼12-16 hours post transfection 10 mM sodium butyrate was added to the transfected cells and at∼60 hours after transfection, media containing secreted Fabs was harvested. Fabs were purified using Ni-NTA chromatography as described before^59^ being dialyzed into legobody storage buffer (20 mM HEPES, pH 7.4, 150 mM NaCl) and snap frozen for future use.

Legobody-MBP fusion was expressed as previously described^42^. Briefly, T7 Express lysY BL21 *E. coli* were transformed with the expression plasmid and grown in terrific broth media. Upon reaching an an OD600 =∼0.7, cells were induced with 1mM IPTG and were grown at 37°C for an additional 4 hours before being harvested. The cell pellet was resuspended and was sonicated in purification buffer (25 mM HEPES pH 7.4, 400 mM NaCl, 20 mM imidazole, 1x leupeptin, 1x benzamide, and 5 units/mL benzonase). Legobody MBP was then purified using Ni-NTA chromatography as described before.^59^ Legobody-MBP was dialyzed into legobody buffer (20 mM HEPES, pH 7.4, 150 mM NaCl) and snap frozen for future use.

The AT1R-TRV023-Cmpd-23-AT110i1 complex was formed based on previous complex formation conditions for the AT1R-BRIL-TRV023-AT110i1 complex^41^. Subsequently, additional legobody Fab and legobody MBP were added following the initial complex formation. Briefly, AT1R and AT110i1, were incubated together at a ratio of 1:1.1, in the initial presence of 50 µM TRV023 and Cmpd-23. After overnight incubation at 4°C, excess Legobody components were added at a ratio of roughly 1:1.5 and incubated for an additional 2 hours. Both ligands were diluted down to a final concentration of 50 µM and the full complex was then applied to SEC in SEC buffer (25 mM HEPES pH 7.4, 100 mM NaCl, 2 mM maltose, 50 uM TRV023/Cmpd-23, 0.002% LMNG). Fractions corresponding to the full complex were collected and concentrated to ∼5 mg/mL to be applied to grids.

### AT1R-Giq expression, complex formation and purification

For expression and purification of the AT1R-Giq complex, modifications to the AT1R construct were introduced as described below to facilitate AT1R-G protein complex formation in Sf9 cells. The cDNA encoding human AT1R (NP_000676.1) was cloned into a modified pFastBac1 vector containing an N-terminal influenza haemagglutinin (HA) signal peptide for receptor protein expression. For affinity purification and improved protein yield, an N-terminal FLAG tag (DYKDDDDK) was added, followed by sequential fusion of maltose-binding protein (MBP) and BRIL (apocytochrome b562RIL) tags to the N-terminus of AT1R. To promote complex formation, the NanoBiT tethering system was employed as previously described^60^: the C-terminus of the AT1R was fused to the large NanoBiT fragment (LgBiT), and the C-terminus of Gβ to the small fragment (SmBiT). The engineered human Gαiq subunit, derived from Gαi with its α5-helix replaced by that of Gαq, was generated for AT1R–Gαiq complex formation as previously described^61^. All three G protein subunits—Gαiq, human Gβ1, and human Gγ2—were individually cloned into the pFastBac1 vector. Additionally, the scFv16 construct was cloned into the same vector, containing an N-terminal GP67 secretion signal and a C-terminal 6×His tag.

Sf9 insect cells were used for baculovirus production and protein expression. To generate high-titer recombinant baculovirus, 3 μg of recombinant bacmid DNA was mixed with 9 μL of FuGENE transfection reagent in 1 mL of transfection medium (Expression System) and incubated at room temperature for 20 minutes. Subsequently, 1 × 10⁷ Sf9 cells were resuspended in the transfection mixture and cultured at 27 °C with shaking at 110 rpm. Four hours post-transfection, 4 mL of ESF921 medium was added, followed by an additional 5 mL at 48 hours. After 96 hours of incubation, the culture was centrifuged, and the P0 baculovirus was harvested from the supernatant. The P0 virus was used to infect fresh Sf9 cells to generate P1 baculovirus. For protein complex expression, Sf9 cells at a density of 3×10⁶ cells/mL were co-infected with baculoviruses encoding AT1R-LgBiT, Gαiq, Gβ1-SmBiT, Gγ2, and scFv16. Forty-eight hours post-infection, cells were harvested by centrifugation, and the pellets were stored at −80 °C until use.

Sf9 cell pellets expressing AT1R-LgBiT, Gαiq, Gβ1-SmBiT, Gγ2, and scFv16 were resuspended in buffer containing 20 mM HEPES (pH 7.4) and 100 mM NaCl, supplemented with leupeptin, benzimidamide hydrochloride, apyrase, 20 μM AngII, and 50 μM Cmpd-37. The suspension was homogenized and incubated at room temperature for 2 hours to promote complex formation. Membrane solubilization was performed by adding 0.5% (w/v) LMNG and 0.05% (w/v) cholesteryl hemisuccinate Tris salt (CHS; Anatrace), followed by centrifugation at 25,000 rpm for 30 minutes. The resulting supernatant, containing the solubilized protein complexes, was incubated with Dextrin Beads for 2 hours at 4 °C. After centrifugation at 25,000 rpm for 30 minutes, the supernatant containing the protein complexes was collected and incubated with Anti- FLAG M1 affinity beads for 2 hours. The resin-bound complexes were packed into a column and washed with buffer containing 20 mM HEPES (pH 7.4), 100 mM NaCl, leupeptin, benzimidamide hydrochloride, 0.01% (w/v) LMNG, 0.001% (w/v) CHS, 20 μM AngII, and 50 μM Cmpd-37. The protein complexes were eluted using buffer supplemented with 20 mM HEPES, 100 mM NaCl, 10 mM maltose, 0.01% (w/v) LMNG, 0.001% (w/v) CHS, 20 μM AngII, and 50 μM Cmpd-37. Eluted fractions were concentrated and further purified by size-exclusion chromatography on a Superose 6 Increase 10/300 GL column equilibrated with buffer containing 20 mM HEPES (pH 7.4), 100 mM NaCl, 0.00075% LMNG, 0.00025% glyco-diosgenin (GDN), 0.0002% CHS, 20 μM AngII, and 50 μM Cmpd-37. Fractions corresponding to the target complex were collected and concentrated for cryo-EM sample preparation.

### Grid preparation and cryo-EM data collection

For cryo-EM grid preparation, 3 µL of sample (AT1R-TRV023-Cmpd-23-AT110i1-legobody) was applied to glow-charged UltrAuFoil Holey Gold Films (R 1.2/1.3, Electron Microscopy Sciences) and were vitrified using a Vitrobot Mark IV at 4°C and a humidity of 100%. Samples were subsequently imaged on a Krios G3i (Thermo Fisher) transmission electron microscope equipped with a Gatan K3 detector operated at 300 kV. Images were collected at 81,000x magnification (nominal pixel size of 1.1Å) with a defocus range of -1.0 to -2.0 μm, and each micrograph was imaged with a total dose of 56.3 e⁻/Å² was fractionated into a movie stack of 60 frames.

2.5 μL of AT1R-Giq-Angll-Cmpd-37 complex was dispensed onto glow-charged holey carbon grids (Quantifoil, Au, R1.2/1.3, 300 mesh) and left for 10s. The excess solution was removed by blotting with filter paper, and then the grid was plunged in liquid ethane. All steps above were carried out using a Vitrobot Mark IV at 4C, with humidity of 100%. Data were collected using a FEI Titan Krios microscope operated at 300 kV, equipped with a Gatan K3 Summit counting camera and BioContinuum. EPU software was used for automated data collection. Movies were acquired at a nominal magnification of 105,000x across a defocus range of -1.0 to -2.0 μm and with a pixel size of 1.06 Å. An accumulated dose of 60 e⁻/Å was fractionated into a movie stack of 32 frames.

### Cryo-EM data processing and 3D reconstruction

13002 dose-fractionated movies were collected for the AT1R-TRV023-Cmpd-23-AT110i1-legobody complex and were processed in CryoSPARC v4.0.155. Micrographs were first subject to patch-based motion correction and CTF estimation. An initial subset of 500 micrographs was then used for blob picking resulting in 643,085 particles which were subject to 2D classification and 3D Ab-initio Reconstruction, generating 1 good and 3 bad classes. The resulting good class was used to generate templates for template picking for the entire dataset. 3,344,343 resulting particles were extracted and classified by 3D ab initio reconstruction followed by iterative rounds of heterogeneous refinement. The best-resolved class consisting of 907,280 particles was selected for homogeneous refinement followed by local refinement of the receptor-nanobody region, resulting in a density map at a nominal resolution of 2.97 Å (determined by gold standard FSC using the 0.143 criterion).

The data processing workflow for the AT1R-Giq-Angll-Cmpd-37 complex involved the use of AutoEMage^62^ and cryoSPARC v.4.7.1^63^. Dose-fractionated movies were collected and underwent patch motion correction in AutoEMage. Subsequently, the motion-corrected and dose-weighted micrographs were imported into cryoSPARC for determining the contrast transfer function (CTF) parameters through patch CTF estimation. Local resolution estimation was carried out using Local Resolution Estimation in cryoSPARC, followed by map sharpening with DeepEMhance^53^. A total of 21,261,984 particles were automatically picked from 9,512 micrographs and subjected to multiple rounds of 2D classification for cleaning. Subsequently, 1,898,999 selected particles underwent 3D Ab-Initio Reconstruction and Hetero Refinement. A final round of 2D classification and Rebalance 2D Classes and Rebalance Orientations were used for particle selection, and 276,067 particles were utilized for 3D NU-refinement and Local refinement, resulting in a density map at a nominal resolution of 3.15 Å (determined by gold standard FSC using the 0.143 criterion).

### Model Building and Refinement

Model building and refinement for the AT1R-TRV023-Cmpd-23-AT110i1-legobody complex utilized the existing AT1R-BRIL-TRV023-AT110i1 model (PDB:6OS1) as an initial model, in which the BRIL fusion was deleted. The model was rigid body fitted into the density map in UCSF Chimera and iteratively adjusted in Isolde^64^ and Coot^65^.

Model building and refinement for the AT1R-Giq-AngII-Cmpd-37 complex utilized the AlphaFold2 (AF2) predicted models as initial models^66^. These models were rigid body fitted into the density maps using UCSF Chimera^67^ and then iteratively adjusted in Coot^65^.

Further refinement was carried out using phenix. Real_space_refine^68^. Comprehensive model validation was performed using phenix.mtriage ^68^. The cryo-EM data collection, 3D reconstruction, and model refinement statistics are summarized in Table S1. Structural figures were generated using PyMOL or ChimeraX ^69^.

### DEER Spectroscopy and Analysis

4-Pulse deadtime-free DEER was performed at Q-band (34 GHz) on an upgraded ElexSYS 580 (Bruker, Rheinstetten) equipped with an arbitrary waveform generator and a 150 W TWT amplifier. Pulse lengths were optimized via nutation experiments and set to 16 ns/32 ns for 90°/180° pulses, respectively, and the adiabatic pump pulse was realized as a 100 ns/50 MHz chirp pulse 80 MHz above the observe frequency. Temperature was adjusted to 50 K using a cryogen-free cooling system.

DEER analysis was performed in DEERlab (v0.92)^70^ using a Gaussian mixture model and global fitting of all DEER data for the same spin pair. Spin concentration was used as a global fitting parameter of the 3D background model. The most parsimonious model was determined via the Bayesian information criterion (BIC). Confidence bands are calculated via bootstrapping analysis (1000 steps). Raw DEER data including all pulse parameters is accessible via Zenodo under https://doi.org/10.5281/zenodo.15814162. In silico spin labeling of AT1R and distance estimates were calculated in ChiLife (v1.1.3)^71^ using only the receptor and Cmpd-23 or Cmpd-37, but with all other molecules removed. We used the V1M implementation with 1000 step repacking.

### Molecular Dynamics analysis

We analyzed molecular dynamics (MD) simulations performed as part of a previous study^44^. The simulations analyzed are apo (unliganded), AngII-bound, TRV023-bound, TRV026-bound, and TRV055- bound simulations started from the previously published S1I8-bound AT1R structure (PDB 6DO1). In these simulations, the stabilizing nanobody and S1I8 were removed, without intracellular restraints and with D125^3.49^ protonated (neutral). Ligands were modeled into the crystal structure using *in silico* mutation using Maestro (Schrödinger) to match those of each peptide ligand.

TM7 twist was calculated by aligning each simulation frame or PDB structure to the AngII-bound AT1R structure (PDB 6OS0) using the Cα atoms from transmembrane (TM) helices 1–4, which was itself aligned to TM1–4 of the inactive AT1R crystal structure (PDB 4YAY)^72^ in the Orientations of Proteins in Membranes (OPM) database^73^. Next, a vector was computed by taking the difference in XY position of the Cα atoms of C296^7.47^ and N295^7.46^. We report the angle between these vectors in each simulation frame and the reference structure. TM3 twist was calculated in a similar way, except taking the average of the angle between the Cα atoms of L112^3.36^ and N111^3.35^, and of N111^3.35^ and F110^3.34^.

The simulation traces shown in Fig. 4f are from the same AT1R-TRV055 simulation. In all figures showing simulation traces, raw simulation traces are shown in light gray, and smoothed traces are shown in dark gray (smoothing window of 30 ns). In all renderings, unless otherwise noted, structures were aligned on all Cα atoms of AT1R. The Cmpd-23 structure was aligned based on all Cα atoms of AT1R to the canonical and alternative active conformations. Cmpd-23 was positioned (as a rigid body) to best fit between TM6 and TM7 in the inactive conformation, since direct alignment did not place Cmpd-23 in the correct binding location. The TM7 twist measurement of the alternative (orange in Fig. 4f) and canonical (blue in Fig. 4f) are 0.3° and 54.3°, respectively.

### Mutagenesis study via TruPath G protein dissociation assay

For mutagenesis experiments, mutations on AT1R residues surrounding the binding site of Cmpd-23 (I241A and I245A, F301A, K310A) and Cmpd-37 (F110A, W153A, K146A) were generated utilizing the Quickchange Lightning Site Directed Mutagenesis Kit according to the manufacturer’s protocol (Agilent). Gq protein activation was determined by monitoring dissociation of Gαq and Gβγ subunits utilizing a well-established TruPath assay^49^. HEK-293 cells were transiently transfected with either WT or a mutant AT1R, Gαq-RLuc8, Gβ3, and Gγ1-GFP2 expression plasmid constructs in a 0.5 or 1, respectively:1:1:1 ratio using FuGENE 6 (Promega) according to the manufacturer’s instructions with a 5:1 ratio of the reagent over the amount of plasmid DNA. The next day, cells were plated into poly-D-lysine-coated white, clear-bottom 96-well plates at a density of 60,000 cells per well. One more day after plating, media was replaced with HBSS buffer supplemented with 20 mM HEPES, pH 7.4 and the Prolume Purple cell permeable luciferase substrate (Nanolight Technology) at 1 μM followed by treatment of either the DMSO vehicle control or 30 μM of allosteric Cmpds diluted in HBSS supplemented with 20 mM HEPES, pH 7.4 and 0.05% BSA. After incubation for indicated times, cells were stimulated with AngII in a dose-dependent fashion, and Gαq and Gβγ dissociation was determined by monitoring loss of luciferase signal using a ClarioStar microplate reader (BMG Labtech). Changes in BRET ratio (525-530 nm signal/400 nm signal) were measured at 8 minutes (for Cmpd-23) or were expressed as the mean values from four measurements up to 8 minutes after stimulation with AngII.

### Statistical Analyses

All of the dose-response curve fits were obtained using the computer program GraphPad Prism. Statistical analyses for the shift of curves in Fig. 1d-i, and Extended Data Fig. 2d,e,k,m,n were performed using ‘two-way ANOVA’. ***, P < 0.001 compared to the control curve obtained in the presence of the vehicle (DMSO). Statistical analyses for the results in Fig. 2c-f, 6b,h and Extended Data Fig. 2c,l,o were performed using ‘one-way ANOVA’ with ‘Dunnet’s’ multiple comparison post-tests. *, P<0.05, **, P < 0.01; ***, P < 0.001 compared to the control value obtained either in the presence of the DMSO vehicle (Fig. 2c-f and Extended Data Fig. 2c,l,o) or in WT receptor-transfected cells. (Fig. 6b,h). Allosteric values, shown in Extended Data Table 1, were quantified by extended allosteric ternary complex model^29^ using the competition binding data (Fig. 2d,e) and combined operational model of allostery^30–32^ using the functional assay data (Fig. 2f-i).

## Data availability

The atomic models of the AT1R-Cmpd-23-TRV023-AT110i1 complex and the AT1R-Cmpd-37-AngII- Giq complex have been deposited into the Protein Data Bank. The corresponding 3D cryo-EM density maps have been deposited into the Electron Microscopy Data Bank. Raw DEER data including all pulse parameters is accessible via Zenodo under https://doi.org/10.5281/zenodo.15814162.

## Acknoledgements

R.J.L. is a Howard Hughes Medical Institute (HHMI) investigator. This work was supported, in part, by US National Institutes of Health (Grant R01HL16037 to R.J.L., R01HL056687 to H.A.R., R01GM127359 to R.O.D., HL134608 (Project 2) to W.J.K, HL16037-45S1 to A.W.K.), the St. Jude Children’s Research Hospital Collaborative Research Consortium on GPCRs to R.J.L, the German Research Foundation, project number 421152132, subproject A03 and project number 514664767, subproject B07 to M.E., the Jules Stein Endowed Chair to W.L.H. E.J.F. is in the Sarafan ChEM-H Chemistry/Biology Interface Training Program and supported by NIH 5T32GM139791. We thank Sudarshan Rajagopal (Duke University) for his technical advice to calculate allosteric factors. We thank also Biswaranjan Pani, Xiang Zhang, Xingdong Zhang, Haoran Jiang (Duke University), and Duke undergraduate students (Saajan Patel, Alina D. Fernandez, Daliya Rizvi, Jeffrey B. Guerra and Alice Zhang) for their technical advice and assistance. We acknowledge the use of Transmission Electron Microscopy at the Shared Materials Instrumentation Facility (Duke University). We are also grateful to Icee Li for excellent secretarial assistance.

## Author contributions

S.A., D.P.S, and R.J.L. conceived the study. S.A., A.M., and A.L conducted DEL screening. M.V., E.O., G.M., and J.K.M performed DEL screening analysis and synthesis of initial candidate compounds. A.K. and P.X. synthesized Cmpd-23 and Cmpd-37. S.A., S, L., P.X and R.R. designed and conducted *in vitro* and cellular functional experiments. R.T.S. and S.A. performed calculation of allosteric factors. J.K. prepared membranes for the binding experiments. E.Y.L. and A.J isolated adult cardiomyocytes and carried out contractility experiments with assistance of Haoran Jiang. S.M.K. isolated neonatal cardiomyocytes under supervision of W.J.K. S.L., P.X., and S.A. performed protein purification, complex formation, and EM image acquisition. S.L. and P.X., carried out EM data processing and analysis with supervision of N.P. S.A. and S.L. prepared the spin-labeled AT1R for DEER and M.E. conducted DEER experiments and analysis. E.J.F and C.-M.S. performed structural analysis, and performed and analyzed MD simulations, under supervision of R.O.D. S.L., S.A., J.W., C.Q., and P.X. designed and performed the mutagenesis study. S.A., M.E., W.L.H., R.O.D., H.A.R., J-P.S., and R.J.L. supervised research. S.A. and R.J.L. coordinated entire research and collaborations. S.L, S.A., P.X., M.E., E.J.F., R.O.D. and R.J.L. wrote the paper with input from all authors.

## References

1 Givertz, M. M. Manipulation of the renin-angiotensin system. Circulation 104, E14–18 (2001). 10.1161/hc3001.094733

2 Lefkowitz, R. J. A brief history of G-protein coupled receptors (Nobel Lecture). Angew Chem Int Ed Engl 52, 6366–6378 (2013). 10.1002/anie.201301924

3 Liu, S., Anderson, P. J., Rajagopal, S., Lefkowitz, R. J. & Rockman, H. A. G Protein-Coupled Receptors: A Century of Research and Discovery. Circ Res 135, 174–197 (2024). 10.1161/CIRCRESAHA.124.323067

4 Rajagopal, K. et al. Beta-arrestin2-mediated inotropic effects of the angiotensin II type 1A receptor in isolated cardiac myocytes. Proc Natl Acad Sci U S A 103, 16284–16289 (2006). 10.1073/pnas.0607583103

5 Kim, K. S. et al. beta-Arrestin-biased AT1R stimulation promotes cell survival during acute cardiac injury. Am J Physiol Heart Circ Physiol 303, H1001–1010 (2012). 10.1152/ajpheart.00475.2012

6 Abraham, D. M. et al. β-Arrestin mediates the Frank–Starling mechanism of cardiac contractility. Proceedings of the National Academy of Sciences 113, 14426–14431 (2016). doi:10.1073/pnas.1609308113

7 Whalen, E. J., Rajagopal, S. & Lefkowitz, R. J. Therapeutic potential of β-arrestin- and G protein-biased agonists. Trends Mol Med 17, 126–139 (2011). 10.1016/j.molmed.2010.11.004

8 Wisler, J. W., Rockman, H. A. & Lefkowitz, R. J. Biased G Protein-Coupled Receptor Signaling: Changing the Paradigm of Drug Discovery. Circulation 137, 2315–2317 (2018). 10.1161/circulationaha.117.028194

9 Nguyen Dinh Cat, A. & Touyz, R. M. Cell signaling of angiotensin II on vascular tone: novel mechanisms. Curr Hypertens Rep 13, 122–128 (2011). 10.1007/s11906-011-0187-x

10 Rajagopal, S. et al. Quantifying ligand bias at seven-transmembrane receptors. Mol Pharmacol 80, 367–377 (2011). 10.1124/mol.111.072801

11 Strachan, R. T. et al. Divergent transducer-specific molecular efficacies generate biased agonism at a G protein-coupled receptor (GPCR). J Biol Chem 289, 14211–14224 (2014). 10.1074/jbc.M114.548131

12 Singh, K. D. & Karnik, S. S. Implications of β-Arrestin biased signaling by angiotensin II type 1 receptor for cardiovascular drug discovery and therapeutics. Cell Signal 124, 111410 (2024). 10.1016/j.cellsig.2024.111410

13 Thal, D. M., Glukhova, A., Sexton, P. M. & Christopoulos, A. Structural insights into G-protein- coupled receptor allostery. Nature 559, 45–53 (2018). 10.1038/s41586-018-0259-z

14 Lorente, J. S. et al. GPCR drug discovery: new agents, targets and indications. Nat Rev Drug Discov 24, 458–479 (2025). 10.1038/s41573-025-01139-y

15 Christopoulos, A. Advances in G protein-coupled receptor allostery: from function to structure. Mol Pharmacol 86, 463–478 (2014). 10.1124/mol.114.094342

16 Grogan, A. et al. A positive allosteric modulator of the beta1AR with antagonist activity for catecholaminergic polymorphic ventricular tachycardia. J Clin Invest 135 (2025). 10.1172/JCI190252

17 Slosky, L. M., Caron, M. G. & Barak, L. S. Biased Allosteric Modulators: New Frontiers in GPCR Drug Discovery. Trends Pharmacol Sci 42, 283–299 (2021). 10.1016/j.tips.2020.12.005

18 Krumm, B. E. et al. Neurotensin Receptor Allosterism Revealed in Complex with a Biased Allosteric Modulator. Biochemistry 62, 1233–1248 (2023). 10.1021/acs.biochem.3c00029

19 Wold, E. A., Chen, J., Cunningham, K. A. & Zhou, J. Allosteric Modulation of Class A GPCRs: Targets, Agents, and Emerging Concepts. J Med Chem 62, 88–127 (2019). 10.1021/acs.jmedchem.8b00875

20 Shen, S. et al. Allosteric modulation of G protein-coupled receptor signaling. Front Endocrinol (Lausanne) 14, 1137604 (2023). 10.3389/fendo.2023.1137604

21 Burford, N. T., Watson, J., Bertekap, R. & Alt, A. Strategies for the identification of allosteric modulators of G-protein-coupled receptors. Biochem Pharmacol 81, 691–702 (2011). 10.1016/j.bcp.2010.12.012

22 Conflitti, P. et al. Functional dynamics of G protein-coupled receptors reveal new routes for drug discovery. Nat Rev Drug Discov 24, 251–275 (2025). 10.1038/s41573-024-01083-3

23 Kenakin, T. P. Biased signalling and allosteric machines: new vistas and challenges for drug discovery. Br J Pharmacol 165, 1659–1669 (2012). 10.1111/j.1476-5381.2011.01749.x

24 Ahn, S. et al. Allosteric “beta-blocker” isolated from a DNA-encoded small molecule library. Proc Natl Acad Sci U S A 114, 1708–1713 (2017). 10.1073/pnas.1620645114

25 Ahn, S. et al. Small-Molecule Positive Allosteric Modulators of the beta(2)-Adrenoceptor Isolated from DNA-Encoded Libraries. Mol Pharmacol 94, 850–861 (2018). 10.1124/mol.118.111948

26 Onaran, H. O. & Costa, T. Where have all the active receptor states gone? Nat Chem Biol 8, 674–677 (2012). 10.1038/nchembio.1024

27 Wingler, L. M. et al. Angiotensin Analogs with Divergent Bias Stabilize Distinct Receptor Conformations. Cell 176, 468–478 e411 (2019). 10.1016/j.cell.2018.12.005

28 Li, A., Liu, S., Huang, R., Ahn, S. & Lefkowitz, R. J. Loss of biased signaling at a G protein- coupled receptor in overexpressed systems. PLoS One 18, e0283477 (2023). 10.1371/journal.pone.0283477

29 Langmead, C. J. Determining allosteric modulator mechanism of action: integration of radioligand binding and functional assay data. Methods Mol Biol 746, 195–209 (2011). 10.1007/978-1-61779-126-0_10

30 Kenakin, T., Jenkinson, S. & Watson, C. Determining the potency and molecular mechanism of action of insurmountable antagonists. J Pharmacol Exp Ther 319, 710–723 (2006). 10.1124/jpet.106.107375

31 Leach, K., Sexton, P. M. & Christopoulos, A. Allosteric GPCR modulators: taking advantage of permissive receptor pharmacology. Trends Pharmacol Sci 28, 382–389 (2007). 10.1016/j.tips.2007.06.004

32 Ehlert, F. J. Analysis of allosterism in functional assays. J Pharmacol Exp Ther 315, 740–754 (2005). 10.1124/jpet.105.090886

33 Kenakin, T. & Strachan, R. T. PAM-Antagonists: A Better Way to Block Pathological Receptor Signaling? Trends Pharmacol Sci 39, 748–765 (2018). 10.1016/j.tips.2018.05.001

34 Zhao, L. H. et al. Conserved class B GPCR activation by a biased intracellular agonist. Nature 621, 635–641 (2023). 10.1038/s41586-023-06467-w

35 Duan, J. et al. GPCR activation and GRK2 assembly by a biased intracellular agonist. Nature 620, 676–681 (2023). 10.1038/s41586-023-06395-9

36 Sun, D. et al. Molecular mechanism of the arrestin-biased agonism of neurotensin receptor 1 by an intracellular allosteric modulator. Cell Res 35, 284–295 (2025). 10.1038/s41422-025-01095-7

37 Shen, S. et al. Structure-based identification of a G protein-biased allosteric modulator of cannabinoid receptor CB1. Proc Natl Acad Sci U S A 121, e2321532121 (2024). 10.1073/pnas.2321532121

38 Sun, Q. et al. A pan-positive allosteric modulator that mediates sustainable GPCR activation. bioRxiv, 2025.2003.2024.645127 (2025). 10.1101/2025.03.24.645127

39 Kobayashi, K. et al. Class B1 GPCR activation by an intracellular agonist. Nature 618, 1085–1093 (2023). 10.1038/s41586-023-06169-3

40 Wingler, L. M., McMahon, C., Staus, D. P., Lefkowitz, R. J. & Kruse, A. C. Distinctive Activation Mechanism for Angiotensin Receptor Revealed by a Synthetic Nanobody. Cell 176, 479–490 e412 (2019). 10.1016/j.cell.2018.12.006

41 Wingler, L. M. et al. Angiotensin and biased analogs induce structurally distinct active conformations within a GPCR. Science 367, 888–892 (2020). 10.1126/science.aay9813

42 Wu, X. & Rapoport, T. A. Cryo-EM structure determination of small proteins by nanobody- binding scaffolds (Legobodies). Proc Natl Acad Sci U S A 118 (2021). 10.1073/pnas.2115001118

43 Elgeti, M. & Hubbell, W. L. DEER Analysis of GPCR Conformational Heterogeneity. Biomolecules 11 (2021). 10.3390/biom11060778

44 Suomivuori, C. M. et al. Molecular mechanism of biased signaling in a prototypical G protein- coupled receptor. Science 367, 881–887 (2020). 10.1126/science.aaz0326

45 Zhang, D. et al. Structural insights into angiotensin receptor signaling modulation by balanced and biased agonists. EMBO J 42, e112940 (2023). 10.15252/embj.2022112940

46 Shukla, A. K. et al. Visualization of arrestin recruitment by a G-protein-coupled receptor. Nature 512, 218–222 (2014). 10.1038/nature13430

47 Latorraca, N. R. et al. Molecular mechanism of GPCR-mediated arrestin activation. Nature 557, 452–456 (2018). 10.1038/s41586-018-0077-3

48 Sutkeviciute, I. & Vilardaga, J. P. Structural insights into emergent signaling modes of G protein- coupled receptors. J Biol Chem 295, 11626–11642 (2020). 10.1074/jbc.REV120.009348

49 Olsen, R. H. J. et al. TRUPATH, an open-source biosensor platform for interrogating the GPCR transducerome. Nat Chem Biol 16, 841–849 (2020). 10.1038/s41589-020-0535-8

50 Onaran, H. O. & Costa, T. Allosteric coupling and conformational fluctuations in proteins. Curr Protein Pept Sci 10, 110–115 (2009). 10.2174/138920309787847644

51 Khanna, A. et al. Angiotensin II for the Treatment of Vasodilatory Shock. N Engl J Med 377, 419–430 (2017). 10.1056/NEJMoa1704154

52 Ahn, S., Kim, J., Hara, M. R., Ren, X. R. & Lefkowitz, R. J. beta-Arrestin-2 Mediates Anti- apoptotic Signaling through Regulation of BAD Phosphorylation. J Biol Chem 284, 8855–8865 (2009). 10.1074/jbc.M808463200

53 Sanchez-Garcia, R. et al. DeepEMhancer: a deep learning solution for cryo-EM volume post- processing. Commun Biol 4, 874 (2021). 10.1038/s42003-021-02399-1

54 Reeves, P. J., Kim, J. M. & Khorana, H. G. Structure and function in rhodopsin: a tetracycline- inducible system in stable mammalian cell lines for high-level expression of opsin mutants. Proc Natl Acad Sci U S A 99, 13413–13418 (2002). 10.1073/pnas.212519199

55 Staus, D. P. et al. Structure of the M2 muscarinic receptor-beta-arrestin complex in a lipid nanodisc. Nature 579, 297–302 (2020). 10.1038/s41586-020-1954-0

56 Dahl, L. et al. Multiplexed selectivity screening of anti-GPCR antibodies. Sci Adv 9, eadf9297 (2023). 10.1126/sciadv.adf9297

57 Hauge Pedersen, M., et al. A novel luminescence-based beta-arrestin recruitment assay for unmodified receptors. J Biol Chem 296, 100503 (2021). 10.1016/j.jbc.2021.100503

58 Martini, J. S. et al. Uncovering G protein-coupled receptor kinase-5 as a histone deacetylase kinase in the nucleus of cardiomyocytes. Proc Natl Acad Sci U S A 105, 12457–12462 (2008). 10.1073/pnas.0803153105

59 Fan, L. et al. Structural basis of psychedelic LSD recognition at dopamine D(1) receptor. Neuron 112, 3295–3310 e3298 (2024). 10.1016/j.neuron.2024.07.003

60 Duan, J. et al. Cryo-EM structure of an activated VIP1 receptor-G protein complex revealed by a NanoBiT tethering strategy. Nat Commun 11, 4121 (2020). 10.1038/s41467-020-17933-8

61 Mao, C. et al. Unsaturated bond recognition leads to biased signal in a fatty acid receptor. Science 380, eadd6220 (2023). 10.1126/science.add6220

62 Cheng, Y. H., Huang, X. J., Xu, B. & Ding, W.: automatic data transfer, preprocessing, real-time display and monitoring in cryo-EM. Journal of Applied Crystallography 56, 1865–1873 (2023). 10.1107/S1600576723008257

63 Punjani, A., Rubinstein, J. L., Fleet, D. J. & Brubaker, M. A. cryoSPARC: algorithms for rapid unsupervised cryo-EM structure determination. Nat Methods 14, 290–296 (2017). 10.1038/nmeth.4169

64 Croll, T. I. ISOLDE: a physically realistic environment for model building into low-resolution electron-density maps. Acta Crystallogr D Struct Biol 74, 519–530 (2018). 10.1107/s2059798318002425

65 Emsley, P., Lohkamp, B., Scott, W. G. & Cowtan, K. Features and development of Coot. Acta Crystallogr D Biol Crystallogr 66, 486–501 (2010). 10.1107/S0907444910007493

66 Jumper, J. et al. Highly accurate protein structure prediction with AlphaFold. Nature 596, 583–589 (2021). 10.1038/s41586-021-03819-2

67 Pettersen, E. F. et al. UCSF Chimera--a visualization system for exploratory research and analysis. J Comput Chem 25, 1605–1612 (2004). 10.1002/jcc.20084

68 Adams, P. D. et al. PHENIX: a comprehensive Python-based system for macromolecular structure solution. Acta Crystallogr D Biol Crystallogr 66, 213–221 (2010). 10.1107/S0907444909052925

69 Pettersen, E. F. et al. UCSF ChimeraX: Structure visualization for researchers, educators, and developers. Protein Sci 30, 70–82 (2021). 10.1002/pro.3943

70 Fabregas Ibanez, L., Jeschke, G. & Stoll, S. DeerLab: a comprehensive software package for analyzing dipolar electron paramagnetic resonance spectroscopy data. Magn Reson (Gott) 1, 209–224 (2020). 10.5194/mr-1-209-2020

71 Tessmer, M. H. & Stoll, S. chiLife: An open-source Python package for in silico spin labeling and integrative protein modeling. PLoS Comput Biol 19, e1010834 (2023). 10.1371/journal.pcbi.1010834

72 Zhang, H. et al. Structure of the Angiotensin receptor revealed by serial femtosecond crystallography. Cell 161, 833–844 (2015). 10.1016/j.cell.2015.04.011

73 Lomize, M. A., Lomize, A. L., Pogozheva, I. D. & Mosberg, H. I. OPM: orientations of proteins in membranes database. Bioinformatics 22, 623–625 (2006). 10.1093/bioinformatics/btk023

