## Extended Data for "Conformational Basis of Functionally Selective Allosteric Modulation of the Angiotensin II type 1 Receptor by Small Molecules"

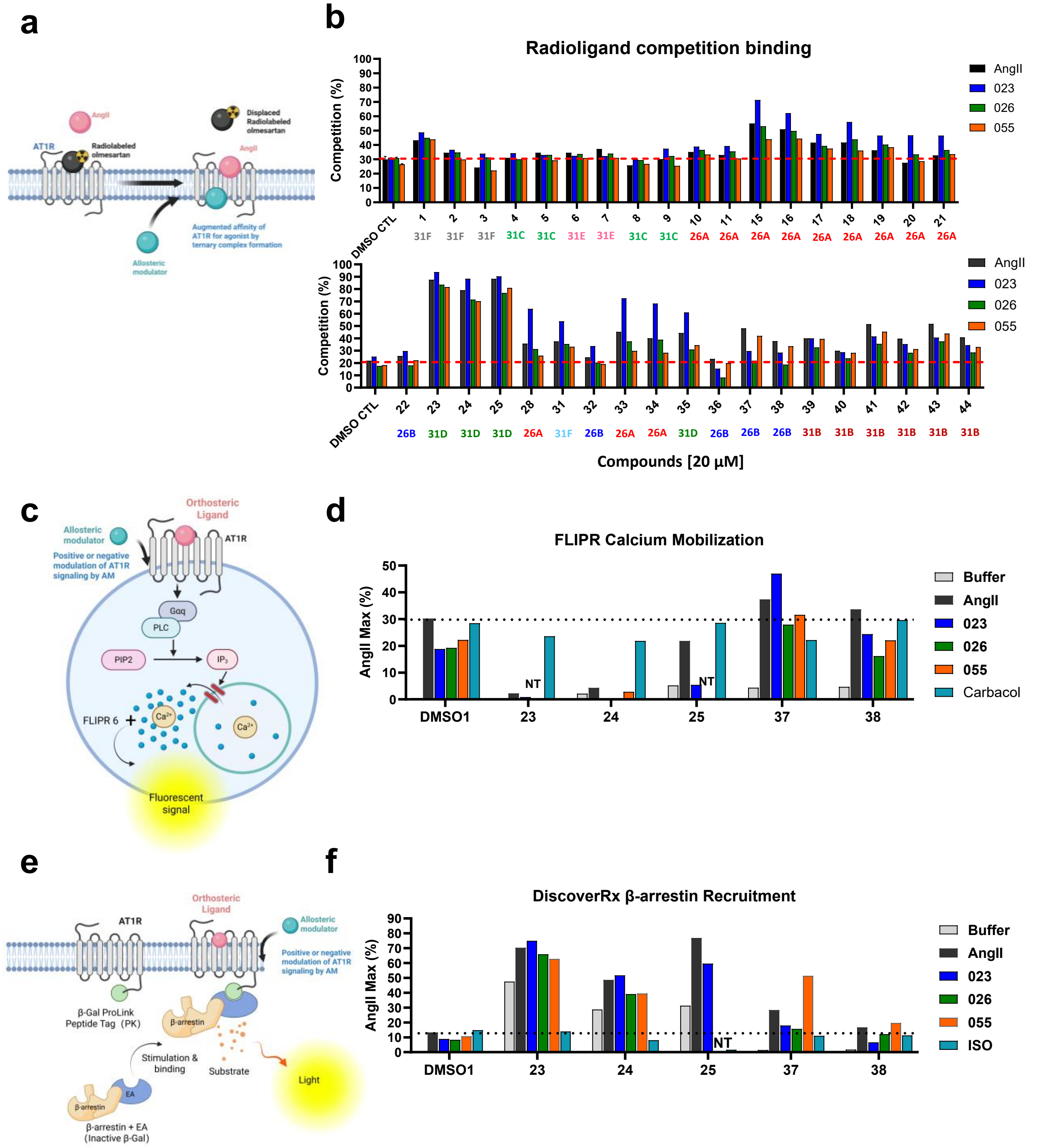


**Extended Data Fig. 1. Initial activity screening with the potential binders through binding and functional activity assays.**

Thirty-seven putative binders from the DEL screen were selected, re-synthesized on a small scale without DNA-tags and tested for their allosteric activities.

(**a**) An illustration for radioligand orthosteric agonist competition binding, created in https://BioRender.com.

(**b**) [$^3$H]-olmesartan was incubated with each of the orthosteric peptide agonists (AngII, TRV023, TRV026 and TRV055) as indicated in the presence of one of the 37 compounds at 20 µM. Dimethyl

sulfoxide (DMSO), vehicle-treated conditions with each of the peptide agonists were utilized as controls. Values were expressed as % competition (% decreases in [$^3$H]-olmesartan binding) over the maximum radiolabeled tracer binding in the absence of an agonist in the vehicle-treated condition. The data was obtained from a single or two (Mean values) independent experiments depending on amounts of synthesized compounds. The 37 compounds were categorized based on their chemical composition similarities. Compounds belonging to the same group are indicated with the same-colored group name.
(**c**) An illustration for AT1R-mediated downstream, Gq protein-dependent intracellular calcium response that was monitored by the FLIPR6 calcium mobilization assay, created in https://BioRender.com.
(**d**) HEK-293 cells stably expressing the AT1R were incubated with one of the indicated compounds at 20 μM for 20 min prior to stimulation with one of the agonists (AngII, TRV023, TRV026 and TRV055) at its around $EC_{25}$ concentrations. Upon agonist stimulation, calcium mobilization was monitored for 2.5 min as described in 'Methods'. The non-specific effect of the selected compounds in a counter assay, in which carbachol-stimulated calcium mobilization in cells expressing endogenous muscarinic acetylcholine receptors, was also monitored.
(**e**) An illustration for the agonist-induced β-arrestin recruitment to the AT1R that was assessed by the PathHunter enzyme complementation assay, created in https://BioRender.com.
(**f**) After pretreatment with one of the selected compounds at 20 μM for 20 min, cells stably expressing AT1R-pk and EA-β-arrestin2 molecules were stimulated with one of the indicated agonists at their around $EC_{10}$ concentration for 30 min. In both assays shown in 'd' and 'f', vehicle (0.2% DMSO)-treated conditions with each of the peptide agonists were utilized as controls. Values were expressed as percentage over the maximum AngII-stimulated response in the vehicle-treated condition obtained from a single experiment. As a counter assay, isoproterenol-induced β-arrestin recruitment to the chimeric β2V2 receptor (β2V2R) in the presence of one of the selected compounds was monitored in cells expressing β2V2R-pk and EA-β-arrestin2. Note that due to the very small amounts of the re-synthesized putative AT1R binders, we performed limited replicates of these initial assays.

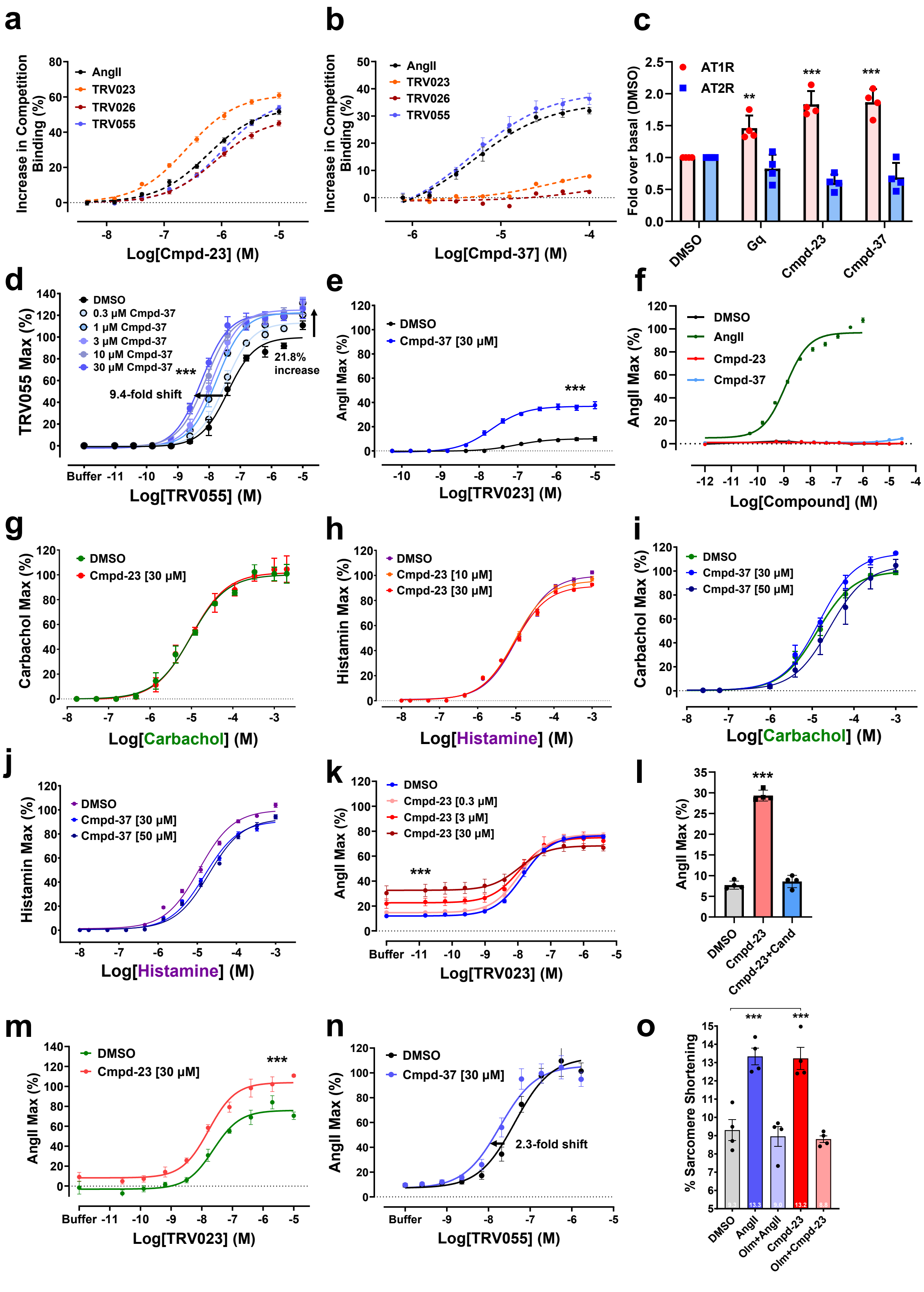
a
Increase in Competition Binding (%)
AngII
TRV023
TRV026
TRV055
Log[Cmpd-23] (M)
b
Increase in Competition Binding (%)
AngII
TRV023
TRV026
TRV055
Log[Cmpd-37] (M)
c
Fold over basal (DMSO)
AT1R
AT2R
**
***
***
DMSO
Gq
Cmpd-23
Cmpd-37
d
TRV055 Max (%)
DMSO
0.3 μM Cmpd-37
1 μM Cmpd-37
3 μM Cmpd-37
10 μM Cmpd-37
30 μM Cmpd-37
***
9.4-fold shift
21.8% increase
Log[TRV055] (M)
e
AngII Max (%)
DMSO
Cmpd-37 [30 μM]
***
Log[TRV023] (M)
f
AngII Max (%)
DMSO
AngII
Cmpd-23
Cmpd-37
Log[Compound] (M)
g
Carbachol Max (%)
DMSO
Cmpd-23 [30 μM]
Log[Carbachol] (M)
h
Histamin Max (%)
DMSO
Cmpd-23 [10 μM]
Cmpd-23 [30 μM]
Log[Histamine] (M)
i
Carbachol Max (%)
DMSO
Cmpd-37 [30 μM]
Cmpd-37 [50 μM]
Log[Carbachol] (M)
j
Histamin Max (%)
DMSO
Cmpd-37 [30 μM]
Cmpd-37 [50 μM]
Log[Histamine] (M)
k
AngII Max (%)
DMSO
Cmpd-23 [0.3 μM]
Cmpd-23 [3 μM]
Cmpd-23 [30 μM]
***
Log[TRV023] (M)
l
AngII Max (%)
***
DMSO
Cmpd-23
Cmpd-23+Cand
m
AngII Max (%)
DMSO
Cmpd-23 [30 μM]
***
Log[TRV023] (M)
n
AngII Max (%)
DMSO
Cmpd-37 [30 μM]
2.3-fold shift
Log[TRV055] (M)
o
% Sarcomere Shortening
***
***
DMSO
AngII
Olm+AngII
Cmpd-23
Olm+Cmpd-23

**Extended Data Fig. 2. Additional pharmacological profiles with Cmpd-23 and Cmpd-37 in radioligand agonist competition binding assays.**

(**a,b**) Increases in the extent of competition binding of each indicated peptide orthosteric agonist against [$^3$H]-Olmesartan binding to the AT1R in the presence of (a) Cmpd-23 and (b) Cmpd-37 were monitored in a dose-dependent manner. Each of the agonists was utilized at approximately their IC20 concentrations. Values were expressed as percentages of increases in the extent of agonist competition against [$^3$H]-Olmesartan binding to the receptor by the allosteric compounds at their indicated concentrations over vehicle (dimethylsulfoxide; DMSO)-treated.

(**c**) [$^3$H]-angiotensin II (AngII) binding to the membranes isolated from either AT1R- or AT2R-expressing Expi293 cells. [$^3$H]-AngII at ~10 nM was used in the presence of 50 μM allosteric compounds. Specific cpm, calculated by subtraction of non-specific cpm obtained in the presence of 20 μM of Olmesartan (for the AT1R) and 10 μM CGP-42133 (for the AT2R), was normalized as fold-increases over DMSO-treated controls.

(**d**) Effects of Cmpd-37 on TRV055-stimulated Gq activation. G protein biased TRV055-stimulated Gq-mediated signaling in the presence of Cmpd-37 at its various concentrations as indicated was monitored by the FLIPR-6 intracellular calcium mobilization assay as described in 'Methods'. Each data point was normalized to the maximal level of the TRV055-induced activity the vehicle (0.3% DMSO) control and expressed as a percentage. The shift of curves was expressed as fold changes in EC50 and % alterations in Emax values.

(**e**) Cmpd-37 robustly increases the minimal calcium mobilization response to stimulation with TRV023. HEK293 cells stably transfected with a rat AT1R expression plasmid were incubated with either 0.3 % DMSO vehicle control or Cmpd-37 at 30 μM for ~20 min. Upon stimulation with TRV023 in a dose-dependent manner, the real time changes in the intracellular calcium level were monitored. Each data point was normalized to the maximal level of the AngII-induced activity in vehicle control and expressed as a percentage.

(**f**) No intrinsic agonistic activity of Cmpd-23 and Cmpd-37 in calcium signaling. U2OS cells stably expressing the AT1R were stimulated with AngII, Cmpd-23 and Cmpd-37 in a dose-dependent manner. Upon stimulation the real time changes in the intracellular calcium level were monitored as described in 'Methods'. Each data point was normalized to the maximal level of the AngII-stimulated activity and expressed as a percentage.

(**g-j**) Counter activity assays. U2OS cells endogenously expressing muscarinic acetylcholine and histamine H1 receptors were incubated with either (g,h) Cmpd-23 at 10 and/or 30 μM or (i,j) Cmpd-37 at 30 and 50 μM as indicated for 15-30 min prior to stimulation with either (g,i) carbachol or (h,j) histamine in a dose-dependent manner. Upon stimulation, intracellular calcium levels were monitored. Areas under the curve (AUC) were calculated at different concentrations of carbachol or histamine to obtain dose response curves. Each data point was normalized to the maximal level of the carbachol- or histamine-induced activity in the vehicle (0.3% dimethylsulfoxide; DMSO) control and expressed as a percentage.

(**k**) The effect of Cmpd-23 on TRV023-stimulated β-arrestin recruitment to the AT1R. After around 15 min incubation with either 0.3% DMSO vehicle control or Cmpd-23 at the indicated concentrations, the extent of TRV-23-stimulated β-arrestin recruitment to the AT1R was measured as described in 'Methods'. Values were expressed as percentage over the maximum AngII-stimulated response in the vehicle-treated condition.

(**l**) Authenticity of Cmpd-23 allosteric agonism in β-arrestin recruitment to the AT1R. Cmpd-23 stimulated β-arrestin recruitment to the AT1R was measured in the absence or presence of the orthosteric antagonist candesartan at 10 μM (10 min pre-incubation). Each bar is expressed as percentage over the maximal level of the AngII-induced activity in vehicle control.

(**m**) The PAM activity of Cmpd-23 in TRV023-stimulated β-arrestin internalization leading to endosomal localization. Cells were pre-incubated with either 0.3% DMSO vehicle control or Cmpd-23 at 30 μM for ~15 min and stimulated with TRV023 in a dose-dependent fashion. β-arrestin internalization was measured as described in 'Methods'. Values were expressed as percentage over the maximum AngII-stimulated response in the vehicle-treated condition.

(**n**) Effects of Cmpd-37 on TRV055-stimulated β-arrestin recruitment. TRV055-induced β-arrestin recruitment to the AT1R in the presence of Cmpd-37 at 30 μM was determined by the PathHunter enzyme complementation assay. Values were expressed as percentages over the maximum agonist-stimulated response in the vehicle-treated condition.

(**o**) Cardiomyocyte contractility was measured as sarcomere shortening after stimulation with either AngII at 10 μM or Cmpd-23 at 25 μM in the presence of either vehicle (DMSO) control or 10 μM Olmesartan (Olm). Values shown are expressed as percentage of sarcomere shortening from 4 wild-type hearts with 8-12 cells analyzed per condition per heart. All the data points in this figure represent mean ± SEM.

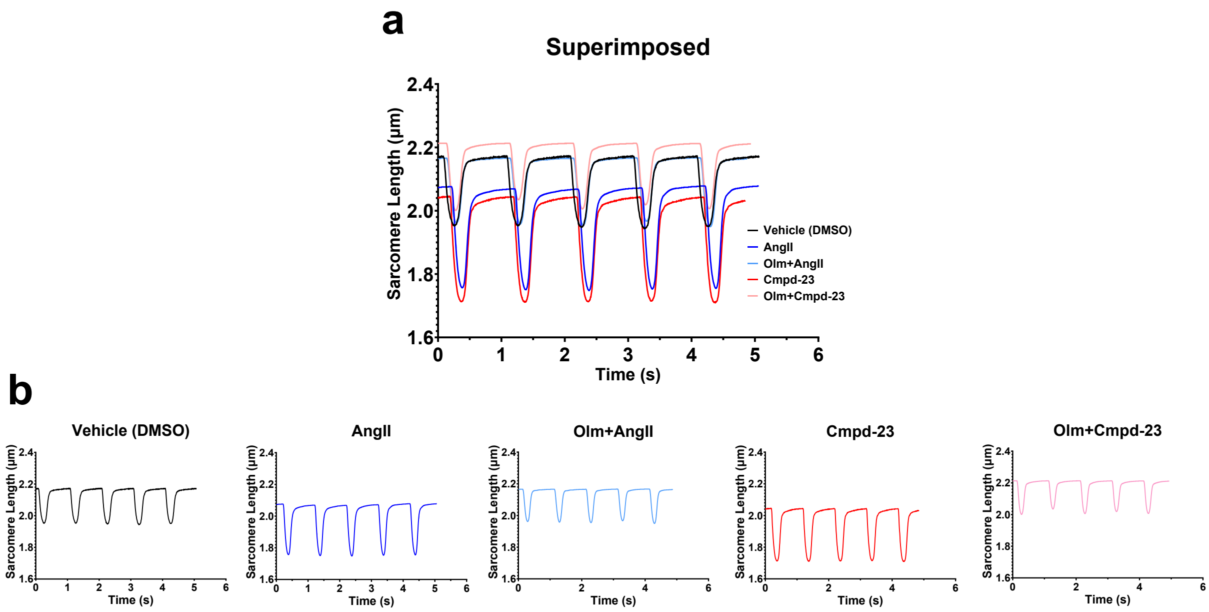


**Extended Data Fig. 3. Representative traces from cardiomyocyte contraction recordings.**
(**a**) Superimposed representative traces from individual conditions.
(**b**) Representative traces from individual conditions as indicated.

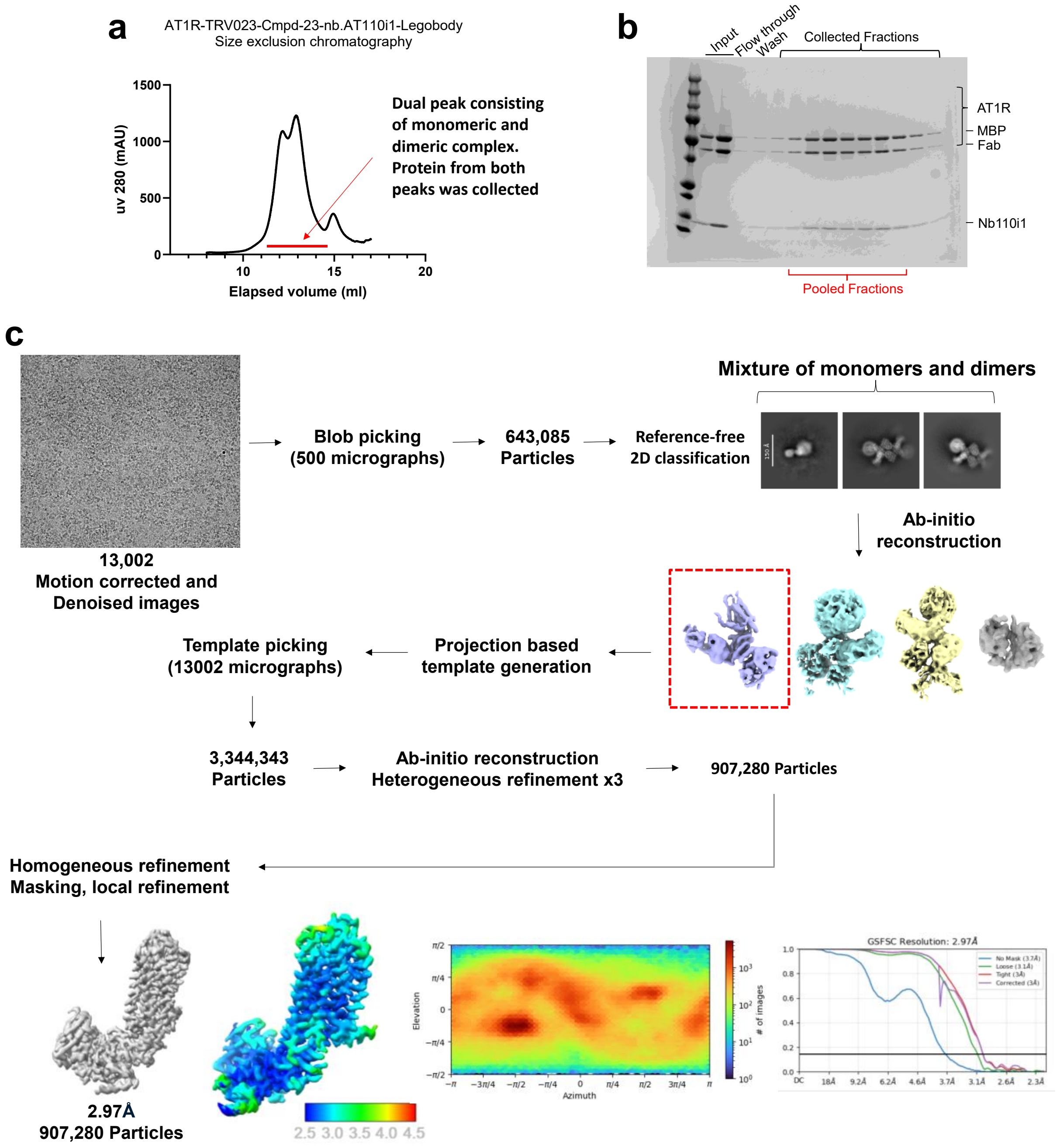


**Extended Data Fig. 4. Purification and cryo-EM structural determination of AT1R-TRV023-Cmpd-23-Nb.AT110i1-Legobody complex.**

**(a)** SEC profile of the final AT1R-TRV023-Cmpd-23-NbAT110i1-legobody complex.

**(b)** Analysis of complex formation in the SEC fractions by SDS-PAGE. The first and the second lanes of 'Input' are samples before and after the concentration process, respectively for SEC loading.

**(c)** Schematic representation of cryo-EM data processing pipeline for the complex. Gold standard Fourier Shell Correlation (GSFSC) plots of the focused refinement reconstruction for the AT1R-TRV023-Cmpd-23-NbAT110i1-legobody complex. A threshold of 0.143 was used to determine the overall resolution of the map.

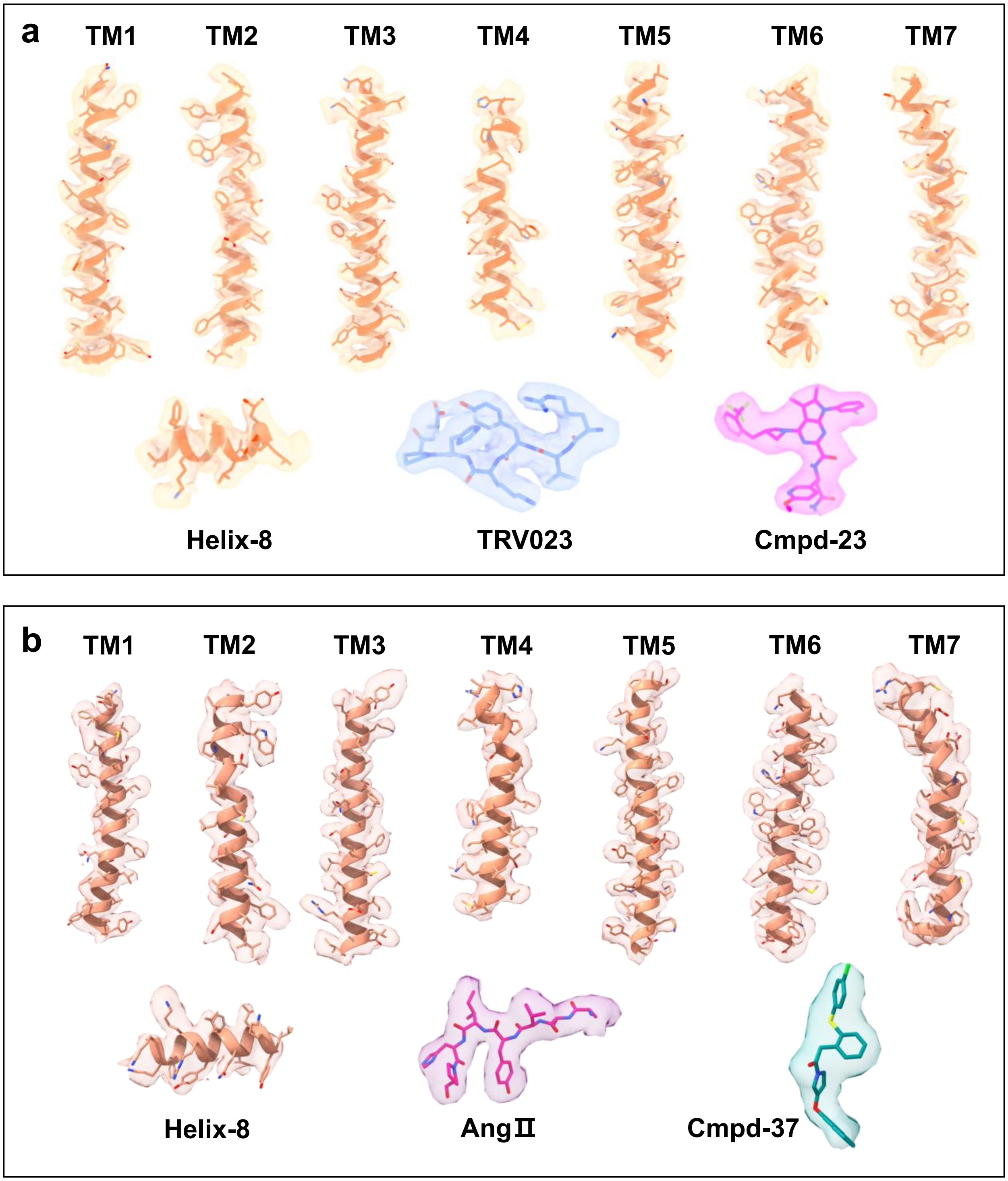


**Extended Data Fig. 5. (a) Model of Cmpd-23-AT1R-TRV023 and (b) Cmpd-37-AT1R-AngII in electron density maps.**

EM densities for TM1-TM7 and helix 8 of the AT1R, the orthosteric ligands (a) TRV023 and (b) AngII, and the allosteric ligands (a) Cmpd-23 and (b) Cmpd-37.

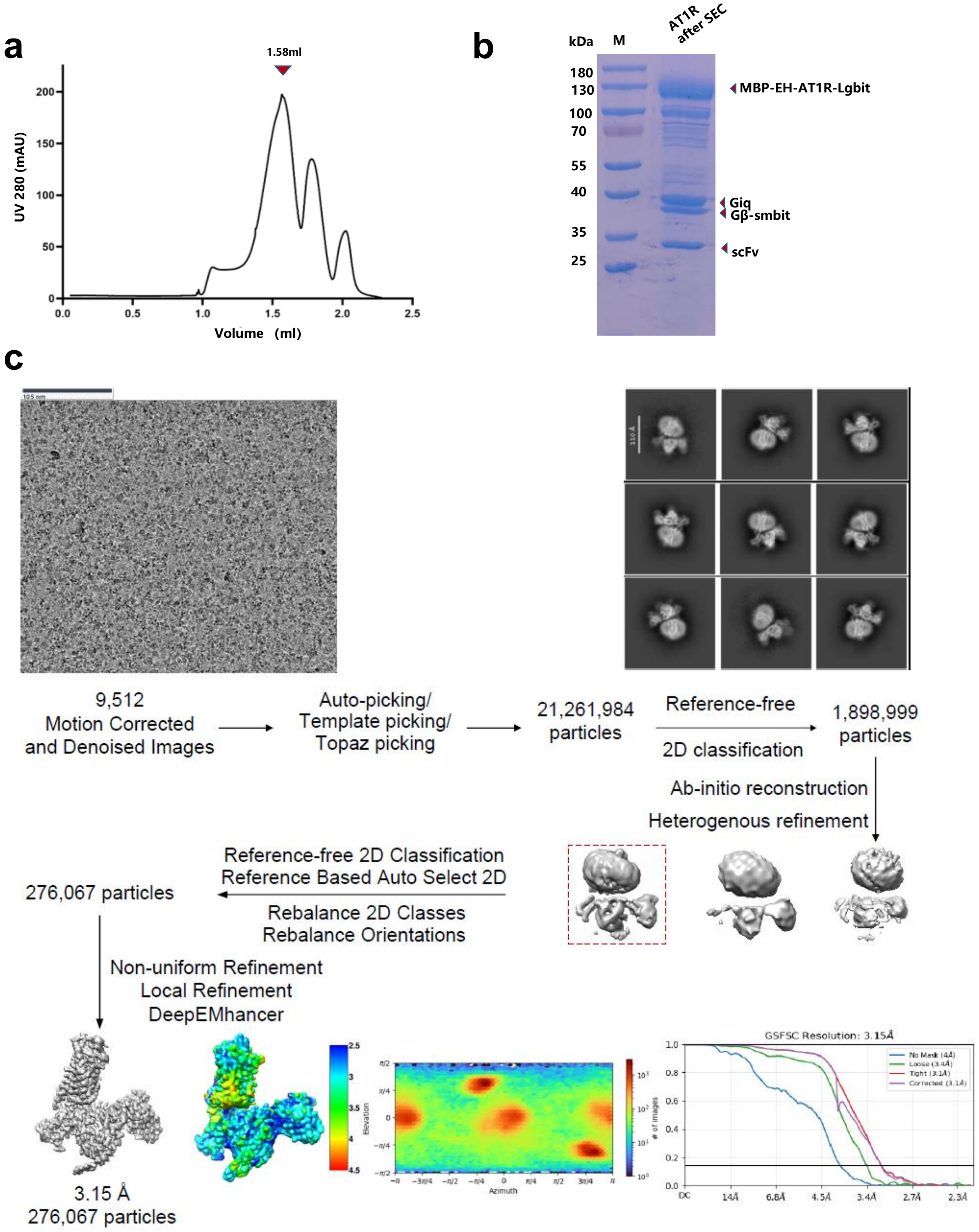


**Extended Data Fig. 6. Purification and cryo-EM structural determination of AT1R-Gq-AngII-Cmpd-37 complex.**

(**a**) SEC of the final AT1R-Giq-AngII-Cmpd-37 complex.

(**b**) Analysis of complex formation in the final fraction applied to the grid by SDS page.

(**c**) Schematic representation of cryo-EM data processing pipeline for the complex. Gold standard Fourier Shell Correlation (GSFSC) plots of the focused refinement reconstruction for the AT1R-Giq-AngII-Cmpd-37 complex. A threshold of 0.143 was used to determine the overall resolution of the map.

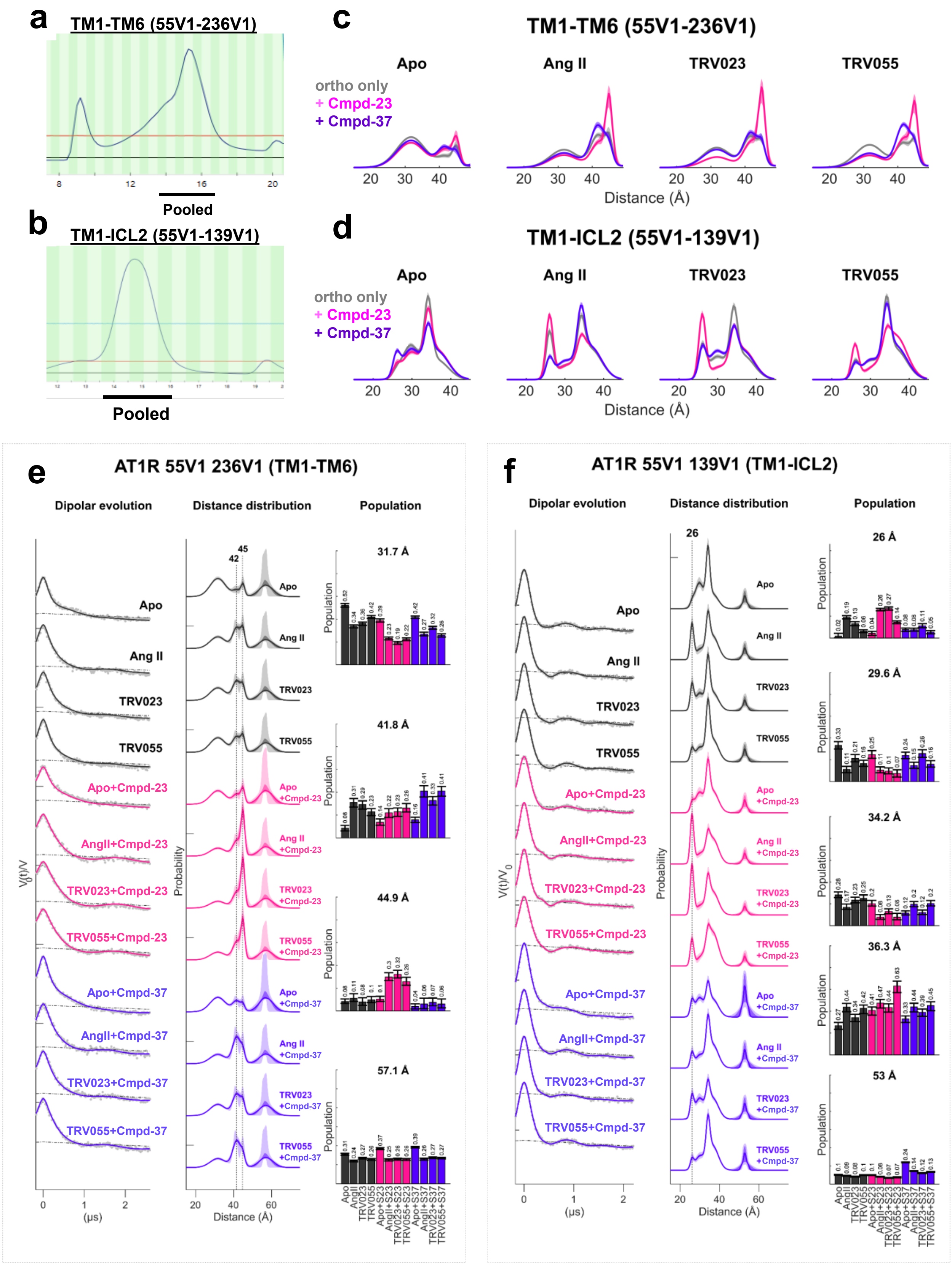

a
TM1-TM6 (55V1-236V1)
Pooled
b
TM1-ICL2 (55V1-139V1)
Pooled
c
TM1-TM6 (55V1-236V1)
Apo
Ang II
TRV023
TRV055
ortho only
+ Cmpd-23
+ Cmpd-37
Distance (Å)
d
TM1-ICL2 (55V1-139V1)
Apo
Ang II
TRV023
TRV055
ortho only
+ Cmpd-23
+ Cmpd-37
Distance (Å)
e
AT1R 55V1 236V1 (TM1-TM6)
Dipolar evolution
Distance distribution
Population
31.7 Å
41.8 Å
44.9 Å
57.1 Å
Apo
Ang II
TRV023
TRV055
Apo+Cmpd-23
AngII+Cmpd-23
TRV023+Cmpd-23
TRV055+Cmpd-23
Apo+Cmpd-37
AngII+Cmpd-37
TRV023+Cmpd-37
TRV055+Cmpd-37
(μs)
Distance (Å)
Probability
Population
f
AT1R 55V1 139V1 (TM1-ICL2)
Dipolar evolution
Distance distribution
Population
26 Å
29.6 Å
34.2 Å
36.3 Å
53 Å
Apo
Ang II
TRV023
TRV055
Apo+Cmpd-23
AngII+Cmpd-23
TRV023+Cmpd-23
TRV055+Cmpd-23
Apo+Cmpd-37
AngII+Cmpd-37
TRV023+Cmpd-37
TRV055+Cmpd-37
(μs)
Distance (Å)
Probability
Population

**Extended Data Fig. 7. Sample Preparation for DEER spectroscopy and DEER profiles with Cmpd-23 and Cmpd-37 across three orthosteric peptide agonists and allosteric ligands alone.**

(**a**) SEC profile of the purified AT1R with the spin labeled Cys 55(TM1) - 236(TM6) pair used for DEER spectroscopy.

(**b**) SEC profile of the purified AT1R with the spin labeled Cys 55(TM1) - 139(ICL2) pair used for DEER spectroscopy.

(**c**) All DEER profiles for TM1-TM6 distances of the AT1R. Data is split into panels by orthosteric site occupancy—Apo (no ligand), AngII, TRV023 and TRV055. Within each panel, distance traces are distinguished by allosteric modulators (AMs) with different colors (no AM, grey; Cmpd-23, pink; Cmpd-37, purple).

(**d**) All DEER profiles for TM1-ICL2 distances of the AT1R. Data is presented as described above for ‘c’.

**(e)** 55V1-236V1 (TM1-TM6). Global fit of all dipolar evolutions to a Gaussian mixture model reveals four conformations, one (31.7 Å) comprising inactive AT1R conformations, two active AT1R conformations around 42 Å and 45 Å which are differentially modulated by Cmpd-23 and Cmpd-37, and one around 57 Å indicating aggregation. Shaded areas indicate 50% and 95% confidence intervals, respectively. The population analysis reveals specific conformational shifts due to binding of orthosteric and allosteric ligands.

(**f**) 55V1-139V1 (TM1-ICL2). Global fit of all dipolar evolutions to a Gaussian mixture model reveals five conformations, including the 26 Å distance peak indicative of the Aoccl2 conformation. The distance at 53 Å indicates aggregated AT1R. Shaded areas indicate 50% and 95% confidence intervals, respectively. The population analysis reveals significant stabilization of the 26 Å by Cmpd-23 and destabilization by Cmpd-37.

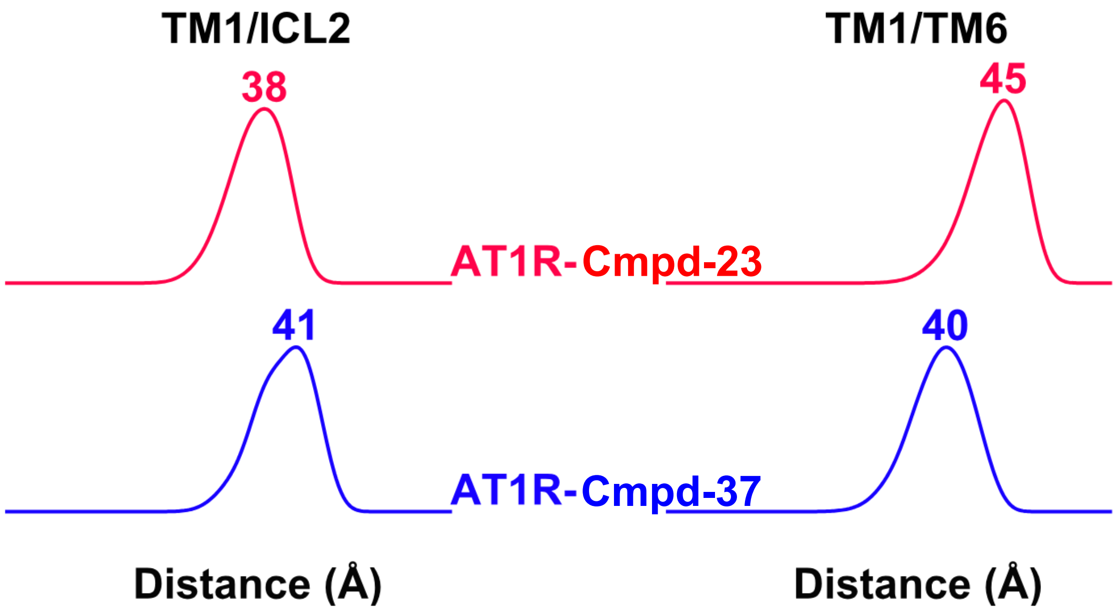


**Extended Data Fig. 8. Simulated DEER distances based on high-resolution structures.**

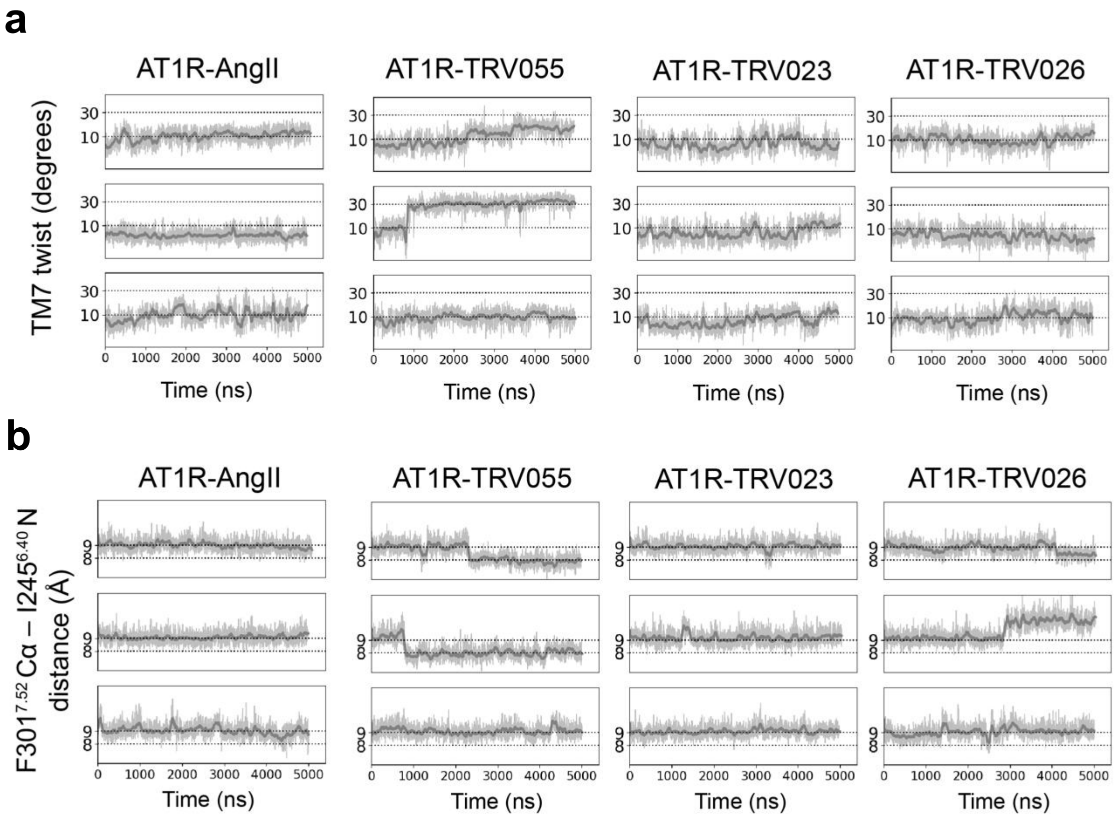


**Extended Data Fig. 9. The TM6–TM7 distance decreases with TM7 twist.**

**(a)** Simulation traces showing the TM7 twist measurement (see 'Methods') in all AT1R-AngII, AT1R-TRV055, AT1R-TRV023, and AT1R-TRV026 simulations (going from left to right). Each simulation was started from 6DO1 with the respective ligand modeled in as described in 'Methods'. Smoothed traces are dark grey (smoothing window of 30 ns), unsmoothed traces are in light grey. Dashed lines at 10° and 30° are shown for reference.

(**b**) Simulation traces showing the F3017.52 Cα – I2456.40 N measurement in all AT1R-AngII, AT1R-TRV055, AT1R-TRV023, and AT1R-TRV026 simulations (going from left to right). Each simulation was started from 6DO1 with the respective ligand modeled in as described in 'Methods'. Smoothed traces are dark grey (smoothing window of 30 ns), unsmoothed traces are in light grey. Dashed lines at 8 Å and 9 Å are shown for reference.

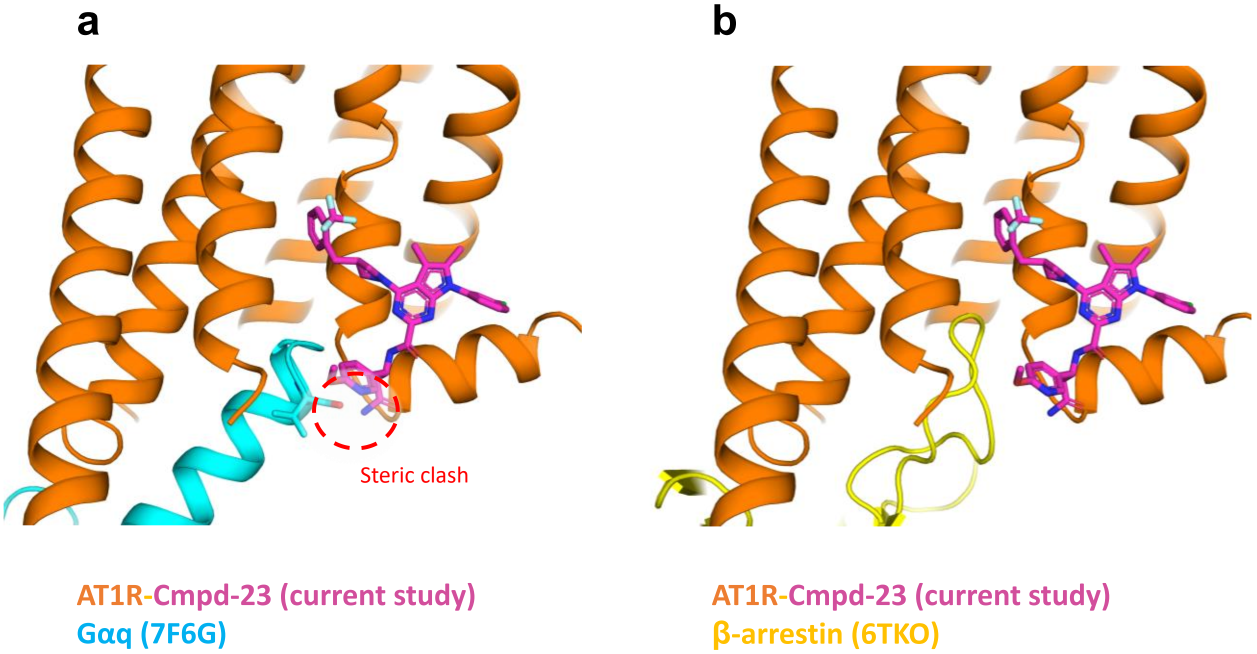


**Extended Data Fig. 10. Cmpd-23 sterically interferes with Gaq, but not β-arrestin coupling to the AT1R.**

Structural models of Gaq (light blue) and β-arrestin (yellow) coupling into the Cmpd-23-occupied AT1R structure observed in this study.

(**a**) Cmpd-23-occupied AT1R structure superimposed with Gαq transducer from an AT1R-Gq complex structure (7F6G). The region of steric clash of Gαq with Cmpd-23 circled in red.

(**b**) Cmpd-23-occuped AT1R structure superimposed with the β-arrestin from a β1AR-β-arrestin complex structure (6TKO). Structural models were generated using the matchmaker tool in ChimeraX based on alignment of the Cmpd-23 bound AT1R to the transducer bound receptor structures.

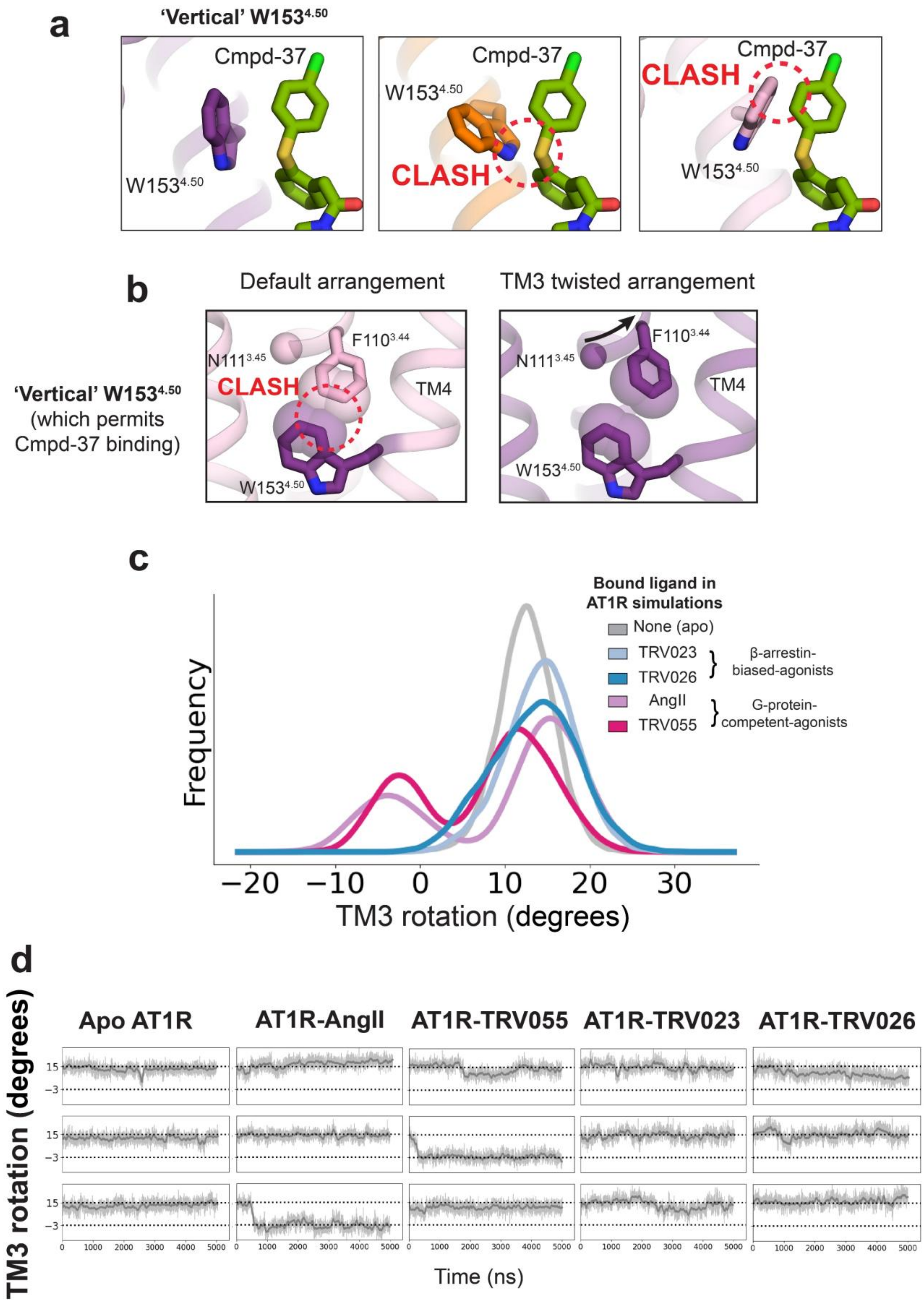
a
'Vertical' W153$^{4.50}$
Cmpd-37
W153$^{4.50}$
CLASH
b
Default arrangement
TM3 twisted arrangement
'Vertical' W153$^{4.50}$
(which permits
Cmpd-37 binding)
N111$^{3.45}$
F110$^{3.44}$
TM4
c
Bound ligand in
AT1R simulations
None (apo)
TRV023
TRV026
β-arrestin-
biased-agonists
AngII
TRV055
G-protein-
competent-agonists
Frequency
TM3 rotation (degrees)
d
Apo AT1R
AT1R-AngII
AT1R-TRV055
AT1R-TRV023
AT1R-TRV026
TM3 rotation (degrees)
Time (ns)

**Extended Data Fig. 11. Cmpd-23 binding is permitted when L112$^{3.36}$ rotates away from the receptor core**

(**a**) Renderings showing Cmpd-37 (green) as modeled in the cryo-EM structure (left, purple), and overlaid with two other AT1R structures (orange, PDB: 6OS1 and pink, PDB: 6DO1) with varying rotamers of W153$^{4.50}$. These renderings show a clash between Cmpd-37 and W153$^{4.50}$ in the two rightmost renderings, demonstrating how Cmpd-37 binding is only accommodated by the 'vertical' rotamer of W153$^{4.50}$ (left). Structures were aligned on all Cα atoms of AT1R TM1-4.

(**b**) Renderings showing the 'vertical' rotamer of W153$^{4.50}$ in structures with default (left, purple) and TM3 twisted (right, pink) arrangements. To make the rendering on the right, the rotamer of W153$^{4.50}$ was modified to be the 'vertical' rotamer using PyMOL's mutagenesis tool. The 'vertical' rotamer of W153$^{4.50}$ can be adopted when TM3 is in the twisted arrangement, not the default arrangement; otherwise, F110$^{3.34}$ would clash with W153$^{4.50}$ (as indicated by overlapping Van der Waals radii). As such, Cmpd-37 binding, which requires adoption of the 'vertical' W153$^{4.50}$ rotamer, is better accommodated by the TM3 twisted arrangement.

**(c)** Distributions of the TM3 rotation measurement across apo (grey), TRV023-bound (light blue), TRV026-bound (dark blue), AngII-bound (light pink), and TRV055-bound (dark pink) AT1R simulations.

**(d)** Simulation traces showing the TM3 rotation measurement in apo (unliganded), AngII-bound, TRV023-bound, TRV026-bound, and TRV55-bound AT1R simulations (see Methods).

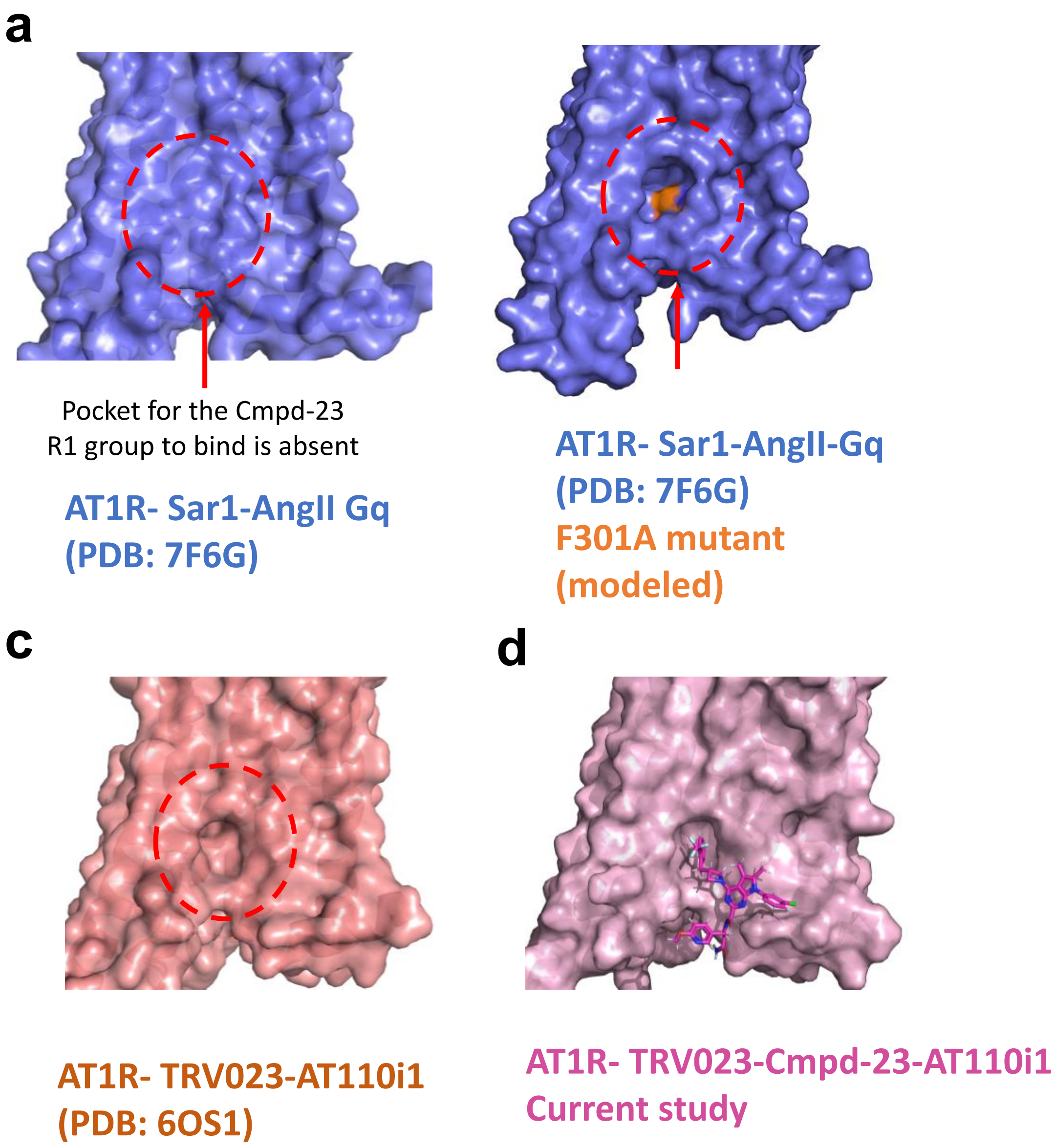


**Extended Data Fig. 12. Cmpd-23 binding region is altered in the F301A mutant, allowing for the possibility of Cmpd-23 binding to the canonical active state.**

**(a)** Binding pocket for the trifluoromethylbenzyl group (designated as R1; Fig. 3d) of Cmpd-23 (circled in red) is absent in the canonical active state

**(b)** Modeled structure of the F301A mutant AT1R showing a suitable pocket for R1 group insertion, which is re-introduced with the F301A mutation. The remodeled surface introduced by the F301A mutation is shown in orange. Receptor surface models are shown as solvent-excluded surfaces.

(**c**) Pocket observed in the previously reported AT1R- TRV023-AT110i1 structure (PDB: 6OS1) is suitable for accommodating the trifluoromethylbenzyl group of Cmpd-23. Notably this pocket for the trifluoromethylbenzyl group is present without Cmpd-23 binding.

(**d**) Actual binding pocket for the trifluoromethylbenzyl group of Cmpd-23 observed in the AT1R-TRV023-Cmpd-23-AT110i1 structure from the current study.

| Allosteric Cooperativity with AngII | | | | | | | | | | | | |
|---|---|---|---|---|---|---|---|---|---|---|---|---|
| **Allosteric Cmpds** | Extended Allosteric Ternary Complex Model | | | | Combined Operational Model Allostery | | | | | | | |
| | Competition Binding | | | | Gq Calcium Response | | | | β-arrestin Recruitment | | | |
| | $K_B$ (μM) | Ki shift (fold) | α | $R^2$ | α | β | $\tau_B$ | $R^2$ | α | β | $\tau_B$ | $R^2$ |
| **Cmpd-23** | 4.2 | 28 | 32.8 | 0.99 | 23.8 | 0.006 | 0 | 0.98 | 17.5 | 0.06 | 0.43 | 0.96 |
| **Cmpd-37** | 12.0 | 7.0 | 8.3 | 0.98 | 2.0 | 2.0 | 0 | 0.95 | 1 | 1 | 0 | 0.95 |

**Extended Data Table 1. Allosteric Cooperativity of the Cmpd-23 and Cmpd-37 with AngII in activity assays.** Allosteric parameters obtained from competition binding data were quantified by extended allosteric ternary complex model[29], while allosteric values from functional assays were quantified by combined operational model of allostery[30-32]. α >1: positive effect on affinity; α =1: neutral cooperativity; α <1: negative effect on affinity. β >1: positive effect on efficacy; β =1: neutral cooperativity; β <1: negative effect on efficacy. $\tau_B$ is the intrinsic efficacy value of allosteric ligands. $\tau_B$ > 0: allosteric agonism.

| | Cmpd-23 (AT1R and AT110i1 only) | Cmpd-37 |
|---|---|---|
| **Data Collection and processing** | | |
| Magnification | 81,000x | 105,000x |
| Voltage | 300 kV | 300 kV |
| Electron exposure | 56.3 e–/Å2 | 60 e–/Å2 |
| Defocus range | -1.0 to -2.0 μm | -1.0 to -2.0 μm |
| Pixel size | 1.1 Å | 1.06 Å |
| Symmetry imposed | C1 | C1 |
| Final particle images | 907,280 | 276,067 |
| Map resolution | 3.0 Å | 3.2 Å |
| FSC threshold | 0.143 | 0.143 |
| **Refinement** | | |
| Initial model used | 6OS1 | 7F6G |
| Map sharpening B factor | 91.3 | 96.0 |
| Model composition: non-hydrogen atoms | 3446 | 8243 |
| Model composition: protein residues | 436 | 1123 |
| Ligands | 3 | 2 |
| **Validation** | | |
| Bond length RMSD | 0.004 | 0.003 |
| Bond angles RMSD | 0.557 | 0.625 |
| MolProbity score | 1.27 | 1.83 |
| Clashscore | 3.67 | 8.14 |
| Poor rotamers (%) | 0.88 | 0.4 |
| **Ramachandran plot** | | |
| Favored (%) | 97.42 | 94.35 |
| Allowed (%) | 2.58 | 5.65 |
| Outliers (%) | 0 | 0 |

**Extended Data Table 2. Cryo-EM data collection and refinement.**
